# Decoding the role of ADAptor2 (ADA2) of HAT complex in autophagy and phospholipid metabolism to maintain ER homeostasis and triterpene regulation

**DOI:** 10.64898/2026.02.09.704976

**Authors:** Alka Rathore, Niveditha N Nawada, Monala Jayaprakash Rao, Manjushri Anbarasu, Ravi Manjithaya, Ashutosh K. Tiwari, CH Ratnasekhar, DK Venkata Rao

**Author notes:** **Corresponding Author Contact details:** Venkata Rao, DK (or), CSIR-Central Institute of Medicinal & Aromatic Plants, Research center, Allalasandra, GKVK (post), Bangalore, Karnataka, INDIA.

## Abstract

In yeast, transcriptional adaptor 2 (ADA2; SAGA complex subunit ADA2), a member of histone acetyltransferase (HAT) complex, regulates transcription through cell signalling, but its precise role in cellular metabolism remains unclear. In this study, genetic loss of *ADA2* (*ada2*Δ) induces squalene (SQ) accumulation, indicating aberrant triterpene metabolism, coupled with endoplasmic reticulum (ER)/nuclear ER (nER) expansion. Lipid analyses of *ada2*Δ revealed elevated phosphatidic acid (PA) and phosphatidylcholine (PC) levels, indicating disrupted phospholipid metabolism. The expanded ER causes basal autophagy elevation, cellular recycling, and nER phagy, suggesting a regulatory role for *ADA2* in autophagy. Downregulation of phosphatidate cytidylyltransferase (*CDS1*) and inositol-3-phosphate synthase (*INO1*), coupled with elevated PA and PC in *ada2*Δ, points to a significant disruption in cytidine-diphosphate–diacylglycerol and phosphatidylinositol pathway. Overexpression of *CDS1* or *INO1*, or the inositol supplementation, in *ada2*Δ restores SQ, basal autophagy and ER phagy. The observed target of rapamycin Ser/Thr kinase complex (TORC1) activity in *ada2*Δ is due to the high PA content. Rapamycin-mediated inhibition of TORC1 reduced SQ, PA and ER expansion while increasing lipid droplets. In contrast, a rapamycin-treated *ada2*Δ*pah1*Δ strain retained high PA, SQ and ER expansion, underscoring the functional role of TORC1–nuclear envelope morphology protein 1 (Nem1)/sporulation-specific protein SPO7 (Spo7)–Pah1 axis. Notably, SQ levels remained unchanged in a rapamycin-treated *ada2*Δ*atg39*Δ strain, suggesting that loss of nER-phagy receptor, Atg39, impairs the effectiveness of TORC1 inhibition. In conclusion, our data unveiled a critical role for Ada2 in maintaining the intricate relationship between lipid and triterpene/sterol metabolism and connecting autophagy and ER homeostasis.

## Introduction

In eukaryotes, transcriptional activation and repression occur continuously in response to internal and external cellular conditions. Transcriptional regulation is influenced by chromatin modifications, particularly histone acetylation at specific lysine residues, which loosens chromatin and promotes gene expression ([1] [2] [3]). In contrast, transcriptional silencing is linked with reduced nucleosomal acetylation, leading to tighter packing of chromatin ([4]). In yeast, the acetylation is mainly mediated by transcriptional regulatory complexes such as Spt-Ada-Gcn5 acetyltransferase (SAGA), Swi/Snf, and Gal80-Gal1/3. The SAGA activates RNA polymerase II-mediated transcription ([5]), and it consists of four major complexes: Histone AcetylTransferase (HAT) complex, deubiquitinating (DUB) complex, TATA-binding protein-associated factor (TAF) complex, and Suppressor of Ty (SPT) complex ([5]). Previous studies have highlighted SAGA’s role in growth, development, and modulation in response to various biological and environmental stress conditions ([6]).

The histone acetyltransferase (HAT) activity is a key regulatory mechanism during transcription and is mainly driven by the HAT complex, which includes four subunits: Gcn5, Ada2, Ngg1, and Sgf29. The HAT activity plays a crucial role in genome-wide transcription, cell proliferation, and DNA damage mechanisms ([7], [8], [9], [10]). Gcn5 serves as the catalytic subunit of the complex, but its function depends on adaptor proteins such as Ada2, Ngg1, and Ada5. Among these, transcriptional ADAptor2 (ADA2) is essential for Gcn5-mediated acetylation, not through direct catalytic activity, but via physical interaction that enables Gcn5 function ([11], [12]). Despite its importance, the specific role of Ada2 in regulating gene transcription and its broader metabolic impact remain poorly understood. In a recent study, it was demonstrated that dGcn5 acts as an autophagy inhibitor by acetylating transcription factor EB in *Drosophila* ([13]). Therefore, HAT activity plays an active role in autophagy regulation through intracellular acetylation. However, it is unclear whether ADAptor protein2 (Ada2) regulates autophagy in a similar manner.

Earlier research found that the HAT complex is important for the activation of SWI/SNF-dependent genes (i.e., *SUC2*, *INO1*, and Ty elements) through its components, such as ADA/GCN5 ([14]), suggesting a possible role for HAT activity in inositol metabolism. In yeast, inositol tightly controls phospholipid biosynthesis, ([15], [16],[17],[18]), and intracellular inositol and its derivatives regulate autophagy ([19]). The HAT complex regulates the expression of *INO1*, a key gene in phospholipid biosynthesis, by acetylating serine10 and lysine14 residues of histone H3 ([20],[21]). These findings suggest an interconnection between autophagy, inositol signalling, and phospholipid metabolism. However, the precise physiological and biochemical role of HAT complex in inositol signalling and phospholipid pathway regulation remains obscure.

The transcriptional ADAptor2 (Ada2) is highly conserved in eukaryotes, including humans and plants ([22],[23]), and its interaction with Gcn5 is necessary for *in vivo* HAT activity ([24],[11], [25]). However, the role of Ada2 in lipid homeostasis and its downstream metabolic effects, such as autophagy and triterpene biosynthesis, remains unclear. Considering the facts mentioned above, we analyzed the single deletion mutants of HAT complex and assessed the levels of the triterpene precursor, squalene (SQ), a key pathway indicator of triterpene/sterol pathway flux ([26]). The *ada2*Δ phenotype exhibits a notably elevated SQ among the HAT complex mutant phenotypes. In yeast, autophagy regulation, phospholipid, and triterpene metabolism are functionally interconnected ([27],[28],[29]). Our confocal microscopy data revealed an expanded ER/nuclear ER morphology in *ada2*Δ, suggesting altered phospholipid synthesis. This observation was supported by RT-qPCR analyses, which showed reduced expression of *CDS1* and *INO1*, key regulators of the phospholipid biosynthesis pathway. The reduced expression of *CDS1* and *INO1* in *ada2*Δ prompted us to investigate total phospholipid content using two dimensional thin layer chromatography (2D-TLC) and comprehensive lipidomic analyses. These analyses revealed significant phospholipid pathway dysregulation in *ada2*Δ, marked by elevated levels of phosphatidic acid (PA) and phosphatidylcholine (PC). It has been shown that an imbalance of phospholipids, particularly PC, impacts basal autophagy in yeast ([30]). This metabolic imbalance in *ada2*Δ was accompanied by increased basal autophagy and endoplasmic reticulum (ER)-phagy, as demonstrated by our investigations using confocal microscopy and immunoblot assays.

In *ada2*Δ, elevated PA levels coincide with sustained activation of the nutrient-sensing serine/threonine-protein kinase TOR1 (TORC1) during the stationary phase. This led us to investigate whether active TORC1 redirects excess PA toward phospholipid synthesis at the expense of triacylglycerol (TAG). During TORC1 inhibition, by rapamycin treatment, *ada2*Δ exhibits high TAG followed by less PA accumulation, suggesting TORC1-Nem1/Spo7 axis-mediated Pah1 activity that preferentially converts PA to diacylglycerol (DAG), thereby promoting TAG and lipid droplet synthesis. This phenomenon was further proved by deleting the Phosphatidic acid phosphohydrolase/phosphatase 1 (*PAH1)* gene in *ada2*Δ. The *ada2*Δ*pah1*Δ strain exhibits elevated squalene (SQ) and PA levels even in the presence of rapamycin. Similarly, upon rapamycin treatment, the elevated SQ levels were unaffected in *ada2*Δ and *ada2*Δ lacking Atg39, a nuclear ER phagy receptor. Therefore, *ADA2* is critical for the multilevel regulation of metabolic consequences associated with altered lipids. In conclusion, these findings demonstrate the functional role of *ADA2* as an autophagy regulator through phospholipid pathway modulation.

## Results

### Loss of Transcriptional ADAptor 2 (ADA2) causes high squalene synthesis and uncontrolled ER biogenesis

To investigate the involvement of HAT complex (**Fig.1A**) in mevalonate-ergosterol (MEV-ERG) pathway regulation, we carried out gas chromatography (GC) to quantify the SQ, an important intermediate metabolite and a key indicator of MEV-ERG pathway activity. Among the HAT complex mutants analyzed (*gcn5*Δ*, ngg1*Δ*, sgf29*Δ*, and ada2*Δ), only *ada2*Δ exhibited a substantial increase in SQ levels, showing nearly an 8-fold accumulation compared to wild-type (WT) (**Fig.1B**). The RT-qPCR data of HAT complex mutant phenotypes showed elevated transcript levels of MEV-ERG pathway genes (**Fig. S1**), but the most pronounced metabolic effect was specific to the *ADA2* deletion. When *ada2*Δ was complemented with functional *ADA2* with inducible GAL1 promoter, a reduction in SQ levels was observed, supporting the physiological role of *ADA2* in regulating the triterpene/sterol metabolism (**Fig.1C**). Given that high SQ levels are often associated with ER expansion, due to the ER-localized nature of MEV–ERG enzymes, we assessed the ER morphology using Sec12-RFP, a marker unique to the nuclear/ER membrane ([31]). Our confocal analyses revealed an uncontrolled ER proliferation in yeast lacking *ADA2*, suggesting its indirect involvement in maintaining ER homeostasis (**Fig.1D)**. Overall, *ADA2* deletion results in an enlarged ER structure with a significant increase in SQ levels. Given the aberrant ER structure in *ada2*Δ, we further investigated whether ER expansion could trigger general autophagy/ER phagy.

**Figure 1.**
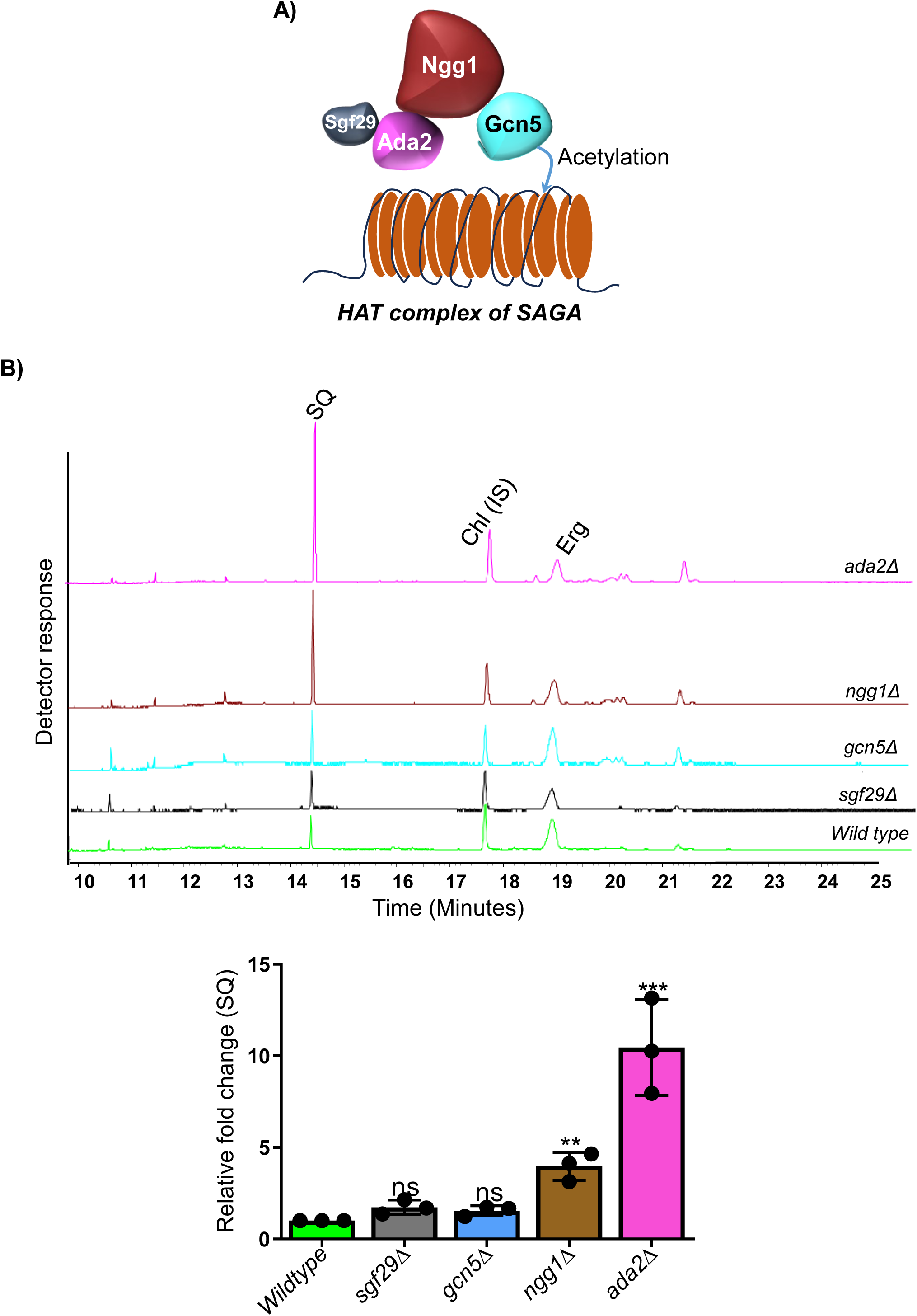
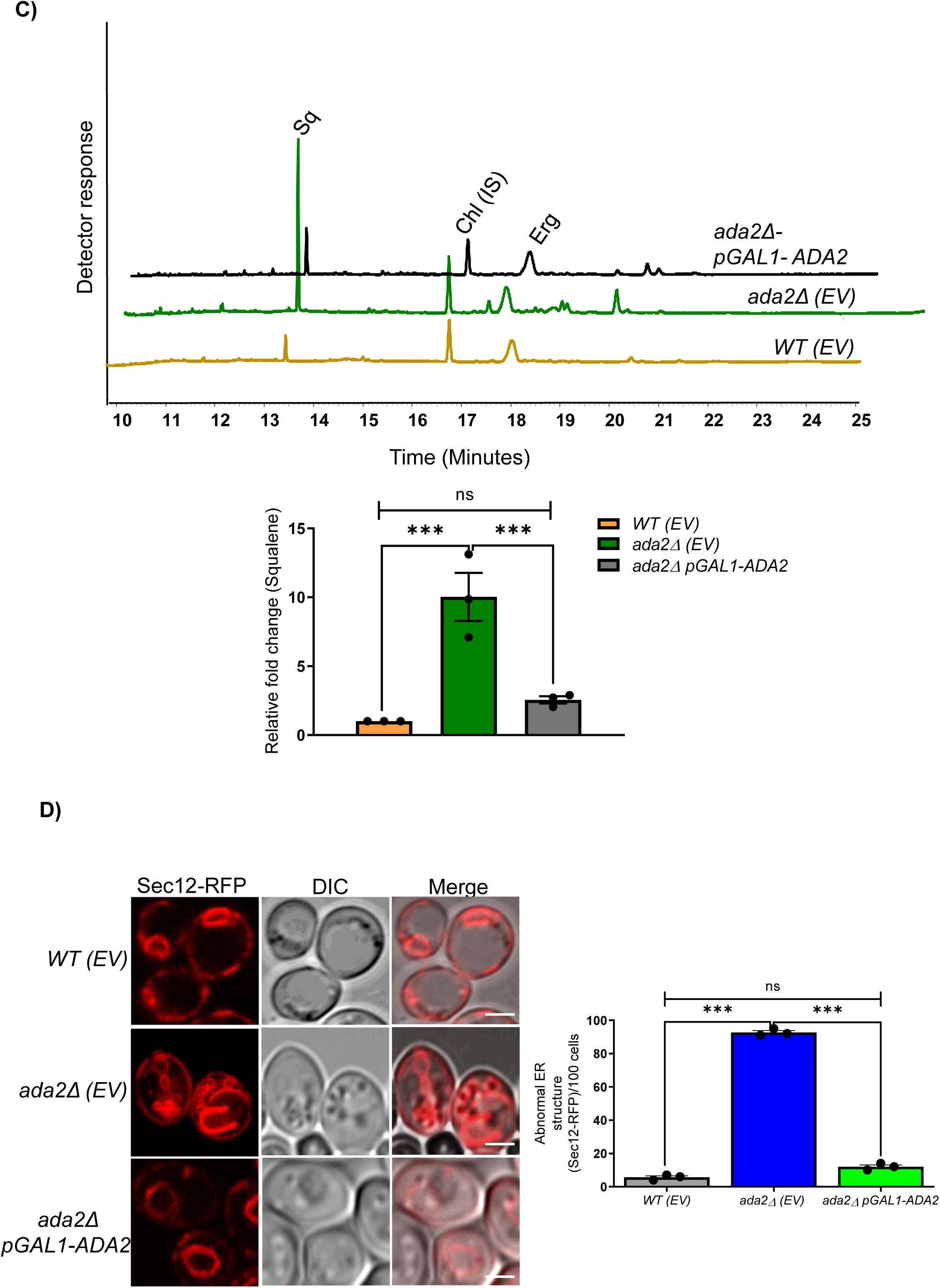
Loss of transcriptional ADAptor (*ada2*Δ) causes aberrant squalene accumulation and endoplasmic reticulum expansion. **A)** Schematic representation of the yeast histone acetyltransferase (HAT) complex composed of four distinct transcriptional protein subunits: Ada2, Ngg1, Gcn5, and Sgf29 (SAGA complex subunits). **B)** Gas chromatography (GC) showing the relative squalene (SQ) levels in single gene deletions of HAT complex (i.e., *sgf29*Δ, *gcn5*Δ, *ngg1*Δ, and *ada2*Δ) and wildtype. Equal optical density (OD_600_) units of stationary phase cells were collected, and the metabolites were extracted for GC analysis as described in the Materials and Methods section. The fold change of SQ was calculated from three independent biological replicates (n = 3). The statistical analysis was carried out using Student’s t-test (Ns, not significant; Ns p> 0.05, **p ≤ 0.01, ***p ≤ 0.001; Error bar ±SD). GC peaks are labelled as follows: SQ, squalene; Chl, Cholesterol (internal standard, IS); Erg, Ergosterol. The bar diagram shows the relative SQ levels in wildtype (green), *sgf29*Δ (grey), *gcn5*Δ (blue), *ngg1*Δ (brown), and *ada2*Δ (pink). **C)** Functional complementation of *ada2*Δ with a galactose-inducible *ADA2* construct (pGAL1-ADA2) shows reduced SQ levels, demonstrating ADA2’s role in MEV-ERG pathway regulation. The *ada2*Δ strain transformed with an empty vector (EV, pESC-LEU2d) acts as a control. The cells were grown to the stationary phase, and equal OD_600_ units of cells were collected for GC analysis to quantify SQ (n = 3; Ns, not significant; Ns p> 0.05, ***p ≤ 0.001 by unpaired Student’s *t* test; error bar ±SD). **D)** The confocal microscopy demonstrates endoplasmic reticulum (ER) morphology in wildtype, *ada2*Δ, and *ada2*Δ pGAL1-ADA2 strains expressing Sec12-RFP (ER marker). The approximately three OD_600_ units of stationary phase cells were used to capture RFP signals using the Zeiss LSM880 Axio Observer confocal microscope. Images were analyzed and processed using ImageJ software (scale bar, 2 µm). The bar diagram shows the abnormal ER structure/100 cells. (Ns, not significant; Ns p> 0.05, ***p ≤ 0.001 by statistical unpaired Student’s *t* test). The standard deviation is indicated by the error bars. SQ, squalene; MEV-ERG, Mevalonate-Ergosterol; RFP, Red Fluorescent Protein.

### Deletion of ADA2 induces general autophagy and ER phagy in yeast

Since previous studies have shown that the ER expansion triggers both ER stress and general autophagy in yeast ([32]), we investigated both general autophagy and ER phagy levels in *ada2*Δ. As expected, increased basal autophagy was observed in *ada2*Δ cells expressing GFP-Atg8 or GFP-Pho8Δ60 compared to the WT. The free-GFP release from GFP-Atg8 was observed in *ada2*Δ (**Fig. 2A**), and confocal imaging further confirmed that the *ada2*Δ possesses high basal autophagy activity, as indicated by vacuoles filled with free GFP. In contrast, the *ada2*Δ pGAL1-*ADA2* strain showed restored GFP-Atg8 puncta similar to WT (**Fig. 2B**). Immunoblot analysis of *ada2*Δ expressing GFP-Pho8Δ60 also confirmed elevated autophagic activity, evidenced by increased free GFP release (**Fig. 2C**).

**Figure 2.**
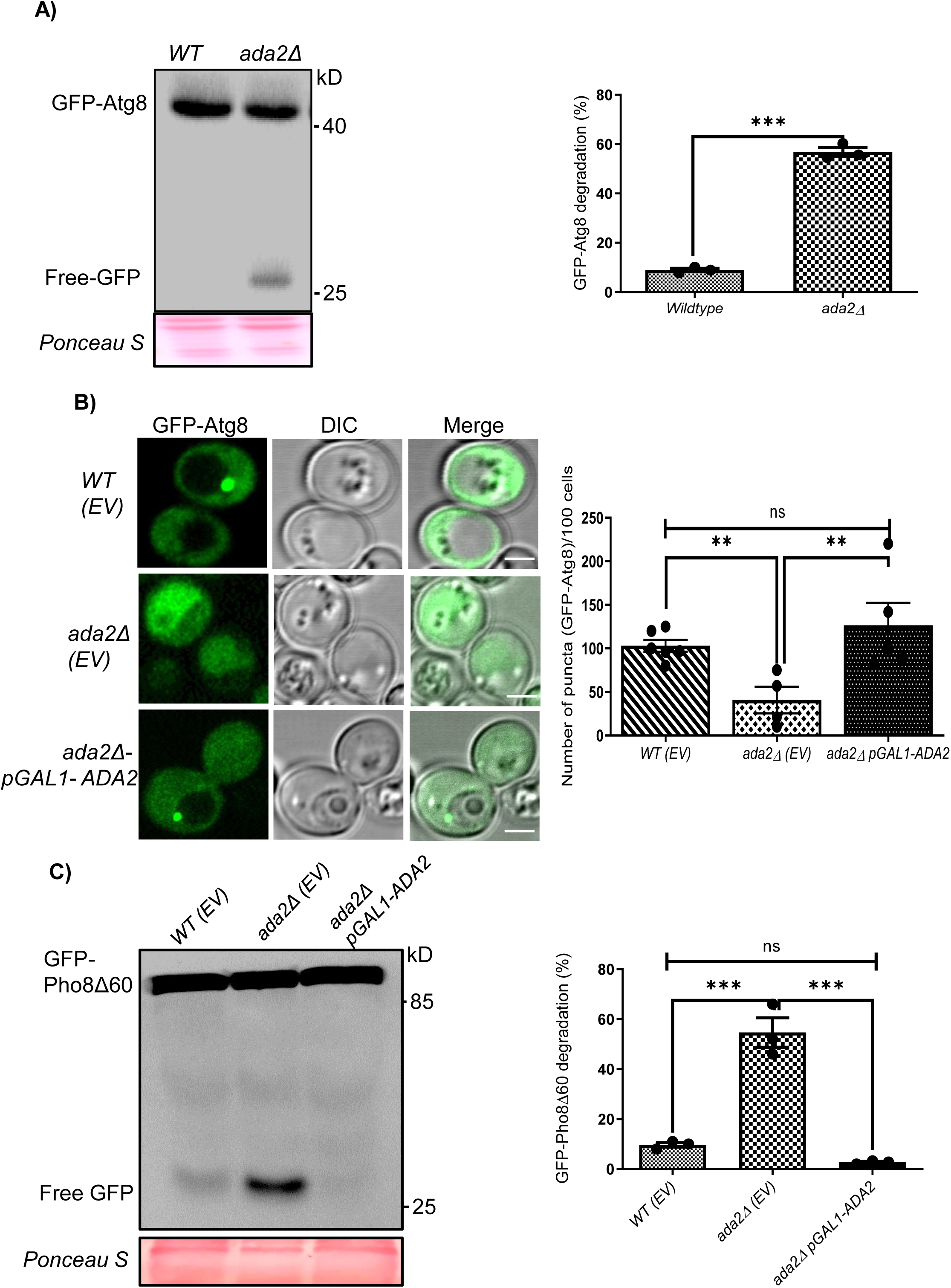
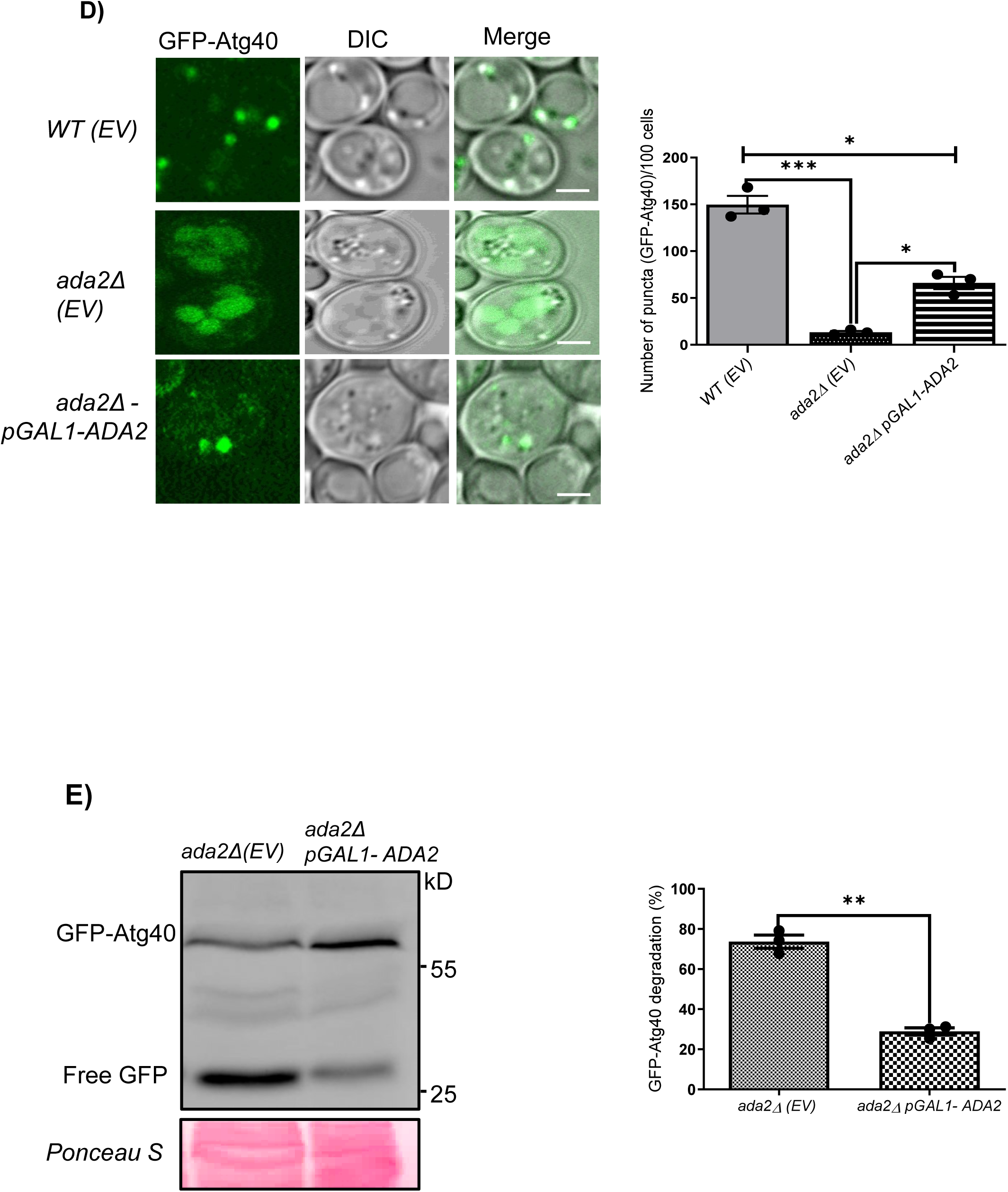
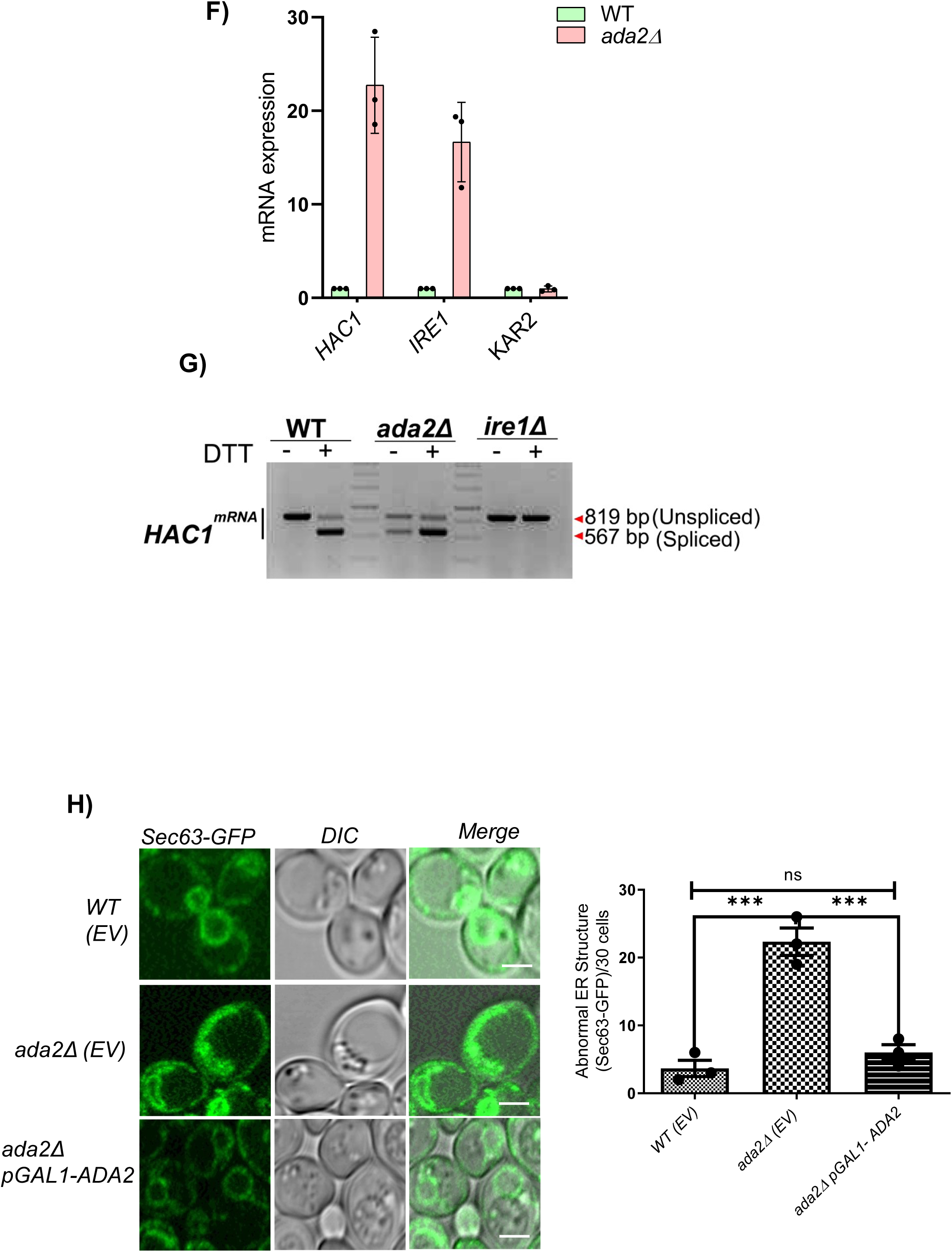
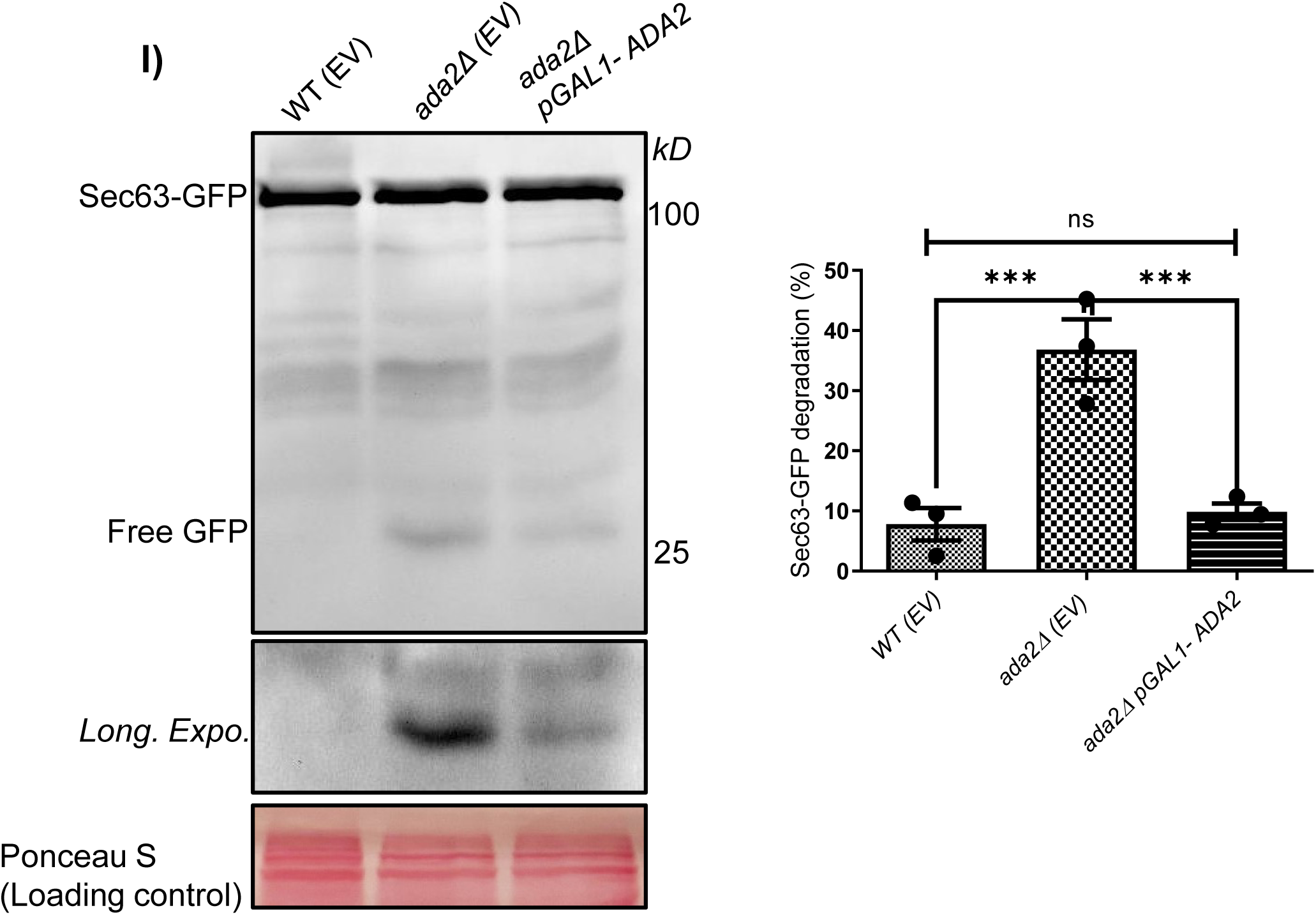
*ada2*Δ exhibits increased general autophagy and ER phagy. **A)** GFP-Atg8 processing assay showing induced basal autophagy in *ada2*Δ. The cells were grown in a selective SD (-His) medium for 24 hrs, and three OD_600_ units of stationary phase cells were harvested for the immunoblotting experiments as described in Materials and Methods. Free GFP release was quantified from three biological replicates using ImageJ software. (n=3; ***p ≤ 0.001 calculated by unpaired Student’s *t* test. Error bar represents ±SD). **B)** Confocal microscopy of GFP-Atg8 transformed in WT(EV), *ada2*Δ(EV), and *ada2*Δ *pGAL1-ADA2* strains (EV, empty vector pESC-LEU2d). GFP signals were imaged using excitation/emission wavelengths of 488/540 nm. Z-stack images (7 stacks) were acquired from three OD_600_ units of stationary-phase cells (Scale bar, 2µm). The GFP-Atg8 puncta were counted using ImageJ cell-counter plugin software (n=3; Unpaired Student’s *t* test P value, Ns, not significant; Ns p> 0.05, **p ≤ 0.01; Error bar represents ±SD). **C)** Immunoblot analysis using GFP-Pho8Δ60 (macro-autophagy marker) shows increased autophagy in *ada2*Δ*(EV)* compared to Wildtype (WT(EV)) and the complemented *ada2*Δ *pGAL1-ADA2* strain (EV, empty vector pESC-LEU2d). Experiments performed in three biological replicates (n=3; Student’s *t* test, Ns, not significant; Ns p> 0.05, ***p ≤ 0.001; Error bar represents ±SD). **D)** To check the ER phagy, GFP-Atg40 was expressed in *ada2*Δ*(EV)*, *ada2*Δ *pGAL1-ADA2,* and WT(EV) strains (EV, empty vector pESC-LEU2d). The confocal imaging of WT(EV) and *ada2*Δ *pGAL1-ADA2* shows punctate GFP-Atg40 structure, while vacuoles with diffused GFP were observed in *ada2*Δ*(EV)*, indicating high ER phagy. Bar graph shows average number of GFP-ATG40 puncta per 100 cells across three biological replicates (n=3); Student’s *t* test performed to check level of significance (P value, ***p ≤ 0.001, *p ≤ 0.05; Error bar denotes ±SD; Scale bar, 2µm). **E)** Immunoblot of GFP-Atg40 processing assay shows free-GFP release in *ada2*Δ*(EV)*, indicating active reticulophagy (EV, empty vector pESC-LEU2d). In contrast, *ada2*Δ *pGAL1-ADA2* exhibits less free GFP release from GFP-Atg40. The Bar graph shows the percent degradation of GFP-Atg40 from three biological replicates (n=3; Student’s *t* test, **p ≤ 0.01; Data shown as ±SD). **F)** RT-qPCR analysis was used to assess the transcript levels of the *HAC1, IRE1*, and *KAR2* genes, which are important indicators of ER stress. *HAC1* and *IRE1* expression levels are considerably higher in the *ada2*Δ strain compared to the wildtype (WT). The entire RT-qPCR experimental procedure is provided in the Materials and Methods section. All experiments were carried out in three biological replicates (n=3; Error bar ±SD). **G)** *HAC1* mRNA splicing was analyzed in wild-type (WT) and *ada2*Δ cells under normal and ER stress conditions. The cDNA from samples were used as a template to amplify the *HAC1* splice variants using primers (Supplementary table S4). The *ada2*Δ mutant displays two *HAC1* spliced mRNA variants, indicative of elevated ER stress relative to the WT. *ire1*Δ acts as control. **H)** Confocal microscopy of Sec63-GFP (a nuclear/ER membrane marker) transformed into *ada2*Δ*(EV)*, *ada2*Δ pGAL1-ADA2, and WT(EV) indicates expanded nER structure in *ada2*Δ*(EV)* strain (EV, empty vector pESC-LEU2d). The bar diagram represents the abnormal nER structures per 30 cells from each of the three biological replicates. (n=3; Student’s *t* test: Ns, not significant; Ns p> 0.05, ***p ≤ 0.001; Error bars indicate ±SD; Scale bar, 2µm). **I)** Immunoblot analysis of Sec63-GFP confirms increased nER degradation in *ada2*Δ*(EV)*, as indicated by higher free GFP release (EV, empty vector pESC-LEU2d). The bar diagram represents the percentage Sec63-GFP degradation in three biological replicates(n=3). Student’s *t* test performed to check the level of significance (P value, Ns, not significant; Ns p> 0.05, ***p ≤ 0.001; Error bars show ±SD).

To assess ER phagy, we examined ER-phagy receptor GFP-Atg40 and ER marker Sec63-GFP in *ada2*Δ. Confocal microscopy revealed vacuolar accumulation of free GFP in *ada2*Δ cells expressing GFP-Atg40, consistent with elevated ER stress. In contrast, punctate GFP-Atg40 structures were present in WT and *ada2*Δ *pGAL1-ADA2* strains, indicating rescued ER-phagy activity (**Fig. 2D)**. Further, GFP-Atg40 processing assay revealed that *ada2*Δ possesses increased basal ER phagy, while this effect was reversed in the complemented strain (**Fig. 2E**). Additionally, transcript level of ER phagy receptor genes, *ATG39* and *ATG40,* were significantly elevated in *ada2*Δ, as shown by RT-qPCR. (**Fig. S2**). Supporting this, our RT-qPCR showed significantly increased expression of ER stress markers *HAC1* and *IRE1* in *ada2*Δ (**Fig. 2F**). To further confirm the activation of endoplasmic reticulum (ER) stress response, which is mediated through the Ire1-dependent pathway, we performed an *HAC1* mRNA splicing assay. In the wild-type strain, only the unspliced *HAC1* mRNA was detected, indicating basal ER homeostasis. In contrast, the *ada2*Δ exhibited the spliced *HAC1* variant, a hallmark of Ire1 activation and ER stress induction (**Fig. 2G**). These results indicate that the ADA2 mutation leads to elevated ER stress, triggering the unfolded protein response (UPR) through the canonical Ire1-Hac1 signalling pathway.

Sec63 protein is an ER-specific marker confined to the nuclear/ER membrane ([33]). Confocal imaging of Sec63-GFP revealed expanded ER structure in *ada2*Δ strain, which was restored to WT levels in the *ada2*Δ *pGAL1-ADA2* strain (**Fig. 2H**). Consistently, immunoblot analysis of Sec63-GFP demonstrated enhanced ER phagy in *ada2*Δ strain (**Fig. 2I**). Overall, the data suggest that *ADA2* deletion leads to enhanced basal autophagy and ER phagy in *S. cerevisiae*, probably due to uncontrolled ER expansion. Next, we have asked whether the expansion of the ER affects the total phospholipids in *ada2*Δ strain.

### *ada2***Δ** strain exhibits elevated PC and PA phospholipids

Phospholipids play a vital role in membrane biogenesis, and our recent studies have linked elevated phospholipid levels with increased squalene accumulation, a key indicator of MEV-ERG pathway activity ([27]). As ER biogenesis is dependent on phospholipid (PL) biosynthesis (**Fig. 3A**), we reasoned that the expanded ER membrane observed in *ada2*Δ may be driven by enhanced PL flux. To test this, we performed two-dimensional thin-layer chromatography (2D-TLC) to analyze total phospholipids. Our results showed a marked accumulation of PC in *ada2*Δ relative to the WT strain (**Fig. 3B).** Complementary lipidomic profiling using LC/MS revealed a significant increase in PC and PA levels in *ada2*Δ (**Fig. 3C&D**), while other phospholipid species levels were found to be decreased **(Fig. S3**). Notably, excess PC flux is known to drive ER expansion ([34]) and has also been linked to increased basal autophagy in yeast ([30]). Next, we employed RT-qPCR to investigate the transcript levels of PL pathway genes to see if *ADA2* deletion causes any changes at the transcriptional level (**Fig. S4**). In contrast to wildtype and other mutants of HAT complex, *ada2*Δ showed significant downregulation of *CDS1* and *INO1* genes. (**Fig.3E, Fig. S5A**), indicating an altered phospholipid metabolism. The observed transcriptional repression of CDS1 and INO1 in the *ada2*Δ led us to investigate whether *ADA2* influences chromatin acetylation at their promoter regions. Chromatin immunoprecipitation (ChIP) assay using an anti-acetyl-histone H3 (Ac-K14) antibody revealed a marked reduction in H3 acetylation enrichment at both the *CDS1* and *INO1* promoter regions in the *ada2*Δ compared with the wild type (**Fig. 3F**). This suggests that the loss of Ada2 function leads to decreased histone acetylation at these loci, correlating with the downregulation of CDS1 and INO1 transcripts. This demonstrates the role of *ADA2* in maintaining H3 acetylation at the *CDS1* and *INO1* promoters for their expression. The GC-MS analysis further demonstrated decreased endogenous myo-inositol levels in *ada2*Δ, resembling the depletion observed in the *ino1*Δ mutant (**Fig. S5B**). The decrease in myo-inositol reflects a repressed *INO1* state in *ada2*Δ, which is caused by hypoacetylation at the *INO1* promoter locus.

**Figure 3.**
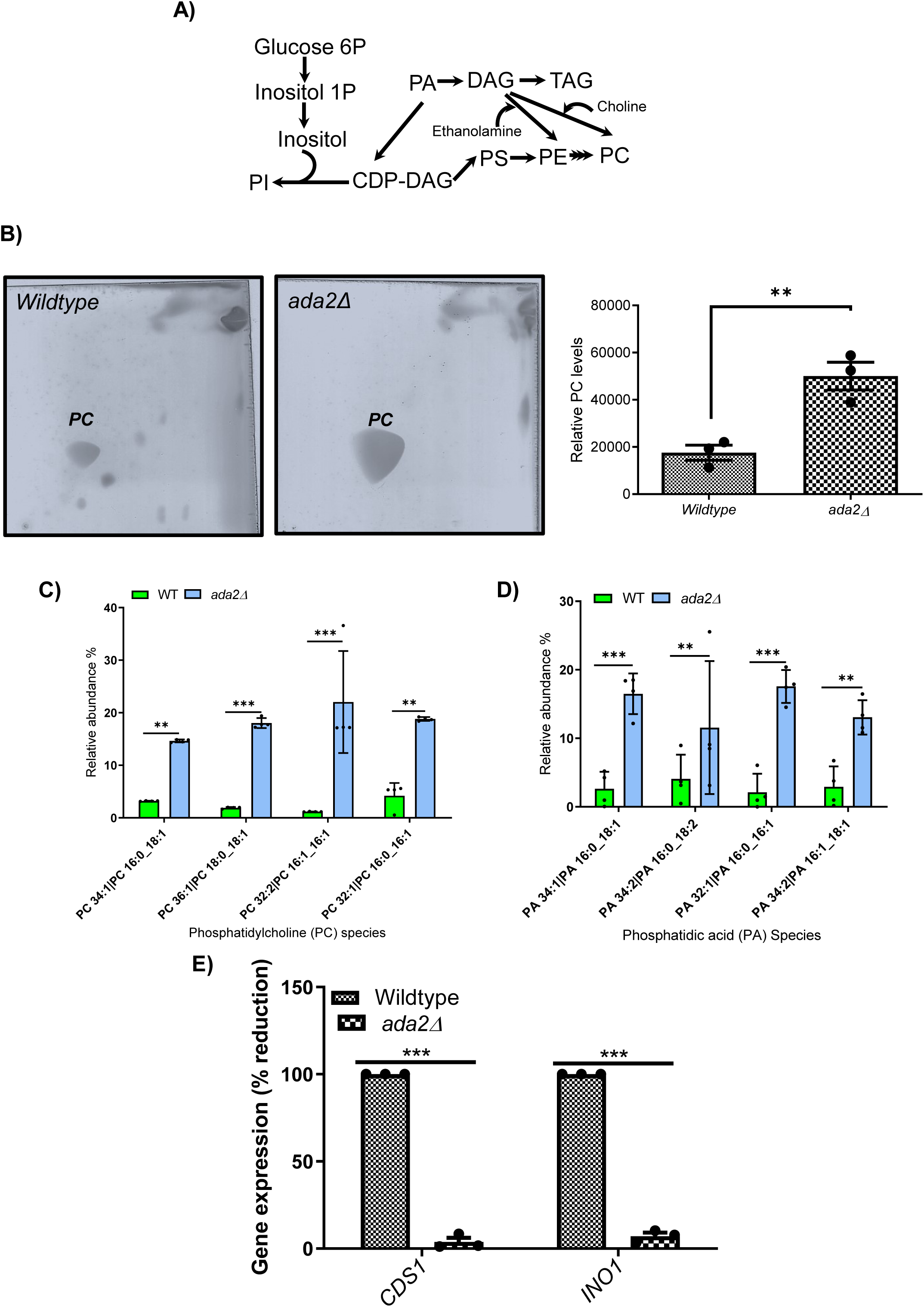
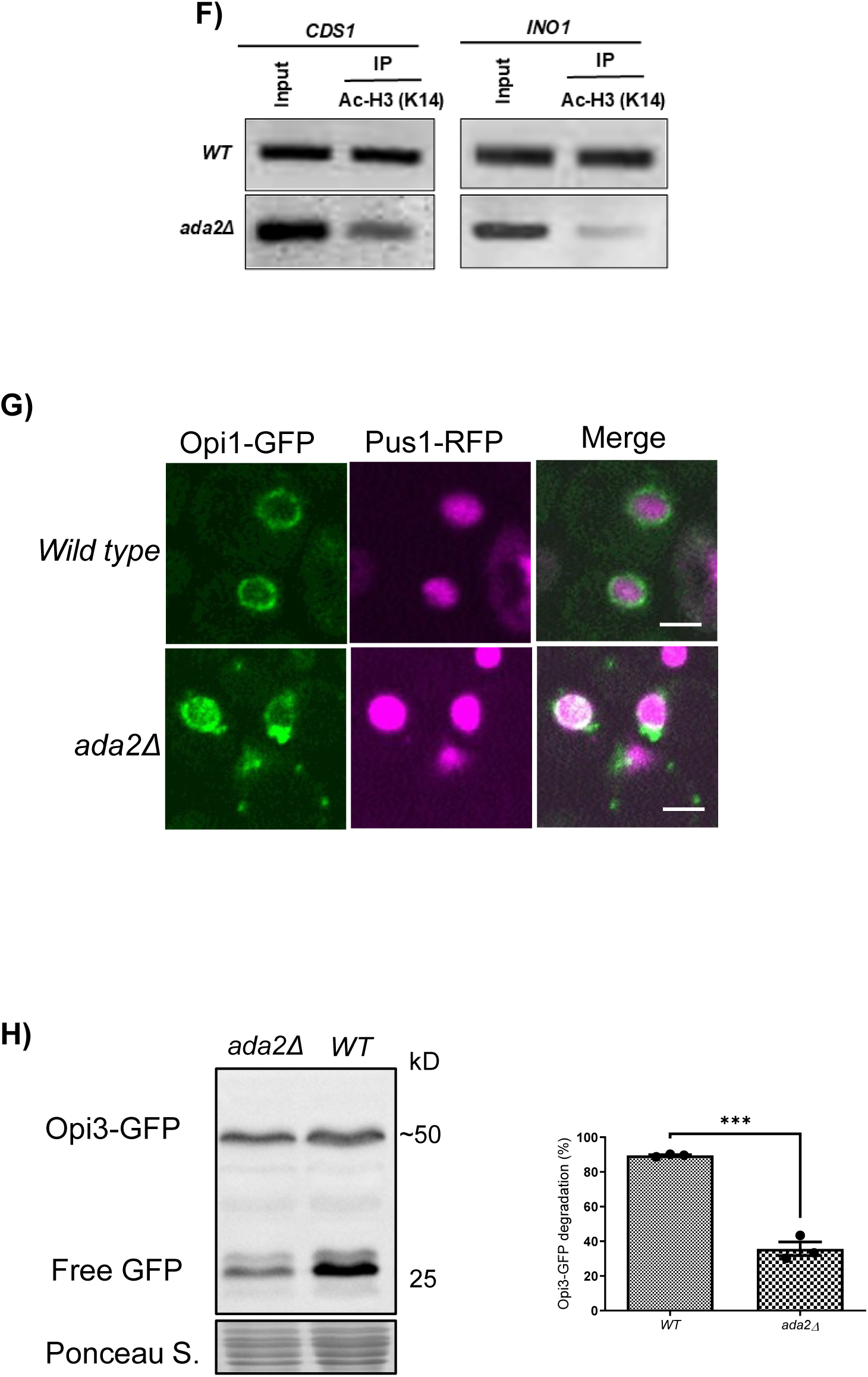
Altered phospholipids resulted in phosphatidylcholine (PC) accumulation in *ada2*Δ. **A)** Schematic diagram of phospholipid biosynthesis in *S. cerevisiae*. **B)** Equal amounts of protein lysate from *WT* and *ada2*Δ were used for total lipid extraction and then separated using two-dimensional thin-layer chromatography, as detailed in the Materials and Methods. The bar diagram shows the relative levels of phosphatidylcholine (PC) spots quantified using ImageJ software. Experiment was performed in three biological replicates (n=3). All statistical analyses were conducted using Student’s *t* test. Level of significance is marked by an asterisk. (P value, **p ≤ 0.01; Error Bars ±SD). **C & D)** Lipidomics analyses of PC and PA quantification using UHPLC - Orbitrap Exploris 240 mass spectrometer system. The equal amounts of protein crude lysate of cell samples were subjected to lipid extraction followed by LC/MS analysis. The detailed description is provided in Material and Method section. (n = 4; P value: **=<0.01; ***=<0.001). **E)** RT-qPCR showing expression of Phosphatidate cytidylyltransferase *1 (CDS1)* and Inositol-3-phosphate synthase *(INO1)* in *WT* and *ada2*Δ. Analysis was done in three biological replicates (n=3; Student’s *t* test, ***p ≤ 0.001; Error bars ±SD). **F)** Loss of ADA2 reduces Histone H3 Acetylation at the *INO1* and *CDS1* promoters. Levels of histone H3 (K14) acetylation at the *INO1* and *CDS1* promoters are significantly reduced in the *ada2*Δ. WT and *ada2*Δ strains were grown under standard conditions, and chromatin immunoprecipitation (ChIP) was performed as described in the Materials and Methods section (n = 3). **G)** Confocal microscopy of Wildtype and *ada2*Δ demonstrates Phosphatidic acid (PA) levels using Opi1-GFP localization marker. RFP-Pus1 was used as a nuclear marker. (GFP, Green fluorescent protein; RFP, Red fluorescent protein). Experiment performed in three biological replicates (n=3, Scale bar, 2µm). **H)** The genomically GFP tagged Opi3 in *ada2*Δ and wildtype were subjected to Opi3-GFP processing assay to check the stability of Opi3. *ada2*Δ exhibited the stability of Opi3 as evidenced by low free GFP as compared to the wildtype. Experiments were performed in three biological replicates (n=3, ***<0.001). Abbreviations: PA, phosphatidic acid; DAG, diacylglycerol; TAG, triacylglycerol; CDP-DAG, cytidine diphosphate diacylglycerol; PS, phosphatidylserine; PC, phosphatidylcholine; PE, phosphatidylethanolamine; PI, phosphatidylinositol.

The *CDS1* gene encodes Phosphatidate cytidyltransferase that catalyzes the conversion of PA to CDP-DAG. This step is crucial for the CDP-DAG pathway as well as the biosynthesis of phosphatidylinositol (PI). The observed downregulation of *CDS1* in *ada2*Δ led us to monitor the PA level by expressing Opi1-GFP. The Opi1 binds to PA and acts as a PA reporter ([35]). The *ada2*Δ expressing Opi1-GFP showed bright punctate structures on the nuclear ER membrane compared to WT (**Fig. 3G**), suggesting high PA accumulation.

In PC accumulation scenario in *ada2*Δ, we performed western blot analyses of the genomically C-terminally GFP-tagged Opi3 to check the Opi3 stability. As Opi3 primarily catalyses the second and third methylation steps in PC biosynthesis, but it can also catalyse the initial methylation at a lower efficiency[36]. Opi3-GFP showed that the Opi3 enzyme remained highly stable in the ada2Δ strain, as indicated by reduced free GFP release comparable to wild-type levels, thereby demonstrating sustained Opi3 enzyme stability in ada2Δ (**Fig. 3H**).

Overall, the RT-qPCR and Opi1-GFP data indicated decreased endogenous Cds1 or Ino1 activity in *ada2*Δ, resulting in abundant PA localized on the nuclear ER membrane. These results indicated that the deletion of *ADA2* affected the transcriptional activation of *CDS1* and *INO1* genes. While, PC accumulation in *ada2*Δ is due to high Opi3 stability. We further asked the question of whether increased Cds1 or Ino1 activity can alleviate the increased basal autophagy in *ada2*Δ strain.

### Increasing Phosphatidate cytidylyltransferase (Cds1) function in ada2***Δ*** augments ER structure, SQ, and autophagy

Further to examine the role of Cds1 activity in regulating ER morphology, SQ, and PA levels, we overexpressed *CDS1* in *ada2*Δ by transforming cells with *pGAL1-CDS1* plasmid. Our RT-qPCR data revealed elevated *INO1* expression when *CDS1* was overexpressed in *ada2*Δ, indicating restoration of inositol synthesis (**Fig. S6**). Notably, the *CDS1* overexpression restored ER morphology and ER stress in *ada2*Δ to levels similar to WT. It was verified by confocal imaging using ER-specific markers such as Sec12-RFP, Sec63-GFP, and ER adapter, GFP-Atg40, in the *ada2*Δ transformed pGAL1-CDS1 strain (**Fig. 4A-C**). We further monitored ER phagy (GFP-Atg40) and general autophagy (GFP-Pho8Δ60 & GFP-Atg8) by performing immunoblotting experiments of *ada2*Δ pGAL1-*CDS1* strain, that showed reduced free GFP release compared with the *ada2*Δ vector control strain (**Fig. 4D-F**). The general autophagy was verified by GFP-Atg8, showing a punctate structure in *CDS1* overexpressing *ada2*Δ (**Fig. 4G**). Thus, *CDS1* overexpression in *ada2*Δ restored both ER phagy and general autophagy. Overall, *CDS1* overexpression data suggest that diverting the excess PA flux into the CDP-DAG pathway restores the ER morphology in *ada2*Δ. Further, it was observed that *ada2*Δ *pGAL1-CDS1* strain displayed a marked reduction in SQ levels, due to restored ER structure and function to its basal level (**Fig. 4H**).

**Figure 4.**
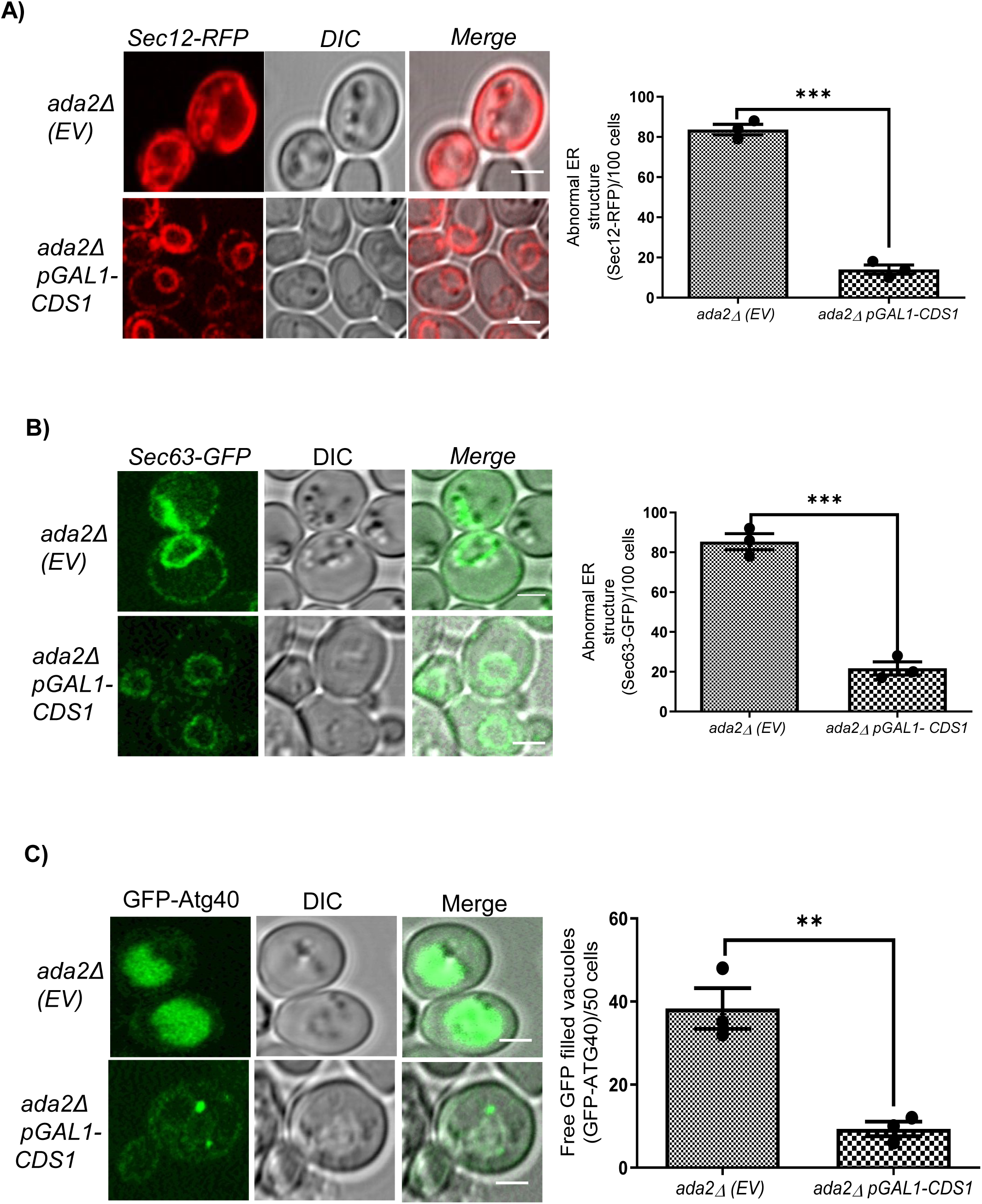
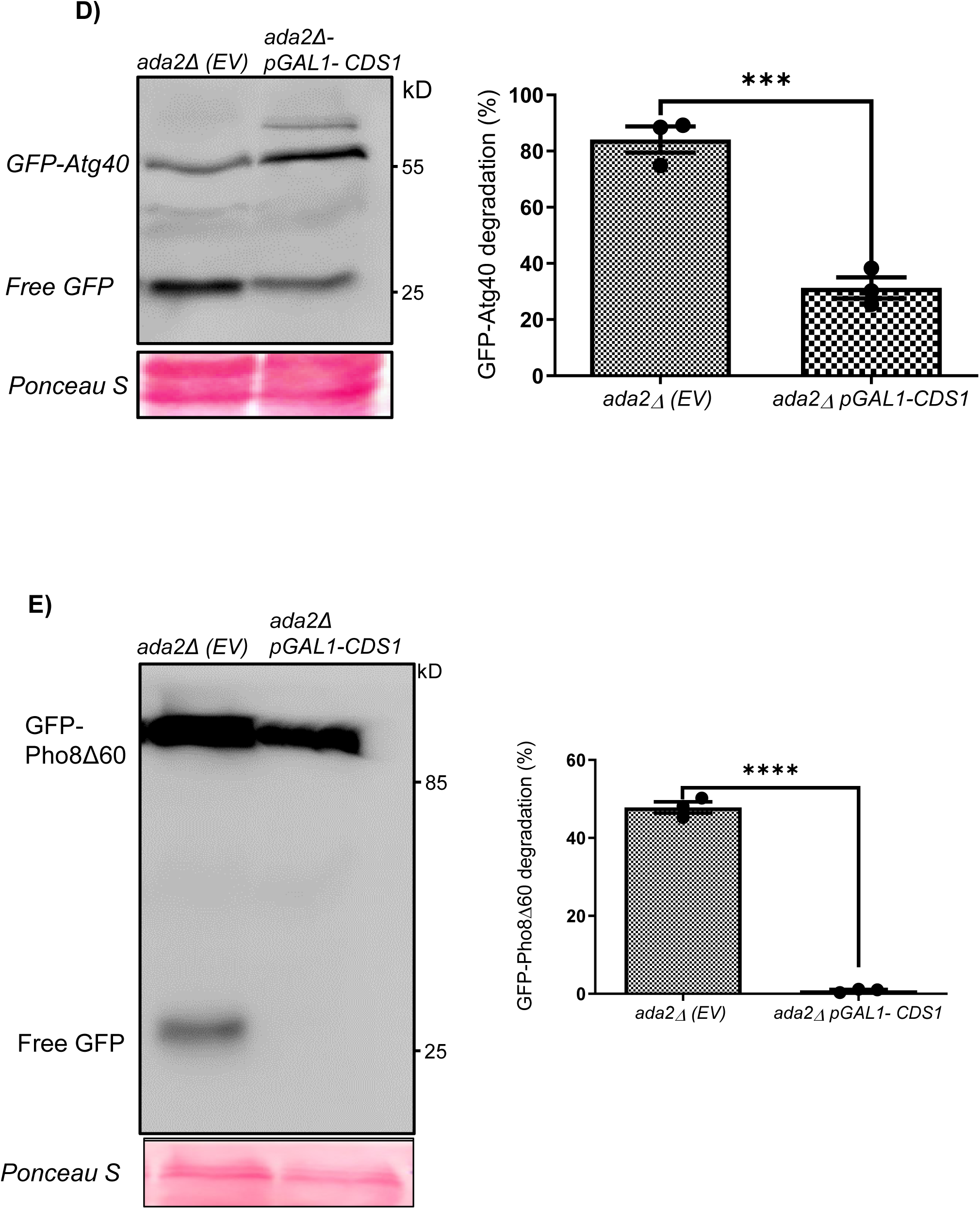
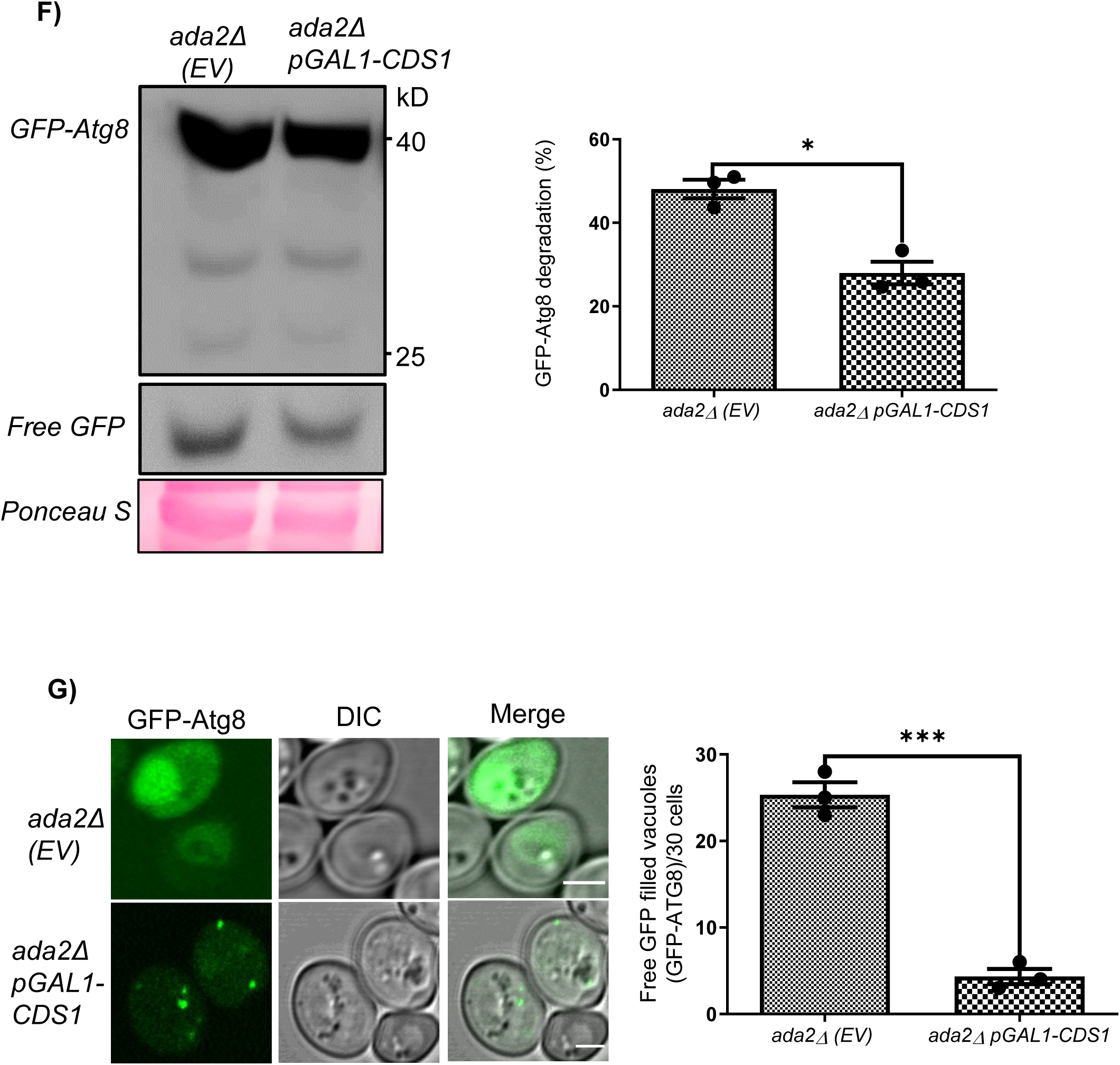
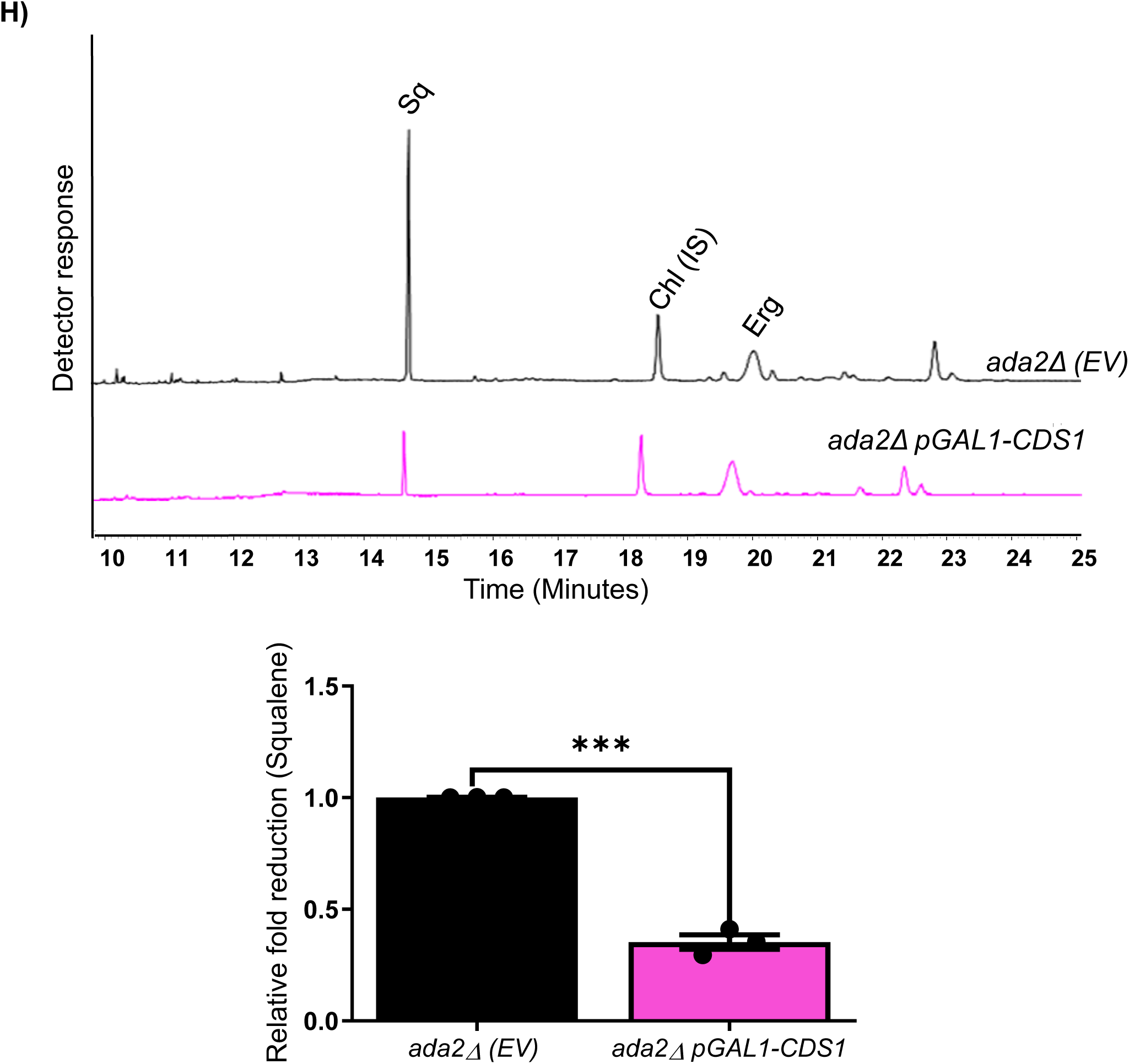
Phosphatidate cytidylyltransferase 1 (*CDS1)* overexpression results in controlled ER biogenesis, reduced autophagy/ER phagy, and Squalene (SQ) levels. **A)** Confocal microscopy shows ER morphology in stationary phase *ada2*Δ*(EV)* and *ada2*Δ *pGAL1*-*CDS1* strains expressing Sec12-RFP (ER marker) (EV, empty vector pESC-LEU2d). The bar diagram represents reduced abnormal ER structure in *ada2*Δ *pGAL1-CDS1* strain compared to *ada2*Δ*(EV)* in three biological replicates (n=3; Student’s *t* test, P value, ***p ≤ 0.001; Error Bars ±SD; Scale bar 2µm). **B)** Confocal images of Sec63-GFP (nuclear ER marker) show reduced ER abnormalities in *ada2*Δ *pGAL1-CDS1* strain compared with *ada2*Δ*(EV)* strain (EV, empty vector pESC-LEU2d). The bar diagram demonstrates reduced abnormal ER in *ada2*Δ pGAL1-CDS1 strain (n=3; Student’s *t* test, P value ***p ≤ 0.001; Error Bars ±SD, Scale bar 2µm). **C)** Confocal imaging of *ada2*Δ expressing *pGAL1-CDS1* strain shows punctate structures of GFP-Atg40, demonstrating ER phagy (EV, empty vector pESC-LEU2d; Scale bar 2µm). The bar diagram quantifies the free-GFP distribution in the vacuole, and the data were obtained from three biological replicates (n=3), each of which had 50 cells (Student’s *t* test, P value, **p ≤ 0.01; Error Bars ±SD). **D)** Immunoblot analysis confirms reduced free-GFP release from GFP-Atg40 in *ada2*Δ *pGAL1-CDS1* strain, indicating decreased ER phagy. The percentage decrease in the GFP-Atg40 degradation is represented as a Bar diagram. Experiment was performed in three biological replicates. (n = 3; Student’s *t* test, P value, ***p ≤ 0.001; Error Bars ±SD). **E)** To monitor general autophagy, the GFP-Pho8Δ60 plasmid was transformed into both *ada2*Δ*(EV)* and *ada2*Δ *pGAL1-CDS1* strains (EV, empty vector pESC-LEU2d). Immunoblot analysis reveals reduced free-GFP release, indicating reduced general autophagy in *ada2*Δ upon pGAL1-CDS1 overexpression. The bar diagram represents the relative GFP-Pho8Δ60 degradation in *ada2*Δ*(EV)* and *ada2*Δ pGAL1-CDS1 strains. The data represent three biological replicates (n=3; Student’s *t* test, P value, ****p≤ 0.0001; Error Bars represent ±SD). **F)** The CDS1-mediated reduction of general autophagy was further confirmed by the GFP-Atg8 processing immunoblot assay, where *ada2*Δ pGAL1-CDS1 strains showed reduced free GFP in comparison to *ada2*Δ*(EV)*. Bar diagram shows percentage of GFP-Atg8 degradation in three biological replicates (n=3, Student’s *t* test, P value, *p ≤ 0.05; Error Bars ±SD). **G)** The confocal microscopy shows a punctate structure of GFP-Atg8 in *ada2*Δ pGAL1-CDS1, in contrast to vacuole-filled free GFP in *ada2*Δ*(EV),* where EV is empty vector (pESC-LEU2d), and the Scale bar is 2µm. The bar diagram represents the vacuole filled with free GFP for 30 cells present in each biological replicates (n=3, Student’s *t* test, P value, ***p ≤ 0.001; Error Bars ±SD). **H)** Gas chromatography analysis demonstrates *CDS1* overexpression in *ada2*Δ leads to reduced squalene accumulation. The relative fold reduction of squalene is depicted as a bar diagram for three biological replicates (n=3; Student’s *t* test, P value, ***p ≤ 0.001; Error Bars ±SD). Abbreviation: Sq, squalene; Chl, cholesterol internal standard; Erg, ergosterol.

### ada2*Δ* with increased Inositol-3-phosphate synthase (Ino1) function leads to the restoration of ER and autophagy

Another important phospholipid pathway gene, *INO1*, that encodes Inositol-3-phosphate synthase, was found to be downregulated in *ada2*Δ. Ino1 is a rate-limiting enzyme in PI pathway and inositol biosynthesis. The *INO1* expression or exogenous inositol supplementation leads to PA depletion in yeast ([37],[18]). As the endogenous PA pool is higher in *ada2*Δ, we were curious whether an increase in inositol levels, either by overexpressing *INO1* or exogenous supplementation of inositol, could deplete the PA pool that can restore the ER structure, autophagy, and triterpene synthesis. To assess this, we examined the ER structure in *ada2*Δ strain expressing *pGAL1-INO1* using the ER marker, Sec12-RFP. Confocal microscopy revealed that the ER morphology was similar to that of the WT. A comparable native ER morphology was observed in *ada2*Δ supplemented with 200 µM of inositol (Hereafter *ada2*Δ^(Exo.Ino)^) **(Fig. 5A-B).** Similarly, using Sec63-GFP, the unexpanded nuclear ER and reduced nucleophagy were observed in both *INO1* overexpressing *ada2*Δ and *ada2*Δ ^(Exo.Ino)^ strains (**Fig. 5C**). This was further supported by GFP processing assay using Sec63-GFP, a nuclear ER marker, where reduced free GFP release was seen in *ada2*Δ pGAL1-*INO1* strain (**Fig. 5D),** while *ada2*Δ showed an increased free GFP, resembling with *ino1*Δ (**Fig. S7A**). These findings suggest that ADA2-mediated inositol levels are crucial for maintaining ER homeostasis in yeast. Further confocal and immunoblotting studies using GFP-Atg40 showed reduced ER phagy in *ada2*Δ expressing pGAL1-INO1 and *ada2*Δ^(Exo.Ino)^ (**Fig. 5E-G & Fig. S7B**). Likewise, the GFP-Pho8Δ*60* and GFP-Atg8 processing assays revealed decreased basal autophagy in *ada2*Δ expressing pGAL1-*INO1* and *ada2*Δ^(Exo.Ino)^ strains (**Fig. 5H-J & Fig. S7C-D**).

**Figure 5.**
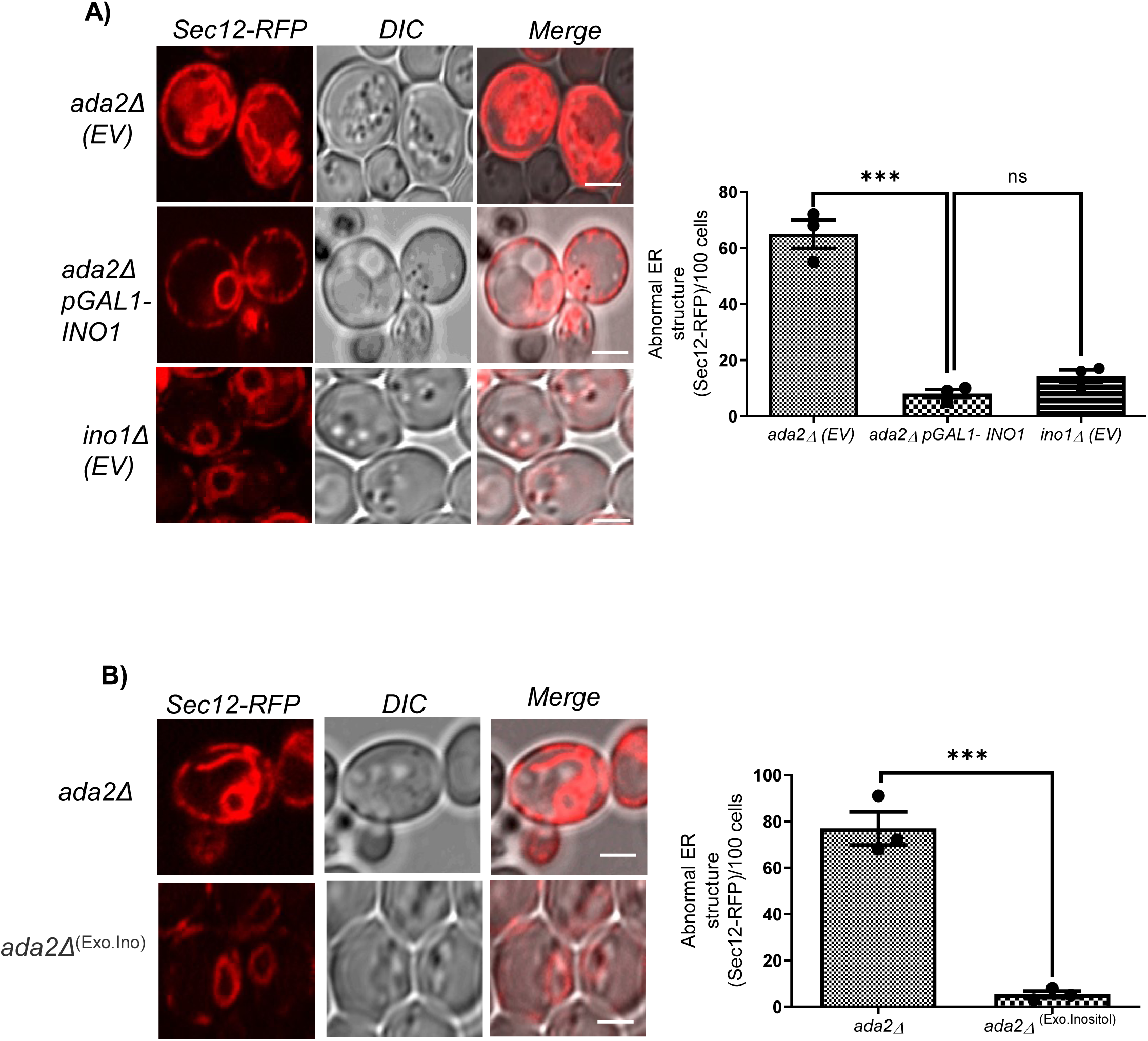
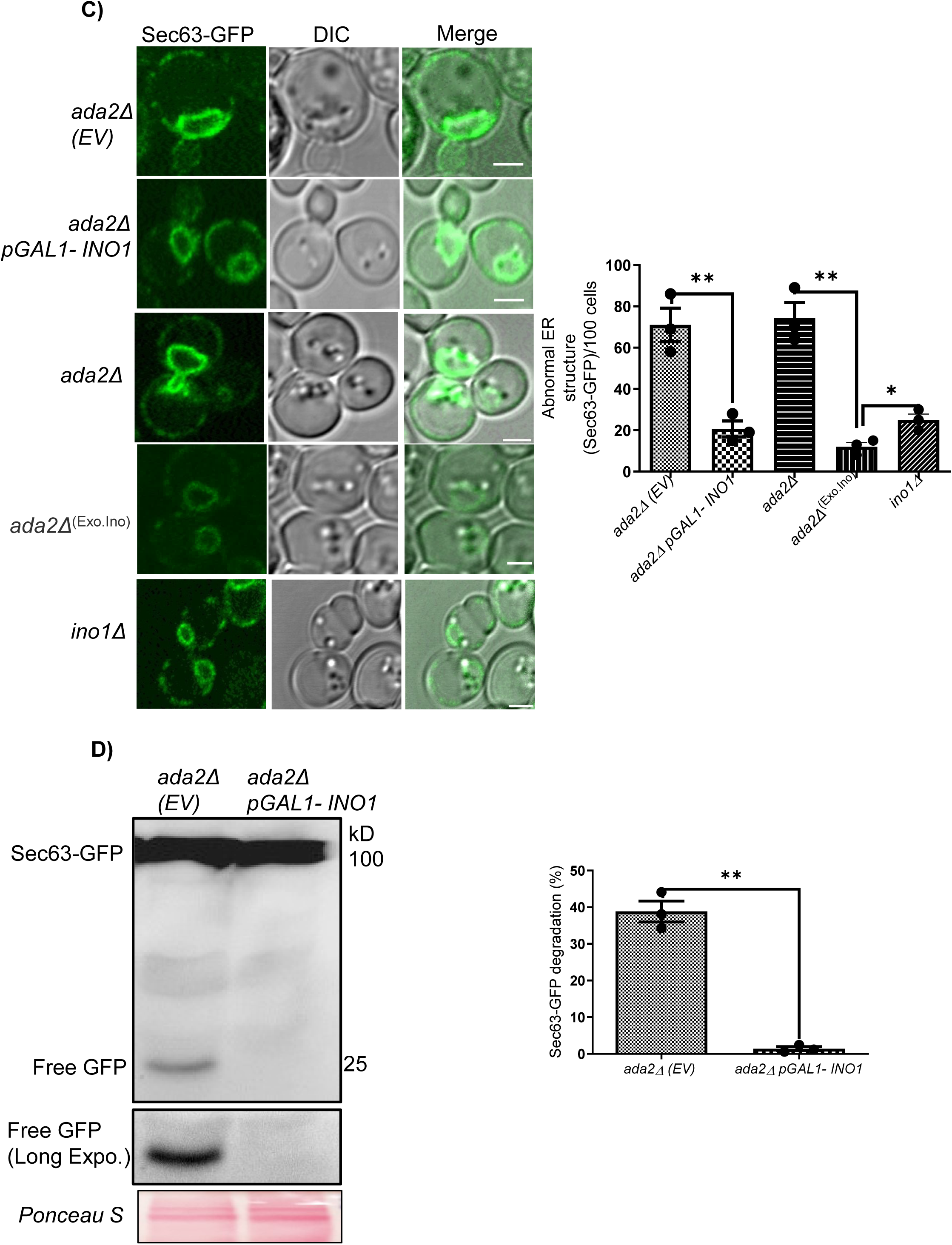
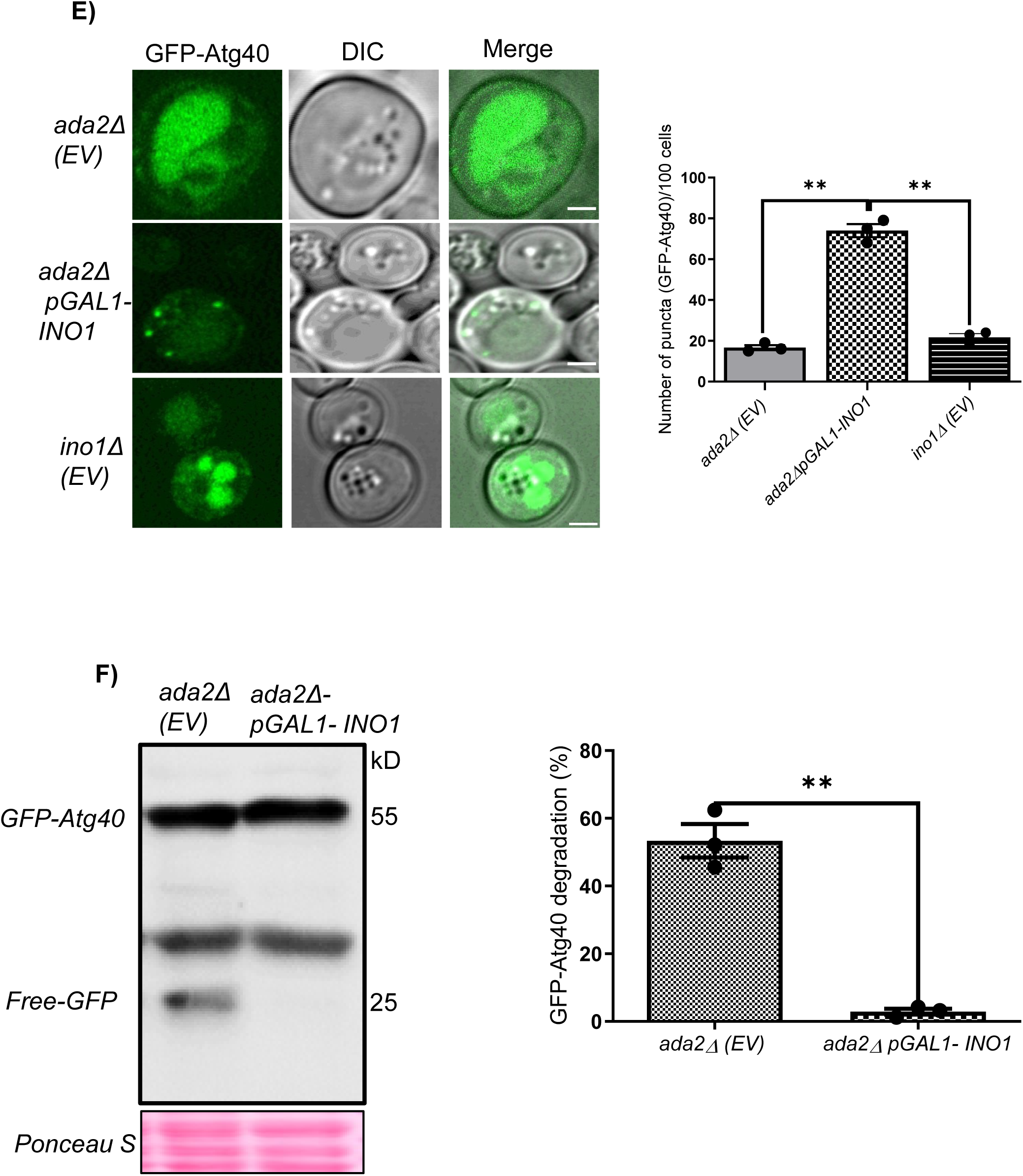
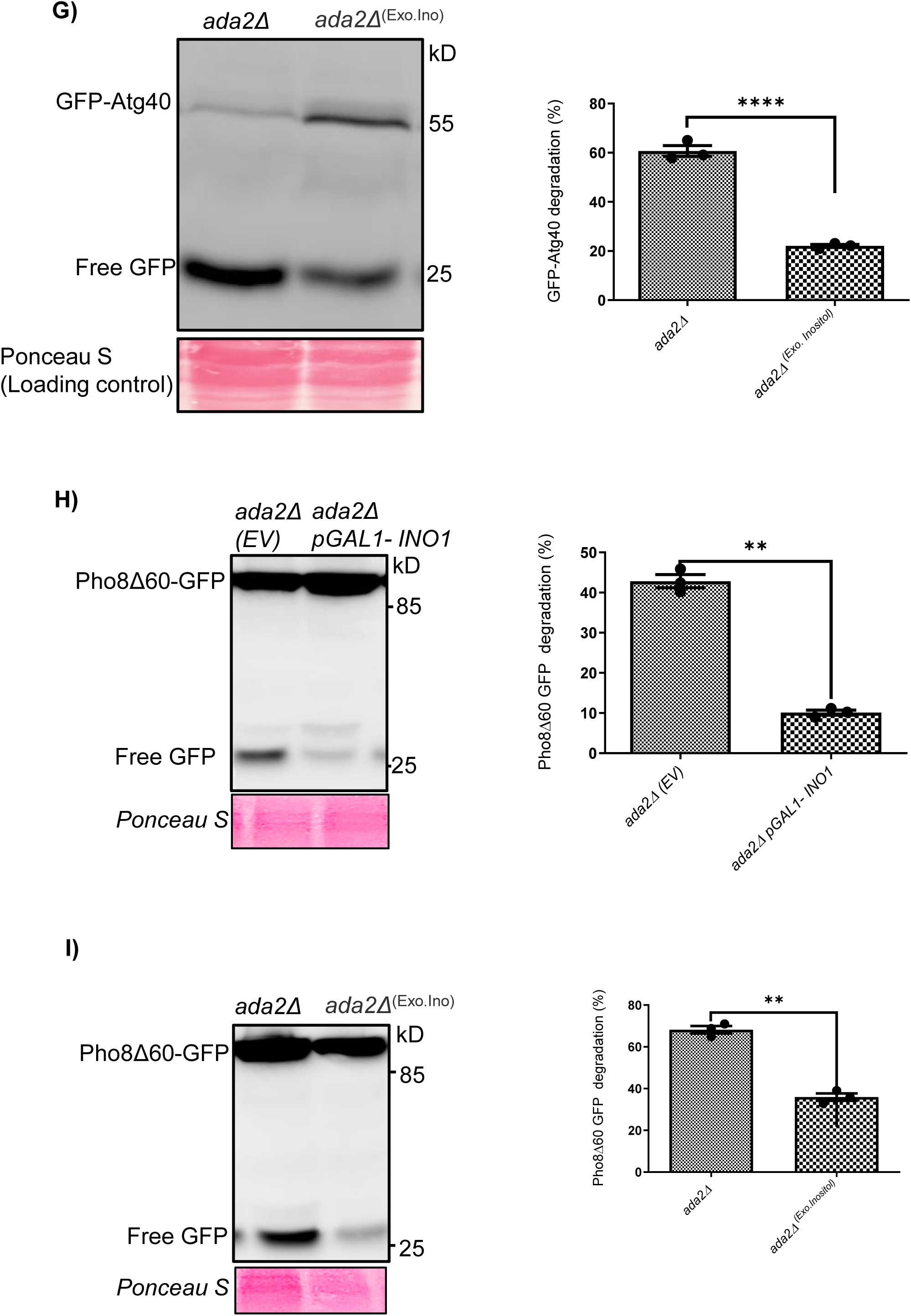
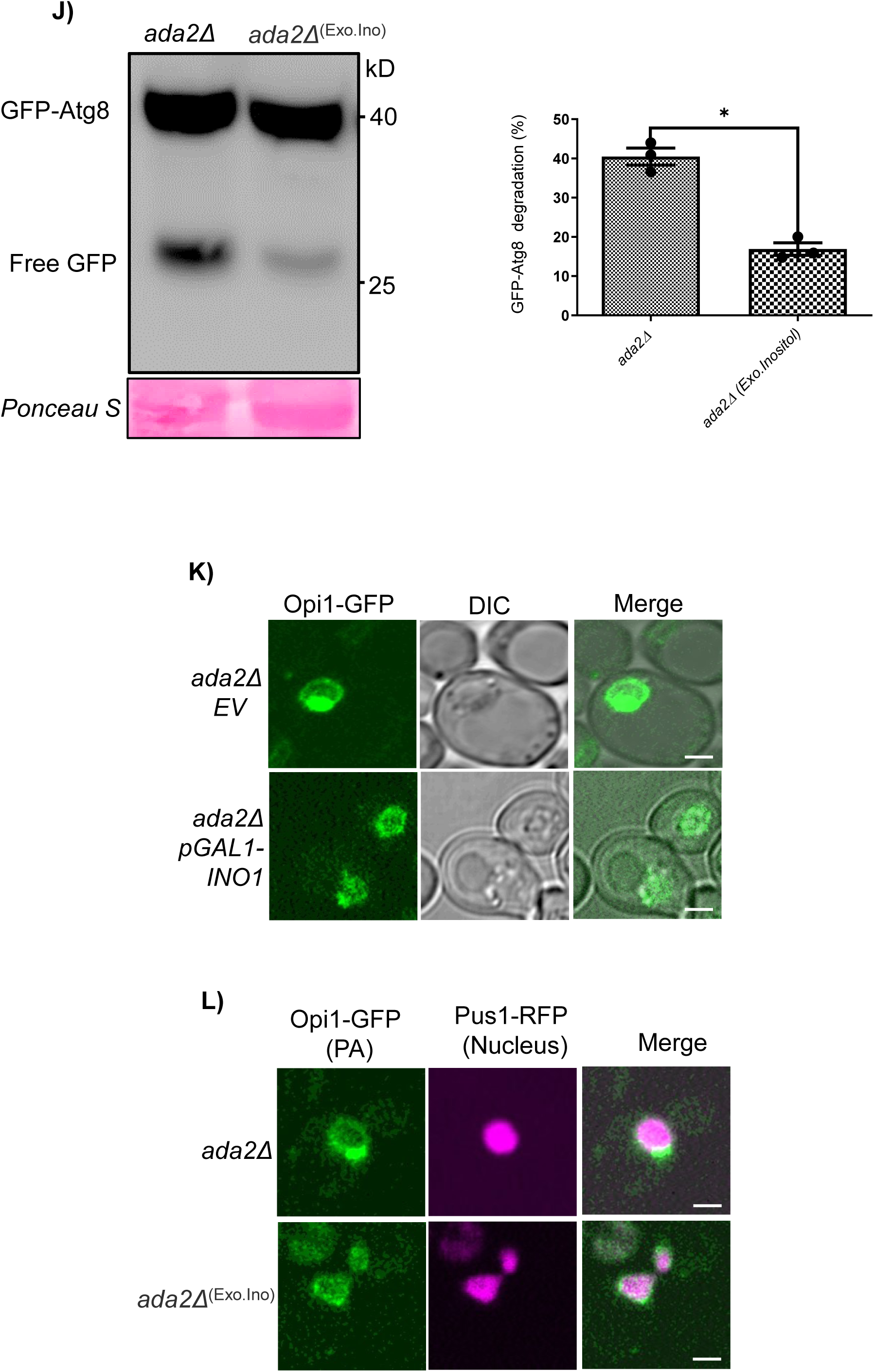
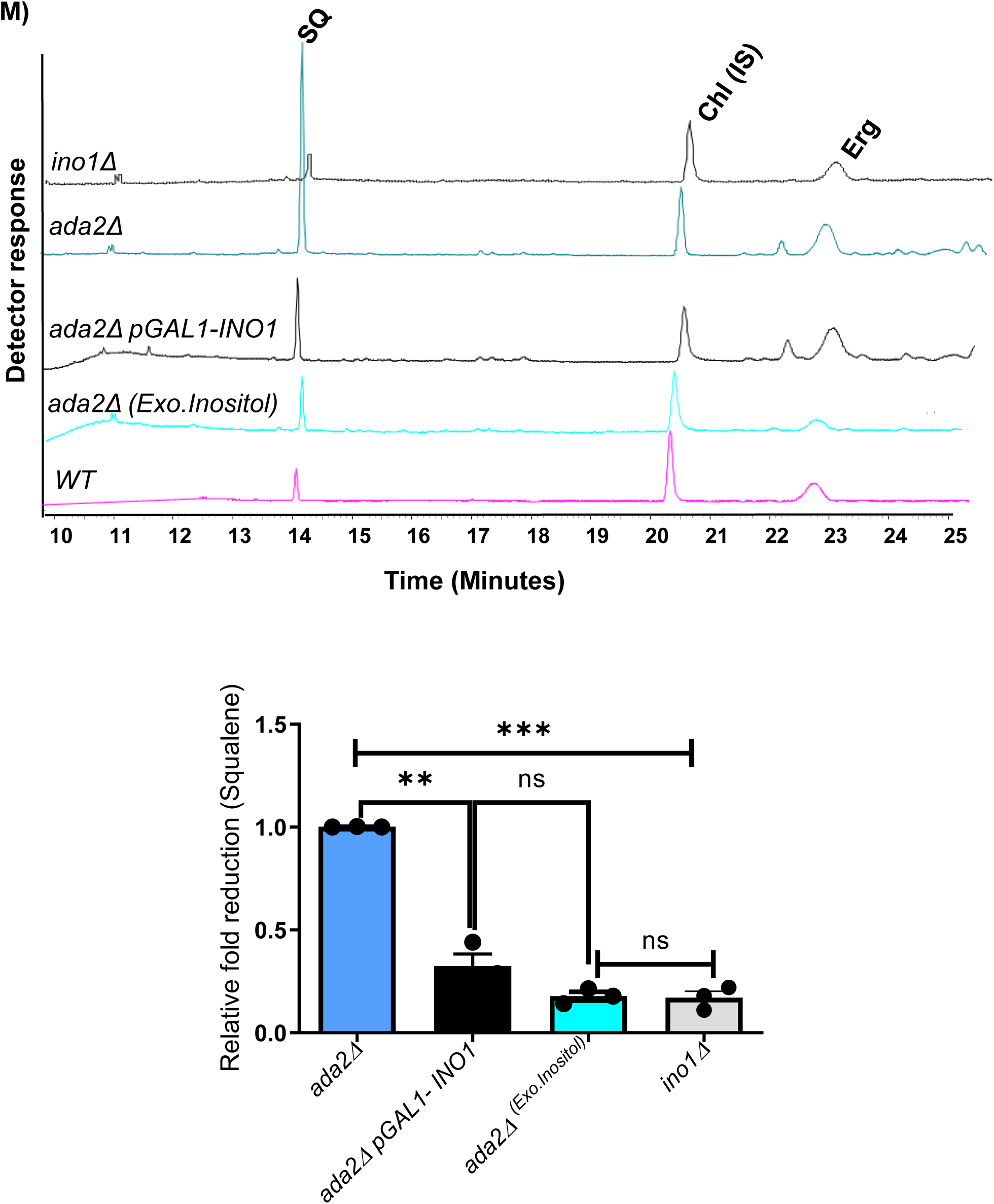
Overexpression of Inositol-3-phosphate synthase *(INO1)* or inositol addition restores endoplasmic reticulum (ER) structure, autophagy, and triterpene synthesis. **A)** Confocal imaging of stationary-phase cells expressing Sec12-RFP in *ada2*Δ*(EV)*, *ada2*Δ *pGAL1-INO1,* and *ino1*Δ*(EV),* where EV is empty vector (MoClo-HIS3 plasmid). The bar diagram represents a decrease in abnormal ER structure in *ada2*Δ *pGAL1-INO1* and *ino1*Δ*(EV)* in comparison to *ada2*Δ*(EV)* in 100 cells per biological replicate (n=3; Student’s *t* test, P value, Ns, not significant; Ns p> 0.05, ***p ≤ 0.001; Error Bars ±SD; Scale bar 2µm). **B)** *ada2*Δ supplemented with 200 µM inositol (hereafter *ada2*Δ ^(Exo.Ino)^) shows a decrease in ER expansion in comparison to *ada2*Δ. The bar graph represents the reduction in abnormal ER structure in *ada2*Δ^(Exo.Ino)^ strains. (n=3; Student’s *t* test, P value, ***p ≤ 0.001; Error Bars ±SD; Scale bar 2µm). **C)** Imaging analyses of *ada2*Δ *pGAL1-INO1*, *ada2*Δ^(Exo.Ino),^ and *ino1*Δ strains with Sec63-GFP (an nER marker protein), show reduced nuclear ER expansion in comparison to *ada2*Δ*(EV)* and *ada2*Δ. The bar diagram demonstrates decreased abnormal nuclear ER structure in the indicated strains (n=3; Student’s *t* test, P value, *p ≤ 0.05, **p ≤ 0.01; Error Bars ±SD; Scale bar 2µm). **D)** Immunoblot analyses show no significant free GFP release from Sec63-GFP in the *ada2*Δ *pGAL1-INO1* strain compared to the *ada2*Δ*(EV),* where EV is empty vector (MoClo-HIS3 plasmid). The Bar diagram shows a reduction in nER phagy in the *ada2*Δ *pGAL1-INO1* compared to *ada2*Δ*(EV)* strain (n=3; Student’s *t* test, P value, **p ≤ 0.01; Error Bars ±SD). **E)** The confocal microscopy analyses show the GFP-Atg40 puncta in *ada2*Δ *pGAL1-INO1*, indicating decreased ER phagy in the stationary phase condition in comparison to *ada2*Δ*(EV)* and *ino1*Δ*(EV).* While the bar diagram represents the number of puncta per 100 cells in three biological replicates (n=3; Student’s *t* test, P value, **p ≤ 0.01; Error Bars ±SD; Scale bar 2µm). **F-G)** Immunoblot analyses of *ada2*Δ *pGAL1-INO1* and *ada2*Δ^(Exo.Ino)^ strains exhibit insignificant free-GFP release from GFP-Atg40, demonstrating reduced ER phagy in comparison to *ada2*Δ*(EV)* and *ada2*Δ respectively (EV, empty vector MoClo-HIS3 plasmid). The bar diagram indicates the percent degradation of GFP-Atg40 (n=3; Student’s *t* test, P value, **p ≤ 0.01, ****p≤ 0.0001; Error Bars represent ±SD). **H-I)** Immunoblot analyses for autophagy using GFP-Pho8Δ60 revealed that the release of free GFP from GFP-Pho8Δ60 is reduced in the *ada2*Δ *pGAL1-INO1* strain and similarly in the case of *ada2*Δ^(Exo.Ino),^ indicating a reduction in macro autophagy (n=3; Student’s *t* test, P value, **p ≤ 0.01, Error Bars ±SD). **J)** Immunoblot analyses show a decrease in free-GFP release from GFP-Atg8 in exogenous inositol addition to *ada2*Δ. The Bar diagram represents the decrease in GFP-Atg8 degradation in *ada2*Δ^(Exo.Ino)^. The experiment was performed in three biological replicates (n=3; Student’s *t* test, P value, *p ≤ 0.05; Error Bars ±SD). **K-L)** Confocal images reveal depletion of Phosphatidic acid (PA) (nuclear localized Opi1-GFP) in *ada2*Δ *pGAL1-INO1* and in *ada2*Δ^(Exo.Ino)^ stationary phase cells in comparison to *ada2*Δ*(EV),* where EV is empty vector (MoClo-HIS3 plasmid) and *ada2*Δ strain, respectively. Analysis was done in three biological replicates (n=3, Scale bar 2µm, Z stack=7) where the Pus1-RFP plasmid was used as a nuclear localization marker. **M)** The activity of the Mevalonate-Ergosterol (MEV-ERG) pathway is monitored by quantifying the squalene (SQ) in *ada2*Δ pGAL1-*INO1* and *ada2*Δ ^(Exo.Ino)^ strains. Reduction in SQ indicates decreased MEV-ERG pathway activity. The bar diagram represents the differential fold change of SQ. (n=3; Student’s *t* test, P value Ns, not significant; Ns p>0.05, **p ≤ 0.01, ***p ≤ 0.001; Error Bars ±SD). (Chl, Cholesterol internal standard; Erg, Ergosterol).

We monitored PA accumulation using the Opi1-GFP as a PA monitoring probe ([35]). The bright punctate structures were observed on the nuclear membrane in *ada2*Δ, while Opi1-GFP was diffused within the nucleus of *ada2*Δ expressing *pGAL1-INO1* and in *ada2*Δ ^(Exo.Ino)^ strains, confirming PA depletion (**Fig. 5K-L**). Consistent with this, gas chromatography revealed that squalene (SQ) levels were significantly reduced in *ada2*Δ *pGAL1-INO1* and *ada2*Δ ^(Exo.Ino)^ strains (**Fig.5M**). High PA flux can drive excess PC formation when *INO1* expression is severely impaired ([38],[39]), as seen in *ada2*Δ cells. The RT-qPCR data revealed the restoration of CDP-DAG pathway genes expression in *ada2*Δ *pGAL1-INO1* and *ada2*Δ ^(Exo.Ino)^ strains (**Fig. S7E, Fig. S8**). Collectively, these data suggest that PA levels are tightly linked to the availability of inositol, indirectly influencing ER biogenesis and autophagy.

### ER expansion and SQ levels in ada2***Δ*** are determined by PA-TORC1-mediated regulation

PA is a master regulator of TORC1; thus, its abundance can lead to continuous TORC1 activity. To assess TORC1 activity in *ada2*Δ, we performed the classical Sch9 NTCB cleavage assay ([40]), which revealed high TORC1 activity in *ada2*Δ stationary phase cells compared to WT (**Fig. 6A**). This was further supported by elevated transcript levels of selected Ribosomal Protein of the Large subunit (RPL) genes consistent with enhanced TORC1 signalling in *ada2*Δ (**Fig. 6B**). The activated TORC1 phosphorylates Nem1-Spo7 phosphatase either by activating unknown protein phosphatase or inhibiting uncharacterized protein kinase, rendering it inactive ([41]). As a result, phosphorylated Nem1/Spo7 is unable to dephosphorylate Pah1, rendering it functionally inactive ([42]). The inactive Pah1 is unable to convert PA into diacylglycerol (DAG), which blocks the downstream synthesis of triacylglycerol (TAG). Lipidomic findings showed reduced TAG levels in *ada2*Δ during the stationary phase (**Fig. S9**), indicating strong TORC1 activity due to high PA content **(Fig. 6A & Fig. 3D)**. To further investigate the involvement of TORC1-mediated PA mobilization in *ada2*Δ, TORC1 activity was inhibited by rapamycin, a competitive inhibitor of TORC1. Upon rapamycin treatment, *ada2*Δ accumulated large numbers of lipid droplets, indicative of restored Pah1 activity leading to increased TAG synthesis (**Fig. 6C, Fig. S9**). This suggests that when TORC1 is inhibited by rapamycin, the active Nem1-Spo7 complex perhaps dephosphorylates the Pah1 ([41]). The activated Pah1 then channels PA into TAG synthesis ([43]). Notably, rapamycin treatment also led to a significant reduction in SQ levels (**Fig. 6D)**, which is likely due to rapid PA mobilization to TAG, resulting in ER degradation (**Fig. S10**). Our lipidomics data showed that rapamycin-treated *ada2*Δ had higher TAG levels, indicating that PA is primarily mobilized to TAG biosynthesis due to inactive TORC1 (**Fig. S9**). The decrease in PA levels was further supported by Opi1-GFP nuclear localization in rapamycin-treated *ada2*Δ cells (**Fig. 6E**). Overall, it is proposed that TORC1 is active in *ada2*Δ, due to the high PA content, which inactivates the Nem/Spo7 phosphatase complex via TORC1-mediated phosphorylation, whereas in rapamycin-treated *ada2*Δ, TORC1 is inactivated, causing the Nem/Spo7 phosphatase complex to become active, eventually activating Pah1 activity via Nem/Spo7-mediated dephosphorylation (**Fig. 6F**). The Pah1 phosphatase activity dephosphorylates the PA to DAG that largely drives to TAG synthesis ([44]).

**Figure 6.**
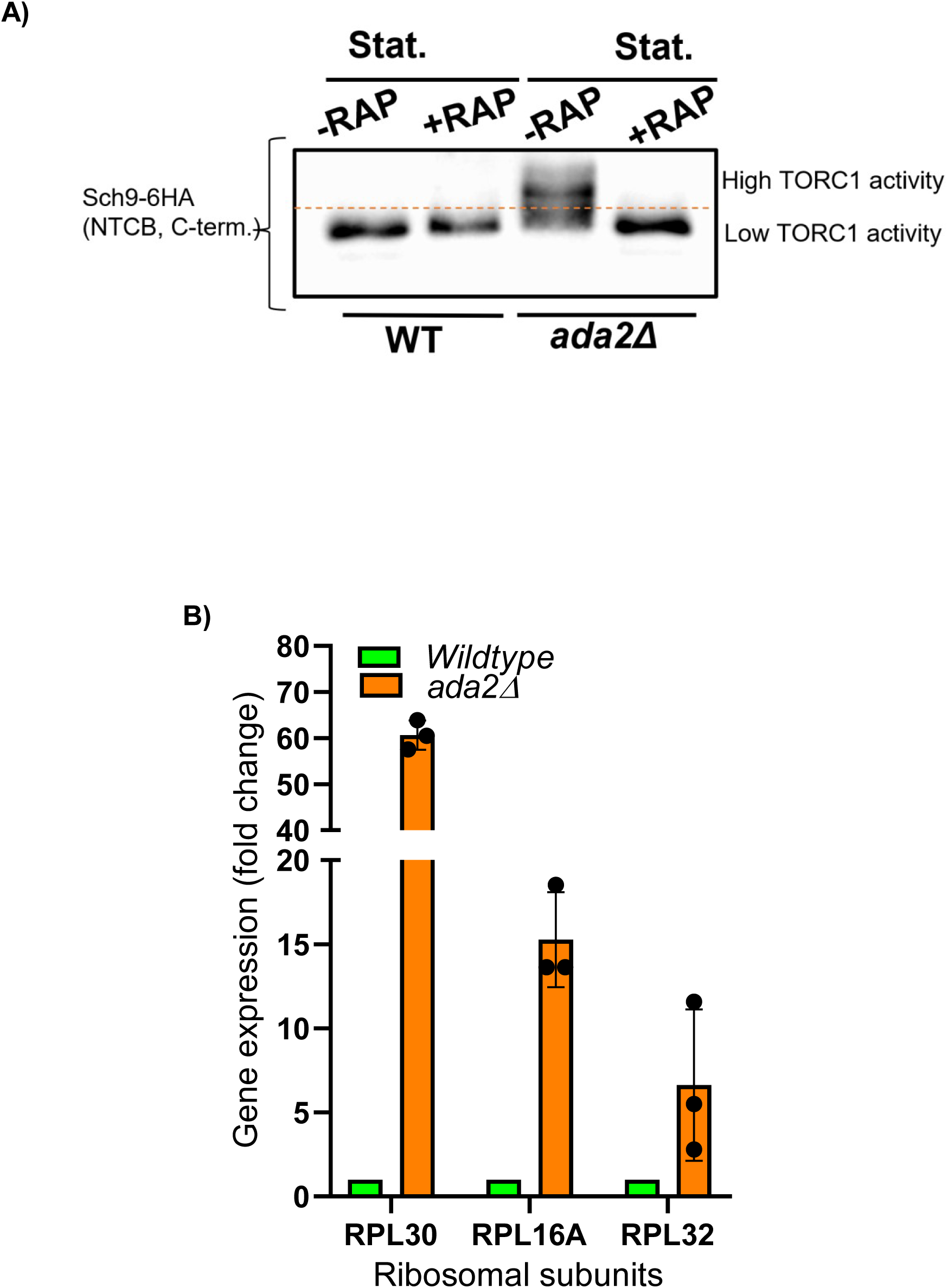
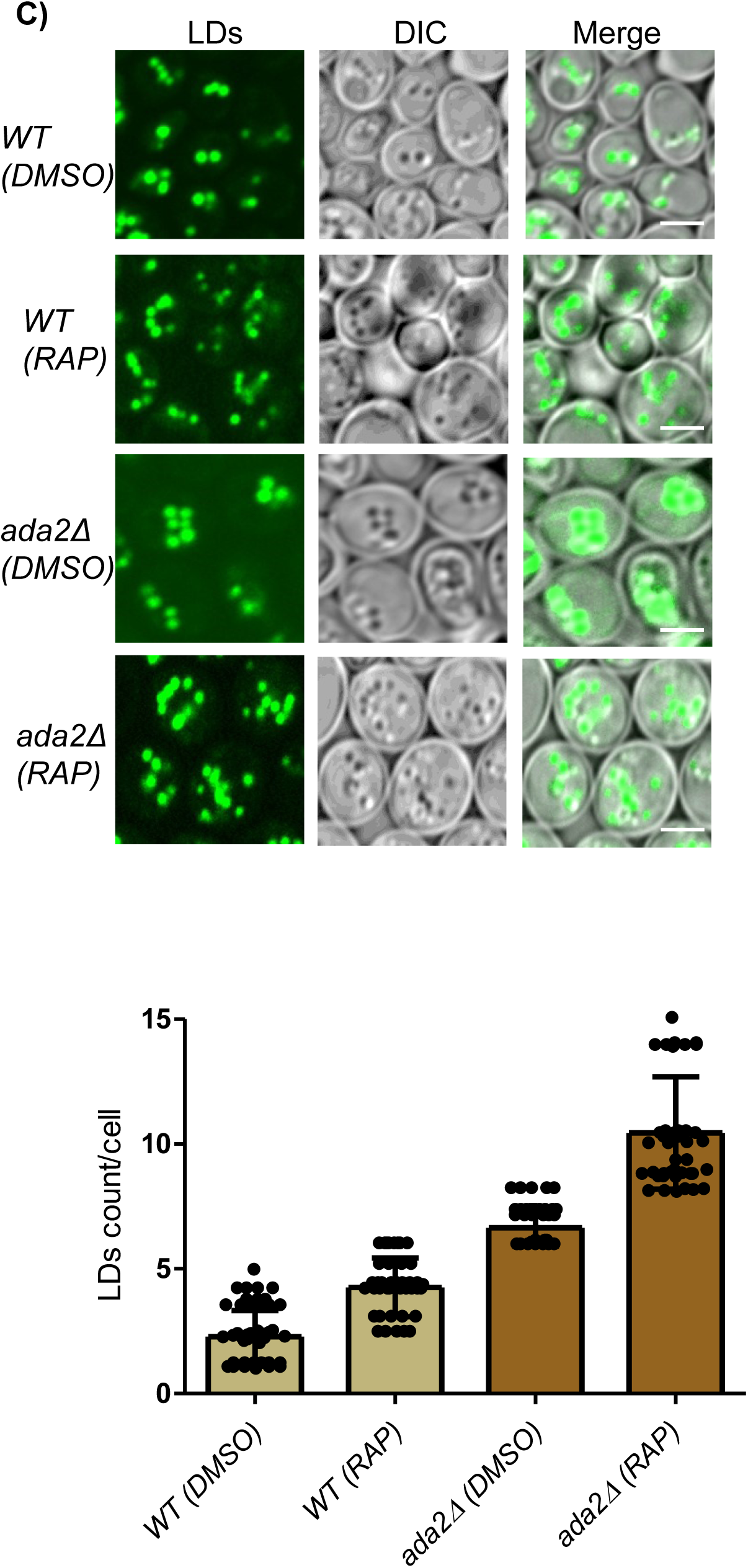
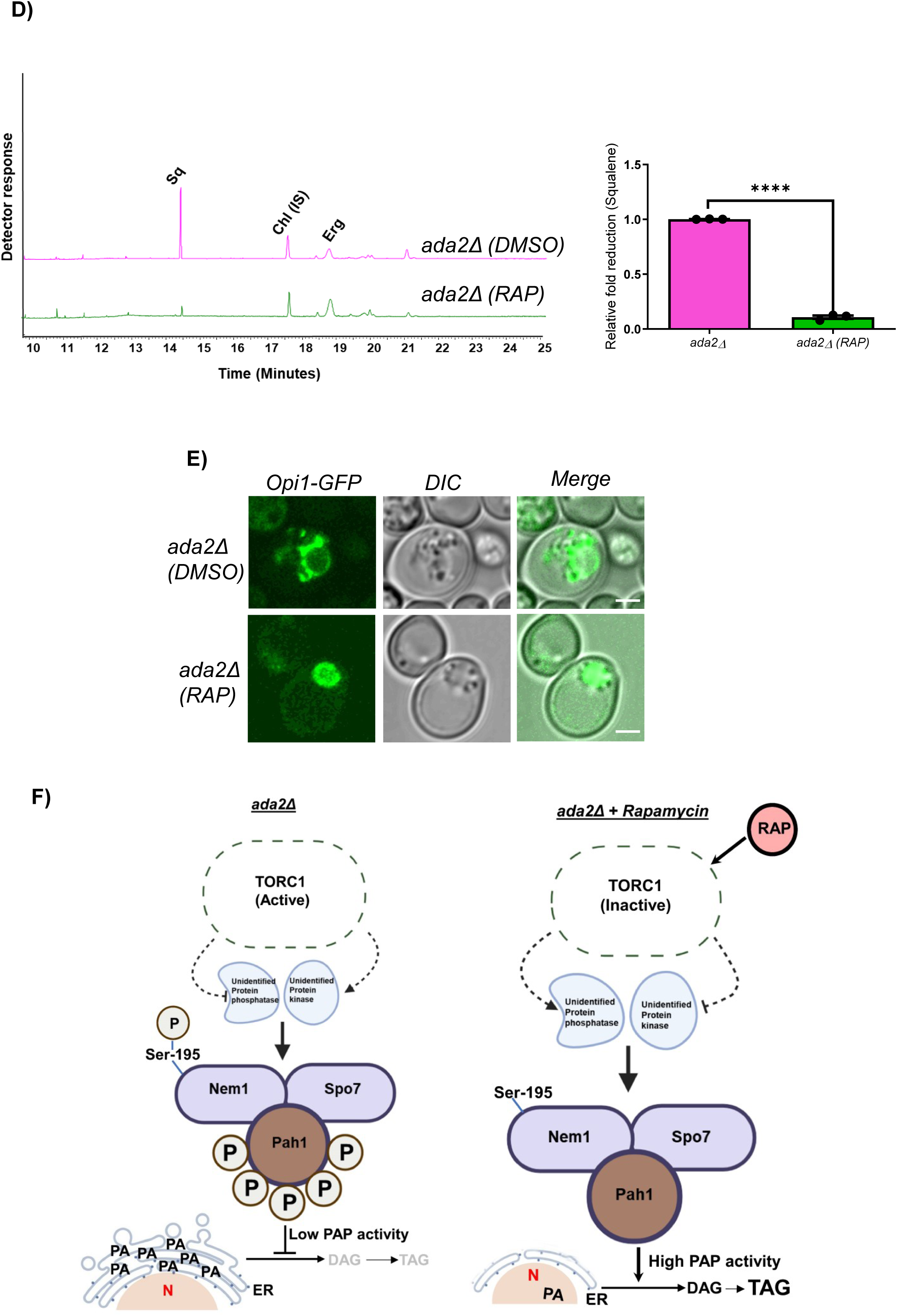
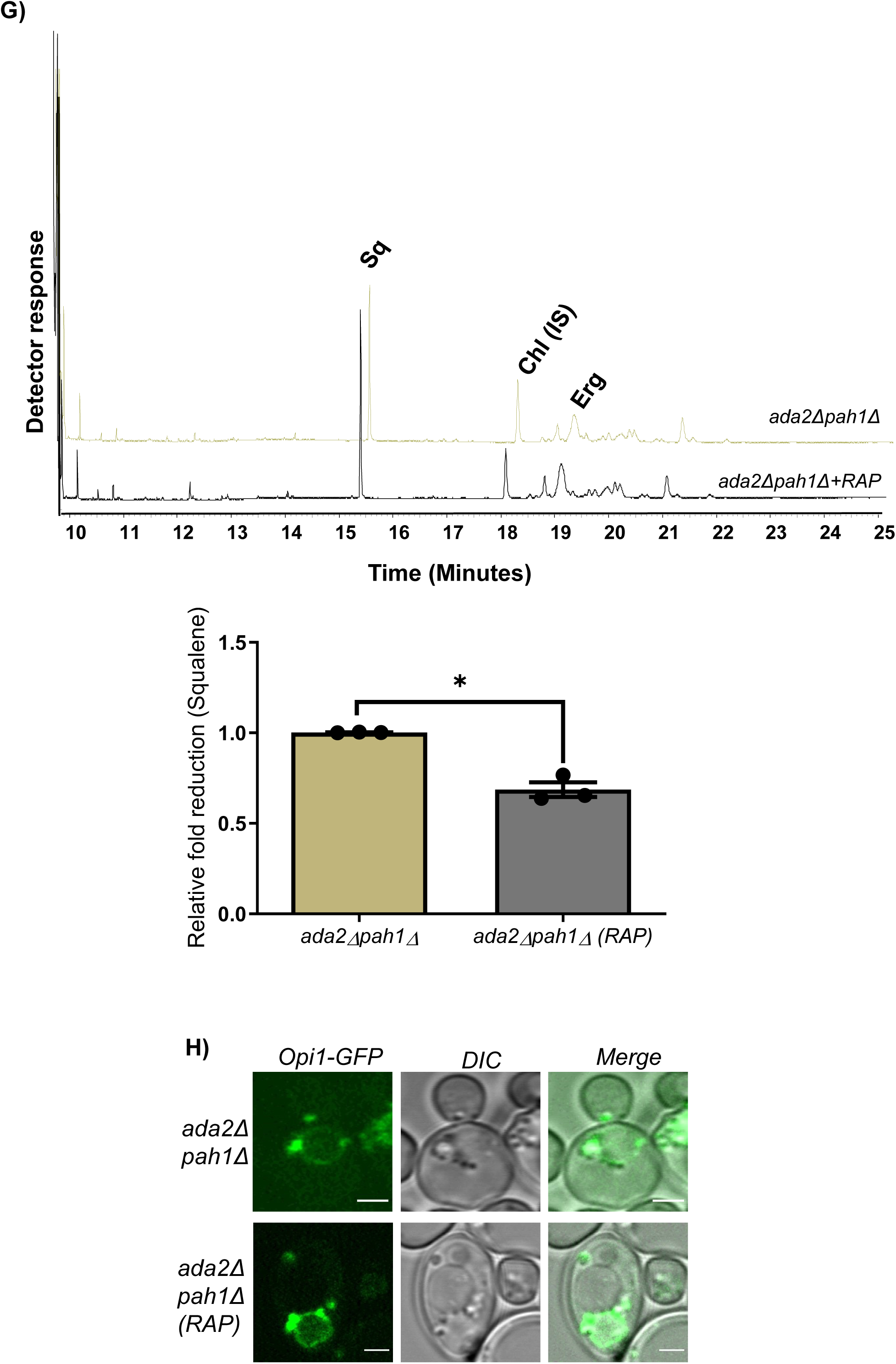
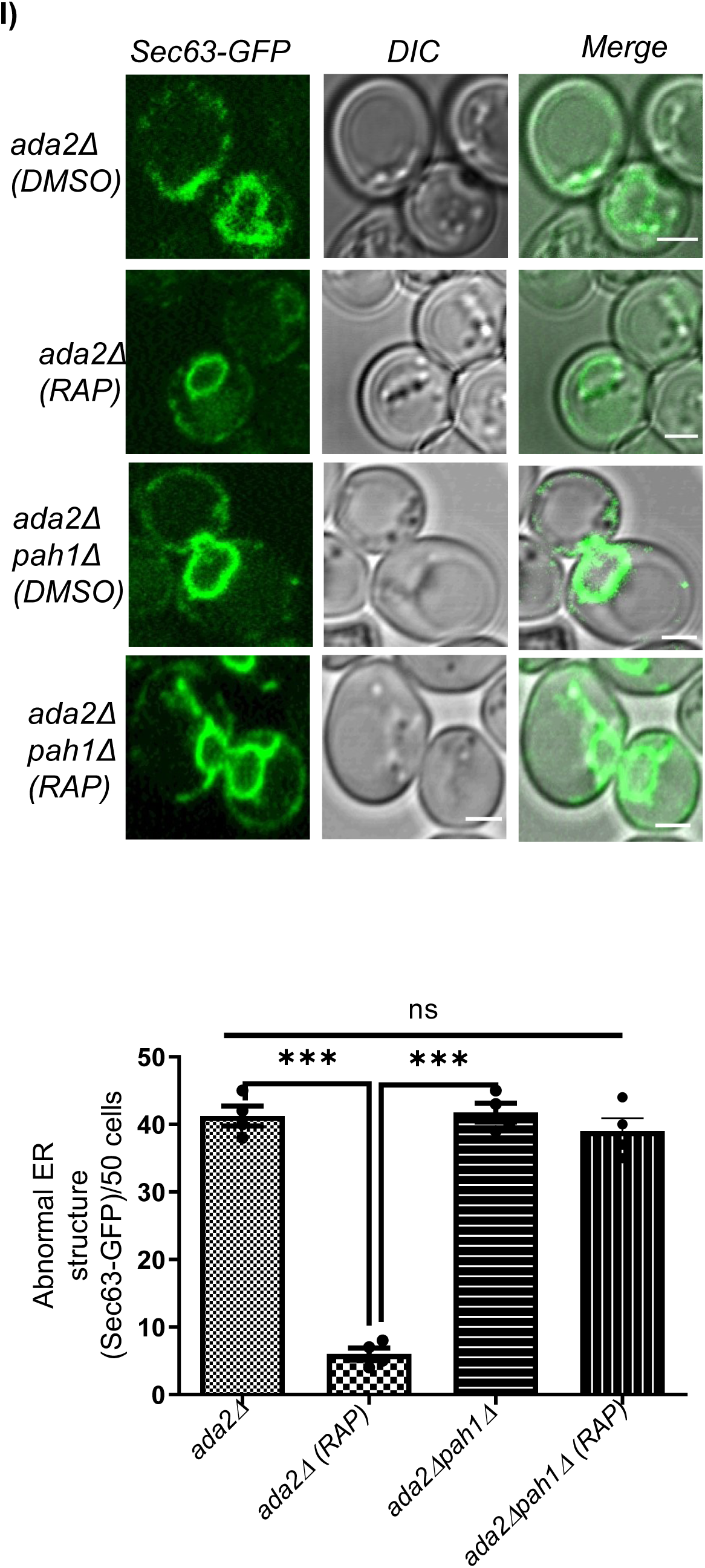
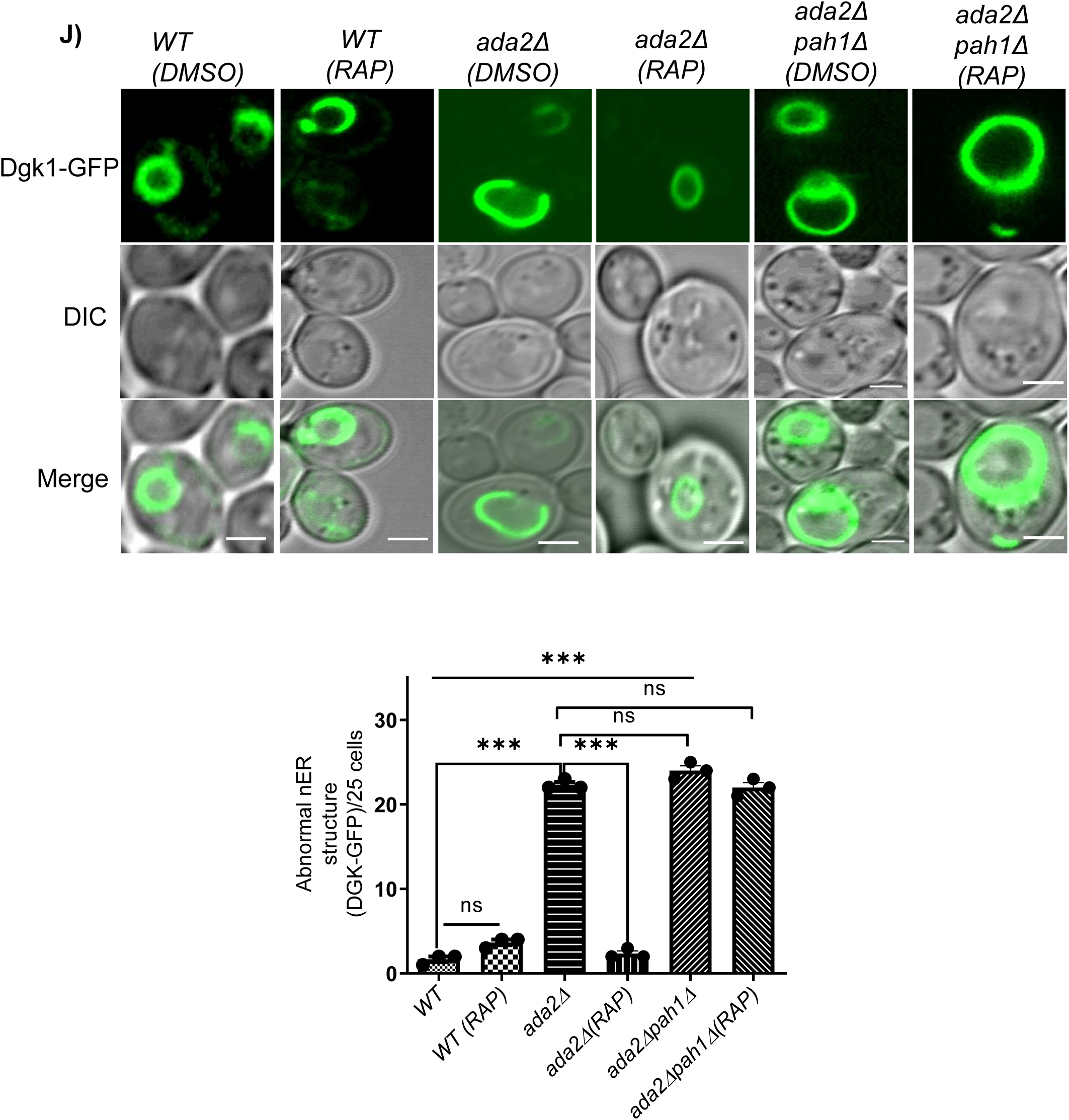
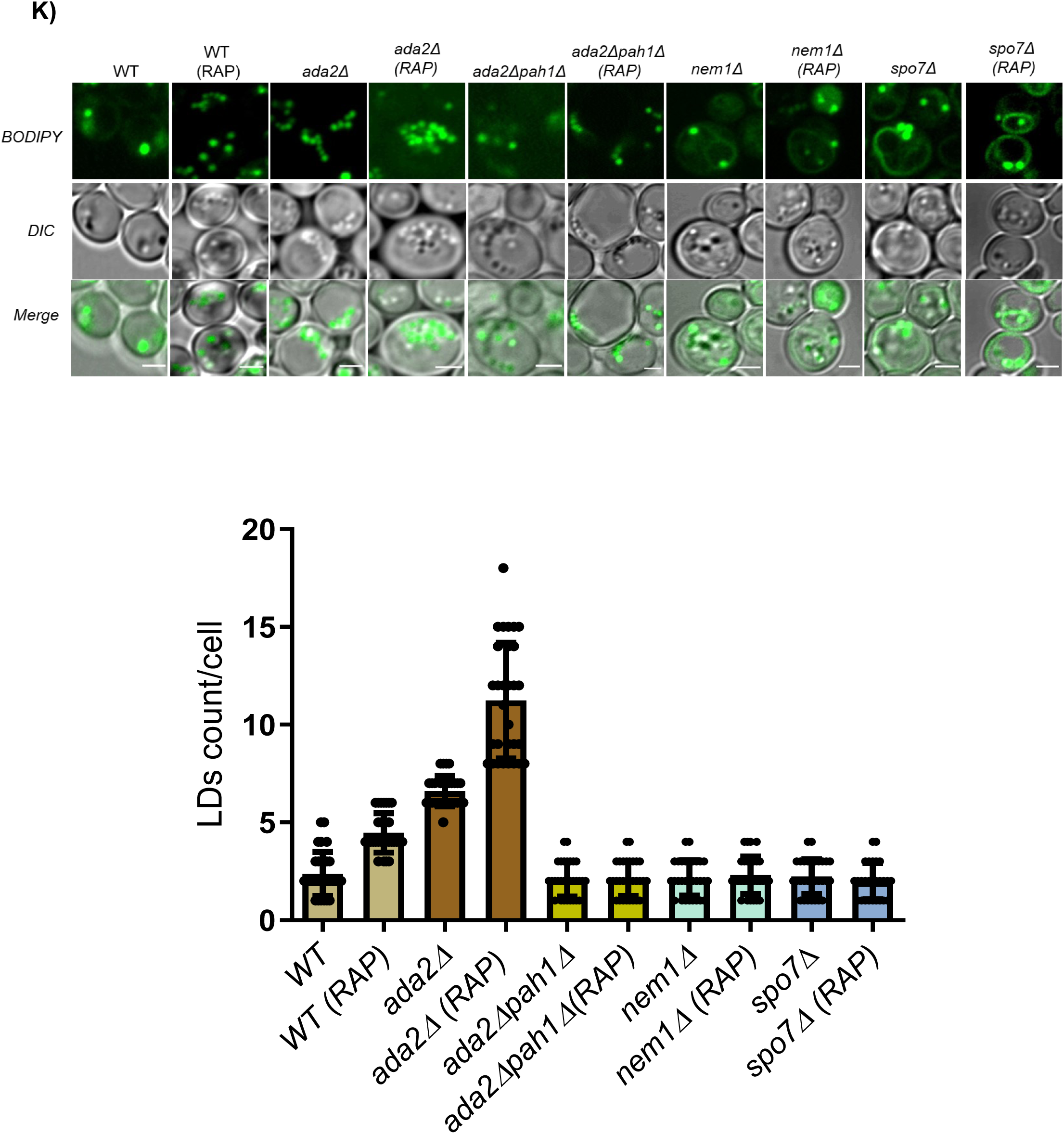
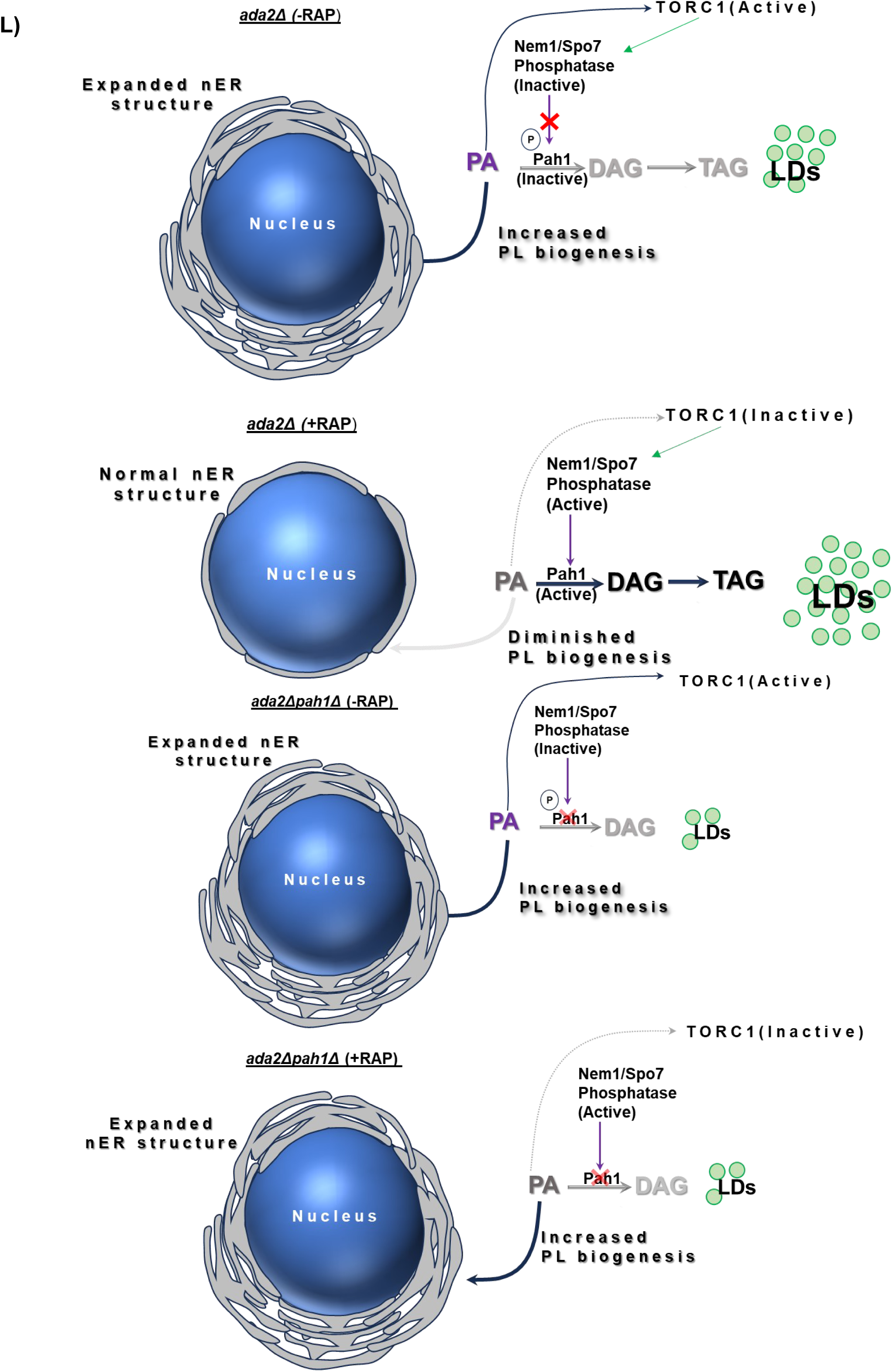
*TORC1* inactivation in *ada2*Δ led to Pah1 activation through the Nem1/Spo7-Pah1 axis, resulting in rapid phosphatidic acid (PA) mobilisation. **A)** mTORC1 activity was assessed in stationary phase *WT*, ada2Δ, and rapamycin-treated cells using the Sch9 NTCB cleavage assay. In *ada2*Δ cells, elevated mTORC1 activity is indicated by the presence of the upper, phosphorylated C-terminal Sch9 band. In contrast, wild-type and rapamycin-treated cells exhibit reduced mTORC1 activity, as shown by lower bands. The experiment was performed in three biological replicates (n=3). **B)** RT-qPCR analysis of Ribosomal Protein of the Large subunit (i.e., RPL30, 32 & 16A) transcripts in *WT* and *ada2*Δ stationary phase cells reveals upregulation of ribosomal gene expression in *ada2*Δ. Wildtype is normalized to one, and the relative fold changes were analyzed. (n=3; Error bars denote ±SD). **C)** Confocal microscopy shows increased lipid droplets (LDs) in rapamycin-treated *ada2*Δ (RAP) cells in the stationary phase. The bar graph provides quantitative lipid droplets (WT, number of cells=33; WT (RAP), number of cells=34; *ada2*Δ, number of cells=27; *ada2*Δ (RAP), number of cells=30 analysed in each of their biological replicates (n=3); Error bars denote ±SD; Scale bar 2 µm). **D)** The GC chromatogram shows reduced squalene (SQ) level in rapamycin-treated *ada2*Δ, indicating reduced MEV-ERG pathway activity, possibly due to rapamycin-induced ER degradation. The bar diagram shows the relative fold reduction of SQ in rapamycin-mediated inactivation of TORC1 in *ada2*Δ (n=3; P value was calculated using Student’s *t* test, ****p ≤ 0.0001; Error bar ±SD). **E)** The confocal image using GFP-Opi1, a PA-specific marker, shows depletion of PA in rapamycin-treated *ada2*Δ (n=3; Scale bar 2µm). **F)** Schematic representation of the proposed mechanism: rapamycin inhibits TORC1, thereby activating the Nem1–Spo7 complex, which dephosphorylates and activates Pah1, promoting PA conversion. **G)** The GC analyses illustrate no significant change in SQ levels upon rapamycin treatment in the *ada2*Δ*pah1*Δ strain compared to the untreated control. These results illustrate the importance of Pah1 in maintaining lipid homeostasis and ER function. The bar diagram represents the relative fold change in SQ. (n=3; Student’s *t* test P value, *p ≤ 0.05; Error bar ±SD). **H)** Confocal microscopy images of Opi1-GFP show PA levels in both *ada2*Δ*pah1*Δ and rapamycin-treated *ada2*Δ*pah1*Δ stationary phase cells. The results indicate that PA is predominantly localized on the nuclear ER, even in the presence of rapamycin. This observation suggests that rapamycin treatment does not alter the subcellular distribution of PA in the *ada2*Δ*pah1*Δ strain. Seven different confocal Z-stack images were captured, and the images were processed using ImageJ software (n=3; Scale bar, 2µm). **I)** Confocal microscopy of Sec63-GFP shows abnormal nuclear ER proliferation in *ada2*Δ*pah1*Δ under both rapamycin-treated and untreated conditions, whereas *ada2*Δ exhibits reduced ER phagy in the presence of rapamycin. The bar diagram depicts abnormal ER structure in *ada2*Δ and *ada2*Δ*pah1*Δ cells treated with or without rapamycin (n=3; Data analyzed by Student’s *t* test, Ns, not significant; Ns p> 0.05, ***p ≤ 0.001; Error bar ±SD; Scale bar 2µm). **J)** Confocal microscopy images of Dgk1-GFP, a nuclear ER marker, show nuclear ER expansion in *ada2*Δ under control conditions. While under rapamycin-treated conditions, *ada2*Δ shows a reduced nER structure. In contrast, *ada2*Δ*pah1*Δ exhibits abnormal nuclear ER structure both with or without rapamycin treatment. The bar diagram depicts the abnormal nER structure observed under rapamycin treatment and control conditions in all yeast strains, and the data were obtained from 25 cells from each biological replicate (n=3, Student’s *t* test P value, Ns, not significant; Ns p> 0.05, ***p ≤ 0.001; Error bar ±SD; Scale bar 2 µm). **K)** Confocal microscopy shows lipid droplets stained with BODIPY (493/503) in various strains. The rapamycin-treated *ada2*Δ*pah1*Δ exhibits reduced LDs as compared with the *ada2*Δ treated with rapamycin (n=3; Scale bar 2 µm; Error bar ±SD). **L)** Diagram illustrating TORC1-mediated regulation of phospholipid biogenesis in *ada2*Δ and *ada2pah1*Δ cells. In *ada2*Δ, rapamycin inactivates TORC1, enabling the Nem1–Spo7 complex to activate Pah1, which converts PA into DAG and ultimately TAG, reducing ER expansion. In *ada2*Δ*pah1*Δ, the absence of Pah1 prevents PA mobilization, leading to continuous PC synthesis and ER proliferation, even under TORC1 inhibition. Abbreviations: SQ, squalene; Chl, cholesterol internal standard; Erg, ergosterol; RAP, rapamycin; DMSO, Dimethyl sulfoxide acts as control; LDs, lipid droplets; Stat., stationary phase; nER, nuclear endoplasmic reticulum; N, nucleus; PA, phosphatidic acid; nER, nuclear endoplasmic reticulum; GC, gas chromatography; PLs, phospholipids.

To confirm the role of Pah1 in this process, we deleted *PAH1* in the *ada2*Δ background. The *ada2*Δ*pah1*Δ strain exhibited persistently high SQ levels even after rapamycin treatment (**Fig. 6G**). Similarly, using Opi1-GFP confocal imaging, we found that excess PA remained on nuclear ER as a bright punctate structure in both control and rapamycin-treated conditions (**Fig. 6H**). This indicates a disrupted TORC1–Nem1/Spo7–Pah1 axis in the absence of Pah1. Additionally, confocal imaging with Sec63-GFP and Sec12-RFP revealed continuous ER expansion in *ada2*Δ*pah1*Δ, regardless of rapamycin treatment (**Fig. 6I**; **Fig. S10**). Similar results were obtained using Dgk1-GFP, a nuclear ER membrane marker, further confirming nuclear membrane expansion in *ada2*Δ*pah1*Δ **(Fig. 6J).** In contrast, reduced lipid droplets were observed in *ada2*Δ*pah1*Δ strain treated with rapamycin, suggesting PA is solely mobilized to membrane biogenesis rather than lipid droplets due to lack of Pah1 activity (**Fig. 6K & L**).

The TORC1-NEM1/SPO7-PAH1 axis is essential for the proper localization of ER phagy receptors ([42]), and nuclear ER-phagy is mainly driven by the nuclear ER-specific receptor, Atg39 ([45]). Since *ada2*Δ cells treated with rapamycin exhibited a rapid decline in SQ levels, we investigated the role of Atg39 in this context. To do so, we deleted *ATG39* in the *ada2*Δ background and quantified SQ levels with and without rapamycin treatment. Unlike *ada2*Δ, the *ada2*Δ*atg39*Δ strain maintained the elevated SQ levels even after TORC1 inhibition (**Fig. S11**). These results confirm that Atg39 is critical for ER phagy, and its deletion leads to high ER-bound pathway activity, including MEV-ERG pathway.

## Discussion

The present study demonstrates that the loss of *ADA2* in HAT complex is associated with altered lipid metabolism, leading to ER expansion and abnormal triterpene/sterol metabolism. We began by examining each HAT-subunit deletion individually, assessing its impact on triterpene/sterol metabolism by measuring SQ metabolite, a key indicator of mevalonate-ergosterol (MEV-ERG) pathway activity ([26]). Among HAT mutant strains, *ada2*Δ displayed significantly elevated SQ levels, consistent with increased MEV–ERG pathway activity and ER expansion as reported in our earlier work ([28]). Using ER-specific markers, we confirmed that *ada2*Δ cells exhibit a visibly expanded ER structure. This expansion may result in enhanced activity of HMG-CoA reductase, an ER-localized enzyme whose activity is linked to SQ synthesis ([46]), leading to high SQ levels. The ER expansion is often associated with ER stress. Supporting this, RT-qPCR analysis revealed an increased expression of *HAC1* and *IRE1*, key markers of unfolded protein response (UPR), along with detectable spliced HAC1, indicating heightened ER stress in *ada2*Δ. This is in line with the known role of ER stress in triggering autophagy in yeast ([47], [32], [28]). We assessed autophagy using GFP-Atg8 and GFP-Pho8Δ60 cleavage assays. Our results confirmed that *ada2*Δ undergoes ER stress-induced autophagy.

The ER expansion and elevated autophagy in *ada2*Δ hints at the possible deregulation of phospholipid (PL) metabolism. Since PLs are essential for membrane biogenesis and ER integrity ([48], [49]), we profiled lipids in *ada2*Δ and found significantly high levels of phosphatidylcholine (PC) and phosphatidic acid (PA), and reduced endogenous inositol levels, pointing to imbalanced PL metabolism. We hypothesize that excess PC in *ada2*Δ promotes ER expansion, which in turn exacerbates ER stress—a connection between PC accumulation, ER biogenesis, and stress has been previously documented in the literature ([50],[51]). To identify the source of this imbalance, we quantified transcripts of key PL-biosynthetic genes, *CDS1* and *INO1,* that showed down-regulation, and hypoacetylation at *CDS1* and *INO1* promoter regions, as revealed from our ChIP assay, further implying reduced Cds1 and Ino1 activities. Because Cds1 converts PA to CDP-DAG, the gateway to PI and other PLs ([52]), its repression explains the excess PA we observed. Further, the elevated Opi3 activity in *ada2*Δ likely results in enhanced PC synthesis through the CDP-choline–independent pathway, ensuring sufficient PC levels under conditions where the CDP-choline pathway may be limited or less efficient. Thus, impaired *CDS1/INO1* expression traps PA upstream, skews flux toward PC via increased Opi3 activity, and fuels both ER expansion and stress in *ada2*Δ. Our Opi3-GFP processing assay underscores Opi3’s functional stability in the *ada2*Δ background, where minimal free GFP release, indicating resistance to vacuolar turnover or proteolytic processing. This stability aligns with elevated PC accumulation.

Since *ada2*Δ cells show low *CDS1* expression, in addition to ER expansion, heightened autophagy, and excess PA, we overexpressed *CDS1* in *ada2*Δ, which restores ER morphology and decreases both ER phagy and general autophagy. This consequently also reduces SQ levels. RT-qPCR further revealed that *INO1* transcripts increased in the *ada2*Δ strain expressing *CDS1*, indicating reactivation of the PI biosynthetic pathway. These results suggest that Cds1 overproduction not only diverts PA into the CDP-DAG branch but may also transcriptionally up-regulate *INO1*. Thus, impaired *CDS1* and *INO1* expressions result in increased PA accumulation in *ada2*Δ that leads to abundant PC synthesis. However, the precise route by which surplus PA feeds into PC remains to be elucidated.

An intricate relationship exists between SAGA-mediated histone acetylation, inositol metabolism, and PC synthesis. Cytosolic inositol limitation is known to elevate levels of PA and its downstream product, PC ([39],[35],[53]), and Gcn5/SAGA complex dependant acetylation regulates *INO1* expression ([20]). The low *INO1* expression in *ada2*Δ suggests that *ADA2,* a subunit of the SAGA/HAT complex, plays an important role in inositol regulation. Restoring *INO1* expression (or supplying exogenous inositol) in *ada2*Δ mirrors the effects of *CDS1* overexpression, leading to reduced autophagy, restored ER morphology, and a decrease in SQ levels. This is supported by our GFP-Opi1 data, which indicates depleted PA levels when Ino1p activity is increased or exogenous inositol is added in *ada2*Δ. Furthermore, it has been reported that inositol addition depletes ER-localized PA by promoting its conversion to PI ([35]). These studies demonstrate a link connecting Ino1p activity to autophagy regulation through changes in phospholipid levels, particularly PA. When cells are inositol-deficient, this condition can trigger the unfolded protein response (UPR), leading to expansion of the ER membrane to accommodate increased levels of ER-resident proteins, including those involved in the mevalonate–ergosterol (MEV–ERG) pathway ([54],[55]). Because PA depletion shifts Opi1 into the nucleus, it represses the Ino2/Ino4 phospholipid-transcription complex, thereby tempering phospholipid synthesis and ER expansion ([38]). These findings support a model in which ADA2 maintains lipid balance by enabling *CDS1* and *INO1* transcription—likely through SAGA-mediated chromatin remodelling. When ADA2 is lost, inositol scarcity and PA accumulation trigger ER proliferation, heightened autophagy, and elevated triterpene/sterol levels. Precisely how endogenous inositol levels feed back to squalene biosynthesis remains an open question.

Lipid droplet analysis using BODIPY staining showed that there were more lipid droplets in *ada2*Δ during the stationary phase. Lipidomic data, however, showed that *ada2*Δ had lower triglycerides (TAG). It is possible that the enhanced lipid droplets are caused by high SQ levels that are sequestered as lipid droplets rather than high TAG ([56],[57]). Knowing that *ada2*Δ displayed altered lipids, specifically increased PA and PC and decreased neutral lipid, TAG, we hypothesized that the Target of Rapamycin1 (TORC1) complex may play a role in PA mobilization.

PA activates TORC1 ([58], [59]), which in turn inhibits Pah1 by suppressing the Nem1–Spo7 phosphatase complex ([60],[61]). When TORC1 is inhibited, Nem1–Spo7 becomes active and dephosphorylates Pah1, restoring its function. Given the significant TORC1 activity in *ada2*Δ stationary phase cells, as confirmed by the Sch9 NTCB cleavage assay, we hypothesized that low Pah1 activity in *ada2*Δ may be responsible for the reduced triglyceride synthesis even in the presence of substantial PA levels. To test whether mobilizing excess PA to TAG through Pah1 activation would affect squalene (SQ) levels, we inhibited TORC1 using rapamycin. Interestingly, with rapamycin treatment, there was a notable decrease in SQ and an increase in TAG, signifying that high Pah1 activity mobilizes excess PA majorly to TAG synthesis. Strong nuclear-localized GFP-Opi1 signals further confirmed PA depletion on the nuclear ER membrane. Additionally, rapamycin treatment led to increased lipid droplet formation, supporting the idea that Pah1-mediated conversion of PA to DAG and subsequently to TAG was restored via the reactivated Nem1–Spo7 complex ([62]). To directly assess Pah1’s role, we deleted *PAH1* in the *ada2*Δ background. Upon rapamycin treatment, the *ada2*Δ*pah1*Δ strain showed no reduction in SQ levels, and PA remained enriched on the nuclear ER membrane, comparable to *ada2*Δ. Moreover, reduced lipid droplet formation in rapamycin-treated *ada2*Δ*pah1*Δ, *nem1*Δ, and *spo7*Δ strains further supported that, in the absence of a functional TORC1–Nem1/Spo7–Pah1 axis, PA is redirected primarily toward membrane biogenesis rather than lipid storage. In short, the excess PA in *ada2*Δ does not mobilize to CDP-DAG or PI synthesis due to poor *CDS1* and *INO1* expression, and concomitant PA-mediated TORC1 activation makes Pah1 inactive. PA is predominantly shunted into PC synthesis, contributing to ER membrane expansion and substantial squalene accumulation.

*ATG39* encodes an ER receptor protein that facilitates the ER phagy ([63]). In *ada2*Δ, the drastic reduction of SQ was observed when cells were treated with rapamycin. This effect is likely due to rapamycin’s dual role in promoting ER phagy and suppressing PC biosynthesis by activating Pah1, which depletes PA, restores normal ER morphology, and reduces SQ accumulation. Since ER is essential for MEV-ERG pathway activity, we investigated whether loss of *ATG39* would impact rapamycin-induced SQ reduction. To this end, *ATG39* was deleted in the *ada2*Δ background. Interestingly, the *ada2*Δ*atg39*Δ strain retained elevated SQ levels even after rapamycin treatment. It is likely that sustained MEV–ERG pathway activity.

To summarize, the *ADA2* of HAT complex acts as an autophagy inhibitor through transcriptional regulation of *CDS1* and *INO1.* Due to impaired Cds1 and Ino1 activities, PA and PC accumulation exceeds the normal limits. As a result, surplus PA is primarily diverted to PC synthesis, driving ER membrane expansion and promoting SQ overproduction via increased MEV–ERG pathway activity. In addition, excess PA levels maintain TORC1 in an active state that negatively regulates Pah1 activity. When Pah1 is inactive in stationary-phase cells, diacylglycerol (DAG) produced by other PA phosphatases may be directed more toward phospholipid synthesis than TAG formation ([64],[44]). Under inositol-limiting conditions in stationary phase, non-canonical routes may also contribute to PC synthesis ([65]). However, there are no definitive reports explaining how excess PA is partitioned between phospholipid and neutral lipid synthesis under conditions of *CDS1* and *INO1* repression. Further research is required to clarify how PA is routed toward PC in *ada2*Δ, as this process profoundly influences multiple aspects of cellular metabolism and homeostasis.

Beyond the significance of understanding ADA2’s role in yeast, our findings may have broader implications for understanding the role of ADA2 in regulating lipid metabolism and its wider metabolic consequences in eukaryotes. Since lipid and sterol imbalances are associated with various human disorders, including metabolic syndrome, this work may contribute to a deeper understanding of metabolic disease states and support the development of future therapeutic strategies. The conservation of ADA2 function across eukaryotes is emphasized by the existence of two human homologues, TADA2a and TADA2b. The human ADA2b mirrors the yeast counterpart in structure and function, suggesting that the mechanistic understanding derived from yeast ADA2 studies is directly translatable to mammalian systems ([22]). It has been postulated previously that the possible interaction of yeast adaptor proteins appears to influence the activity of human p53 in yeast, suggesting that human ADA2 may similarly modulate p53 function ([66]). If hADA2 indeed regulates p53 activity, gaining a comprehensive understanding of the relationship between hADA2 and p53 could provide valuable insights into the mechanisms underlying metabolic diseases. Also, it has been reported that deletion of human ADA2 (TADA2b) increases β-globin production, that leads to reduce the clinical severity of sickle cell disease and β-thalassemia metabolic disorder ([67]). In conclusion, our study not only highlights the critical role of ADA2 in yeast but also suggests its broader significance in regulating lipid metabolism and maintaining metabolic balance across eukaryotes, and provides valuable insights into the molecular basis of metabolic disorders for future therapeutic strategies.

## Methods and materials

### Growth conditions, Yeast strains, plasmids

The various yeast strains used in this study (Table S1) were grown mainly in a synthetic defined (SD) medium (0.67% yeast nitrogen base with ammonium sulfate, 2% dextrose, and amino acids). The plasmids used in this study are listed in Supplementary Table 2. We utilized the previously described GFP-Pho8Δ60[68], pRS316-ATG40-GFP[69], and YCpLAC33-SEC63-GFP[70] plasmids for our experiments to investigate general/macroautophagy and ER-phagy. For galactose overexpression studies, the yeast cultures were grown at 30 °C till the cells reached 0.8 OD_600_ units. The cultures were centrifuged, washed twice, and shifted into an SD medium containing 2% galactose. After 24 hours of induction, the cells were harvested and used for subsequent analysis. The empty vector transformed cells were used as appropriate controls in experiments. For rapamycin treatment, the cells were grown to the log phase, and rapamycin 200 ng (1mg/ml stock in DMSO) was added. The cells were grown to a stationary phase for subsequent analysis. For external inositol addition experiments, the cells were grown for 24 hrs, and inositol (200 µm) was added. The gene knockouts in yeast strains were carried out as described ([71]). The plasmid pUL9 was used to perform a marker swap in yeast strains to overcome the limitation of having the same marker[72]. All the genetic transformations in yeast were carried out using the lithium acetate method as described earlier ([73]). The genomic C-terminal GFP tagging was performed as described earlier ([74]), and the DNA oligos are provided in Supplementary Table 3.

### Metabolite extraction and GC-FID/MS analysis

The stationary phase cells were used for the extraction of squalene from various experimental conditions. The metabolites were extracted using the alcoholic KOH method followed by dodecane extraction as previously described ([26]). The total 1µl was injected with a split ratio of 1:10. The GC chromatographic separation, conditions, and analysis were carried out as described earlier ([27], [28]).

The intracellular inositol was quantified using GC-MS as described earlier ([75]). In brief, an equal amount of yeast cells was extracted with 1000 µL of chilled 80% methanol–water (−20°C). The mixture was sonicated (53 kHz, 5 min), vortexed (1000 rpm, 25 min), and centrifuged (14,000 rpm, 15 min, 4°C). The total metabolite extraction was repeated three times, and the pooled supernatants were evaporated using a vacuum concentrator (Eppendorf Concentrator Plus). The dried extracts were derivatized for GC-MS analyses. The analysis was performed on an Agilent 8890 GC system coupled with a 7010C triple quadrupole MS and a PAL RSI 85 autosampler. The GC separation was achieved using an HP5-MS column (30 m × 0.25 mm × 0.25 µm). A 1 µL aliquot was injected (split ratio 10:1) with helium as carrier gas (0.9 mL/min). The oven temperature was programmed from 70°C (2 min hold) to 295°C at 12.5°C/min, then to 320°C at 25°C/min (3 min hold). Injection port and transfer line temperatures were set at 270°C and 280°C, respectively. The mass spectrometer was operated in full scan mode (40–500 amu) with 70 eV ionization energy, 230°C source temperature, and 150°C quadrupole temperature. Data were analyzed using the Automated Mass Spectral Deconvolution and Identification System (AMDIS v2.0, NIST) for metabolite identification.

### RT-qPCR analyses

Total RNA from different yeast strains was isolated using TRIzol^TM^ Reagent (Invitrogen). Briefly, cells (30 OD_600_) were harvested from a stationary phase culture by centrifugation and washed twice with sterile water. The 0.5 g of acid-washed glass beads and 700 µL TRIzol™ reagent were added to the cell pellet and incubated in ice for 10 min. The pellet was subjected to vigorous vortexing for 45 min for cell lysis at 4 °C. Approximately 200 µL of molecular-grade CHCl_3_ was added and gently inverted 2-3 times. The cell suspension was incubated for 10 min at room temperature, followed by centrifugation at 13,000 rpm for 15 min at 4°C. The supernatant was carefully transferred into fresh 1.5 mL RNase-free tubes. The total RNA was precipitated with 0.5 mL of isopropanol and incubated for 10 min. Following incubation, the tubes were centrifuged at 13,000 rpm for 15 min at 4°C to pellet down the total RNA. The resultant pellet was air-dried and dissolved in 26 µL of RNase-free water, followed by DNase1 treatment. The quality of the total RNA was checked using a bio-spectrophotometer, which was used for cDNA synthesis. The high-capacity cDNA reverse transcription kit (Applied Biosystems, 4319981) was used to synthesize single-stranded cDNA from 2 µg of RNA. The reverse transcription (RT) master mix comprises RT random primers (10X), RT buffer (10X), dNTP mix (100mM), and Multiscribe Reverse transcriptase enzyme (50U/µL). The RT was performed according to the manufacturer’s instructions. The quantitative transcript analysis was carried out using a Step One qPCR machine, with Power SYBR Green (Applied Biosystems, 4367659) using gene-specific RT primers mentioned in Supplementary Table S4. Glyceraldehyde-3-phosphate dehydrogenase (*TDH3*) or actin was used for the normalization of gene expression.

The *HAC1* splicing assay was carried out by performing PCR with *HAC1-*specific primers. The amplified products were resolved using 1.8% agarose gel electrophoresis. For ER stress conditions, we treated the cells with 8 mM Dithiothreitol (DTT) as previously described with minor modification ([76]). In our study, we treated cells with DTT during log phase and allowed them to grow till the stationary phase. The cDNA was constructed as described above.

### Confocal microscopy

Approximately 3 OD_600_ cells were collected from various media conditions for the confocal imaging. The images were captured using confocal microscopy Zeiss LSM 880 AxioObserver with the help of a 100x immersion oil objective. The GFP (excitation: 488 nm, emission 540 nm; detection wavelength between 491-589 nm) and RFP (excitation: 561 nm, emission: 629 nm; detection wavelength between 562-696 nm) signals were detected and analyzed using ImageJ software ([77]). For lipid droplet staining, the 3 OD_600_ cells grown in SD medium were collected and washed with 1X phosphate buffer saline (pH 8.0), followed by lipid droplet staining with 1X PBS buffer containing BODIPY 493/503™ (1 µg/µl) for 30 min in the dark at 30°C. The stained cells were washed twice with 1X PBS and mounted on agarose (2% w/v) pads for microscopy.

### Immunoblotting experiments

The three OD units of TCA precipitated yeast cells were separated on 12% SDS-PAGE. The proteins were transferred to nitrocellulose or PVDF membrane. The protein transfer efficiency was checked using Ponceau S dye, which stains protein bands on the membrane. The membrane was incubated with primary anti-GFP antibody (Roche, catalog no. 11814460001, 1:3000 dilution) at 4 °C overnight. The membrane was washed with 1X PBST to remove unbound primary antibodies three times (10 minutes each time), and the Anti-mouse HRP conjugated IgG secondary antibodies (Bio-Rad, catalog no. 1706516, 1:10,000 dilution) were added and incubated for one hour. The unbound secondary antibodies were washed from the membrane with 1X PBST three times (10 minutes each). Finally, a Western blot was developed using Immobilon® ECL Ultra Western HRP Substrate (WBULS0100; Merck Millipore). The images were captured using the G: Box Chemi XRQ imager (Syngene).

### Total lipid extraction and analyses

The total lipids were extracted from all the yeast strains using the Folch method ([78]). The 30 OD_(600)_ cells were pelleted down by centrifugation at 5,000 rpm for 5 min at 4 °C. The cell pellets were washed twice with sterile water and lysed in 1ml Y-PER™ Yeast Protein Extraction Reagent (Thermo Scientific, 78991) with 1g of sterilized glass beads (Sigma, G8772) to isolate the crude protein lysate by continuous 1 min vortexing and 1 min ice incubation for 45 min. The protein was quantified using Bradford Reagent (Sigma, B6916). The 5 mg protein crude lysate was subjected to chloroform:methanol: acidified water (1:2:1 v/v) and vortexed for 45 min at room temperature, followed by centrifugation for aqueous and organic phase separation. The lower organic layer was collected and washed three times with acidified water (2 % orthophosphoric acid). The organic layer was dried completely using a vacuum concentrator (Concentrator Plus; Eppendorf). The dried total extracted lipids were resuspended in chloroform and methanol (30 µl:15 µl, v/v) for TLC separation. For two-dimensional thin-layer chromatography as described earlier ([79][80]), the sample was applied as a single spot on a TLC plate (silica gel 60, Merck). For the first-dimension separation, the TLC chamber was equilibrated with CHCl_3_: MeOH: 25% ammonia (65:35:0.5, v/v/v) and used as a solvent system. The TLC plate was carefully removed, dried, and placed in the other TLC chamber, which contained a solvent system, chloroform/acetone/methanol/glacial acetic acid/water (50:20:10:10:5, v/v/v/v) for second dimension separation. After the run was completed, the separated phospholipids were identified by immersing the plate in an iodine vapor-saturated chamber for 10-20 minutes. The spots corresponding to PC were identified. The PC levels were quantified using ImageJ software.

### Yeast lipid profiling using high-resolution accurate LC-MS

The total lipids from an equal amount of stationary phase yeast cells (∼30 OD_600_) were extracted as described earlier ([81]). The extracted yeast lipid samples were prepared for analysis by dissolving them in 200 μL of a chilled 1:1 (v/v) solution of absolute acetonitrile and methanol. The various lipid components were then separated using reverse-phase chromatography, a method adapted from previous study ([82]). Lipidomics profiling was performed using a Vanquish UHPLC system coupled to an Orbitrap 240 mass spectrometer (Thermo Scientific). The LC separation was carried out using the same conditions and equipment as reported earlier ([83]). In brief, reversed-phase chromatography method utilized a Zorbax Eclipse Plus C18 column (2.1 x 150 mm, 1.8 μm) with a gradient mobile phase. The mobile phase consisted of solvent A (60% acetonitrile and 40% water) and solvent B (90% isopropanol and 10% acetonitrile). The gradient solvents, gradient profile, LC conditions, and MS parameter settings used in this study were the same as those as previously described[84]. The Xcalibur 4.5 (Thermo Scientific) software was utilized to record and analyse the MS data. Additionally, lock-mass correction was carried out using common low-mass contaminants in each sample run to improve calibration. HCD collision energies of 20, 35, and 60 were used to normalize the collision energies for data-dependent analysis. Lipids were identified and analyzed using LIPIDMAPS and LipidBlast ([85]).

### Sch9 NTCB Cleavage mTORC Assay

To assess mTORC1 activity, Sch9 was C-terminally tagged with 6xHA in both wild-type and *ada2*Δ cells using the standard lithium acetate transformation method. Transformants were selected on nourseothricin-containing media and grown for 24 hours to reach the stationary phase. For rapamycin treatment, cultures were grown to logarithmic phase and treated with rapamycin at the indicated concentration. To prepare protein extracts from stationary phase cells (∼ 40 OD₆₀₀) were harvested and subjected to protein precipitation using 6% trichloroacetic acid (TCA). The resulting pellet was washed with ice-cold acetone, flash-frozen in liquid nitrogen, and stored at –80°C until further processing ([40] [86] [87]). Cell lysis was performed using urea lysis buffer (50 mM Tris-HCl, pH 7.5, 5 mM EDTA, 6 M urea, 1% SDS, 1 mM PMSF) supplemented with phosphatase inhibitors (50 mM sodium fluoride, 2 mM sodium orthovanadate). Cells were disrupted using a bead beater (3 minutes), and lysates were incubated at 75°C for 10 minutes, followed by centrifugation at high speed for 5 minutes at room temperature. The supernatant (200 µL) was subjected to chemical cleavage using 2-nitro-5-thiocyanobenzoate (NTCB) in 0.5 M CHES buffer (pH ∼10.5) overnight at room temperature. Following cleavage, samples were mixed with 2× SDS-PAGE sample buffer containing 20 mM TCEP and 0.5× phosphatase inhibitor cocktail (excluding traditional reducing agents such as DTT or β-mercaptoethanol). Samples were resolved on 8% SDS-PAGE and transferred to PVDF membranes for western blotting. Blots were probed with anti-HA rabbit primary antibody (1:2000 dilution), followed by a horseradish peroxidase (HRP)-conjugated anti-rabbit secondary antibody (1:4000 dilution). Band shifts corresponding to phosphorylated and dephosphorylated forms of C-terminal Sch9 were visualized using chemiluminescent detection (ChemiDoc).

### ChIP analyses

The chromatin immunoprecipitation (ChIP) experiment was performed using previously reported methods ([88][89]). Yeast cells were cultured in SD media for 24 hrs until the stationary phase, and then treated with 1% formaldehyde to crosslink DNA–protein interactions *in vivo*. Following a brief incubation, glycine was added to quench the crosslinking, and cells were harvested and washed with 1X PBS. The cell pellet was resuspended in lysis buffer (including protease inhibitors) and sonicated under optimal conditions to fragment chromatin to an average size of ∼200 bp. Following this, the cell lysate was subjected to centrifugation to remove cell debris, and 1000 µL of the resulting soluble chromatin was collected for immunoprecipitation. The anti-histone H3 (acetyl K14) antibody-ChIP Grade (ab52946, Abcam) was added to the lysate and incubated (at 4 °C, with rotation) for immune precipitation. 50 μL of Protein G Plus-agarose suspension (IP04; Merck, Germany) was added, followed by a series of washes with different wash buffers to reduce nonspecific binding. The chromatin-antibody bound complexes were eluted, and crosslinks were reversed by heating at 65°C overnight, along with RNAse A treatment, followed by proteinase K. The DNA was extracted with 500 μL of phenol:chloroform: isoamyl alcohol solution. The extracted DNA was precipitated with 100% ethanol and dried. The DNA sample was resuspended in 20 µL of milliQ water. To conduct downstream analysis, 50 ng of DNA was used as template in a 23-cycle PCR with locus-specific primers (*CDS1* or *INO1* promoter regions), along with a 50ng of DNA from “input” DNA sample (purified prior to immunoprecipitation) that acts as a control. PCR products were resolved on a 2 % agarose gel to assess the enrichment of H3 acetylation (K14) at the targeted loci. The entire ChIP procedure was repeated in triplicate, yielding consistently reproducible results. Primers used for PCR analysis were as follows: CDS1: 5’-GGGAAGATACTTGGCAAGAAGTGC-3’,5’-CGGATACCCTTCTTGAAAATTTTTCAGATGG-3’; INO1:5’-CATGCCGCATTTAGCCGC-3’, 5’- CCTTTTGTTCTTCACGTCCTTTTTATGAA-3’.

### Statistical analysis

The data is represented as the mean (±SD) from the three independent experiments (n=3) and is analyzed using GraphPad Prism version 5.01 for Windows, GraphPad Software, La Jolla, California, USA, www.graphpad.com. For the comparison of data between two strains, Student’s t-test was performed, whereas one-way ANOVA (Dunnett’s post-test and Bonferroni post-test) was used to analyze multiple data, and two-way ANOVA was used for gene expression analyses. Generally, p-values <0.05, <0.01, and <0.001 were considered statistically significant.

## Abbreviations

Ada2: Adaptor2
SQ: Squalene
ER: Endoplasmic reticulum
PA: Phosphatidic acid
PC: Phosphatidylcholine
PI: Phosphatidylinositol
TORC: Target of Rapamycin Ser/Thr kinase complex
SAGA: Spt-Ada-Gcn5 acetyltransferase
HAT: Histone AcetylTransferase
TAG: Triacylglycerol
DAG: Diacylglycerol
*CDS1*: Phosphatidate cytidylyltransferase 1
*INO1*: Inositol-3-phosphate synthase
ATG: Autophagy
GAL: Galactose
*HAC1*: Homologous to Atf/Creb1 gene
*IRE1*: Inositol Requiring gene1
GFP: Green fluorescent protein
*KAR2*: KARyogamy gene2
PLs: Phospholipids
*OPI1*: Overproducer of Inositol1 gene
Sec: SECretory
*NEM1*: Nuclear Envelope Morphology1 gene
*SPO7*: SPOrulation gene7
*PAH1*: Phosphatidic Acid phosphohydrolase/phosphatase
*DGK1*: Diacylglycerol kinase1
UPR: unfolded protein response
PHO8: PHOsphate metabolism8 gene
ERG: Ergosterol
MEV: mevalonate
WT: Wildtype
ChIP: Chromatin Immunoprecipitation
GC-FID: Gas Chromatography with Flame Ionization Detector
ERG: Ergosterol
Sch9: Serine/threonine-protein kinase
RAP: Rapamycin
DTT: Dithiothreitol
RPL: Ribosomal Protein of the Large subunit
NTCB: 2-Nitro-5-thiocyanatobenzoic acid
SD: standard deviation
LDs: lipid droplets
nER: nuclear endoplasmic reticulum.

## Author Contributions

VRDK designed the experiments and supervised the complete study and drafted the manuscript. AR conducted all the experiments and compiled the data pertaining to the study. NNN was involved in making DNA constructs used in this study. MJR was involved in lipid analysis work. RM and MA helped with designing autophagy assays and the discussion. AKT and RCH carried out lipidomic analysis work.

## Acknowledgments

All authors are grateful to the Director of the CSIR-Central Institute of Medicinal and Aromatic Plants for his continuous support and encouragement. This research was supported by the Department of Biotechnology (DBT), Ministry of Science & Technology, Government of India (No. BT/PR47750/BSA/33/72/2024). AR expresses gratitude to DST – INSPIRE [No. DST/INSPIRE Fellowship/2019/IF190108], New Delhi, for the financial support provided throughout her PhD studies. AR is grateful to Jawaharlal Nehru University, New Delhi, for PhD registration. NNN is grateful to the ICMR for granting a senior research fellowship [No. 45/01/2022-BIO/BMS]. VRDK, AR, and NNN thank CSIR FTT project (No. FTT-FTC/02/05/IMD/2023) for financial assistance. Authors are thankful to Dr. Sunil Laxman, DBT-Institute for Stem Cell Science and Regenerative Medicine, Bangalore, India, for his generous help in conducting Sch9 NTCB cleavage assay at his research facility. The authors are thankful to Kalyani Sai Reju for editing the manuscript.

## Data Availability statement

Data supporting the findings of this study can be obtained from the corresponding author upon reasonable request.

## Supporting Information

**Figure S1.** Transcript levels of key genes of mevalonate-ergosterol (MEV-ERG) pathway in wildtype and histone acetyltransferase (HAT) complex mutants.

**Figure S2.** Transcriptional analysis of autophagy pathway genes in wildtype versus *ada2*Δ cells.

**Figure S3.** Comparative lipidomic profiles of wildtype (*WT)* and *ada2*Δ *strains*.

**Figure S4.** Transcript and protein expression levels of key genes in the phospholipid biosynthesis pathway in wildtype and *ada2*Δ cells.

**Figure S5.** Analysis of *CDS1* and *INO1* gene expression, and myo-inositol levels in wildtype and *ada2*Δ.

**Figure S6.** mRNA transcript levels of phospholipid pathway genes in wildtype, *ada2*Δ *and ada2*Δ *pGAL1-CDS1* strains.

**Figure S7.** Overexpression of *INO1* modulates ER-phagy, general autophagy, and phospholipid pathway gene expression in wildtype and *ada2*Δ.

**Figure S8.** Exogenous inositol supplementation restores transcript levels of phospholipid pathway genes in *ada2*Δ.

**Figure S9.** Comparative neutral lipid profiling of wildtype and *ada2*Δ *±* rapamycin.

**Figure S10.** Confocal microscopy analysis shows ER morphology in wildtype, *ada2*Δ and *ada2*Δ*pah1*Δ cells with or without rapamycin treatment.

**Figure S11.** Gas chromatogram shows no reduction in squalene (SQ) level in *ada2*Δ*atg39*Δ and *ada2*Δ*atg39*Δ*-*treated rapamycin strains.

**Supplementary Table 1.** List of yeast strains use in this study

**Supplementary Table 2.** List of plasmids used in study

**Supplementary Table 3.** List of cloning and knockout primers used in this study.

**Supplementary Table 4.** List of RT qPCR primers used in this study.

