## Supplementary figures for "Decoding the role of ADAptor2 (ADA2) of HAT complex in autophagy and phospholipid metabolism to maintain ER homeostasis and triterpene regulation"

**Supplementary Information**

**S1**

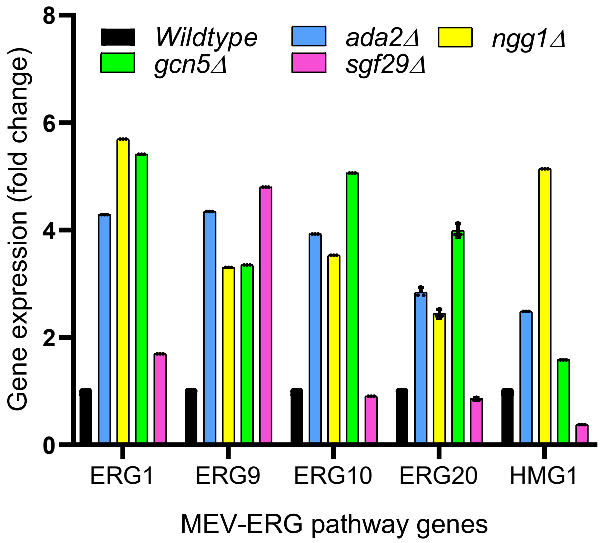


**Figure S1. Transcript levels of key genes of mevalonate-ergosterol (MEV-ERG) pathway in wildtype and histone acetyltransferase (HAT) complex mutants.** RT-qPCR experiment was performed as described in Materials and Methods section to verify the mRNA transcript levels of key MEV-ERG pathway genes in wildtype and HAT complex mutants (n=3, Error bar represents ±SD).

**S2**

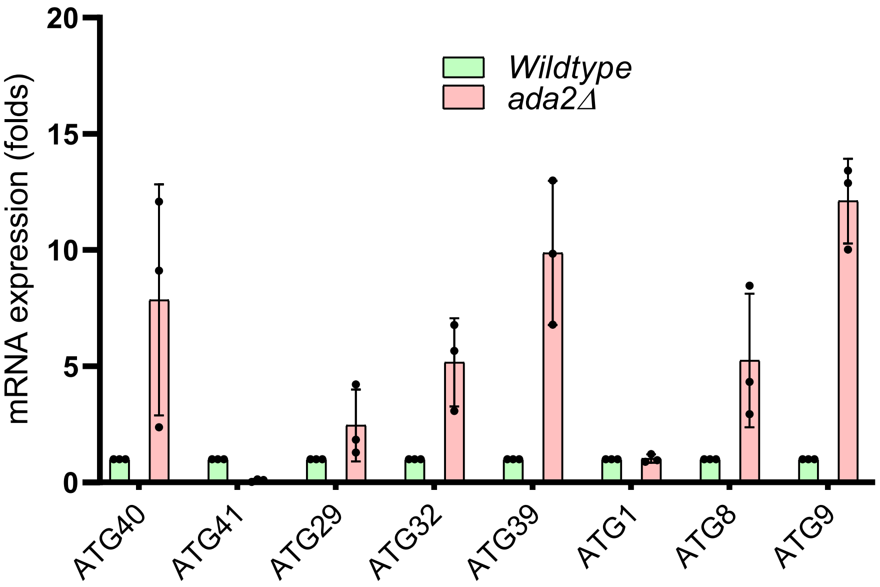


**Figure S2. Transcriptional analysis of autophagy pathway genes in wildtype versus *ada2Δ* cells.** RT-qPCR analysis was performed to investigate the expression levels of autophagy-related genes in wildtype (WT) and *ada2Δ* strains. Notably, *ATG40* and *ATG39* exhibited increased expression in *ada2Δ*, suggesting an upregulation of ER-phagy, which is essential for the selective degradation of endoplasmic reticulum. Furthermore, the elevated expression levels of *ATG8* and *ATG9* indicate a corresponding enhancement in general autophagy. These findings shows high autophagic response in *ada2Δ* strain. All the experiments were conducted in three biological replicates (n=3; Error bar ±SD).

**S3**

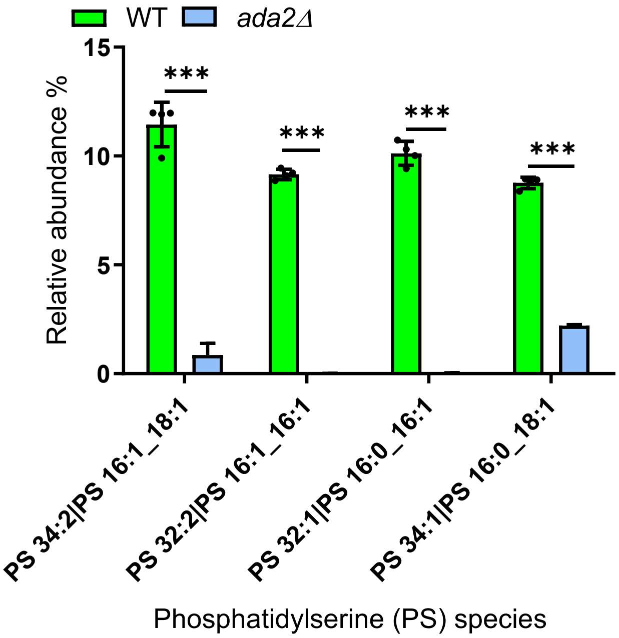


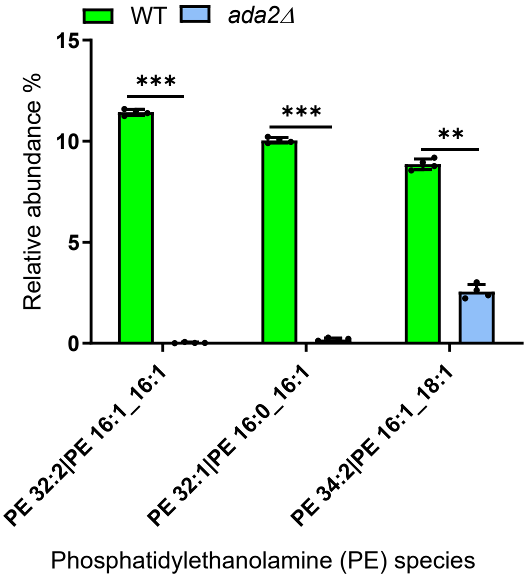


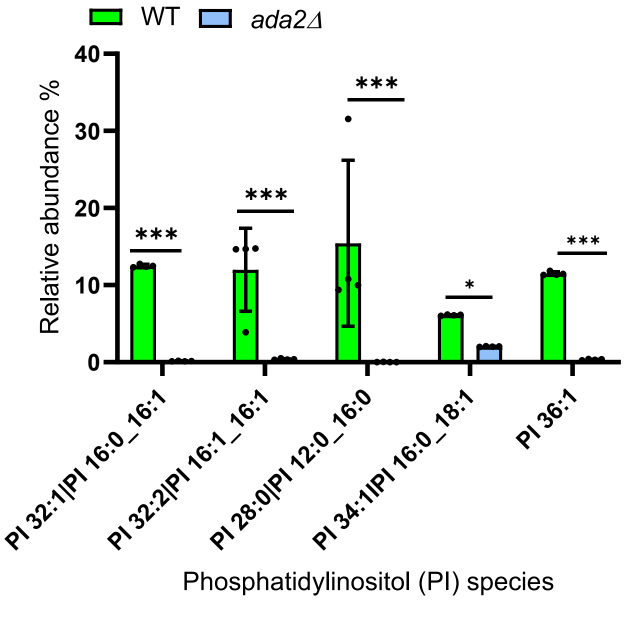


**Figure S3**. **Comparative lipidomic profiles of wildtype (*WT)* and *ada2Δ strains*.** Stationary phase grown equivalent $OD^{600}$ (optical density) of cells were collected and normalised to equal amount of protein crude lysate for lipid extraction. Lipidomics was carried out using a Vanquish UHPLC system coupled to an Orbitrap Exploris 240 mass spectrometer equipped with a HESI II source, operating in both positive and negative ionization modes. Lipid species were identified using the LIPID MAPS and LipidBlast databases. Quantified lipid classes include phospholipids (PS,PE and PI), expressed as relative percentages of total identified lipids. Individual molecular species within these classes were also resolved. Error bar indicate ±SD of four biological replicates, and statistical analysis was carried out using Student's t test, and the level of significance are shown as asterisks (n = 4; P value: ns = non significant, *=<0.1, **=<0.01; ***=<0.001).

**S4**

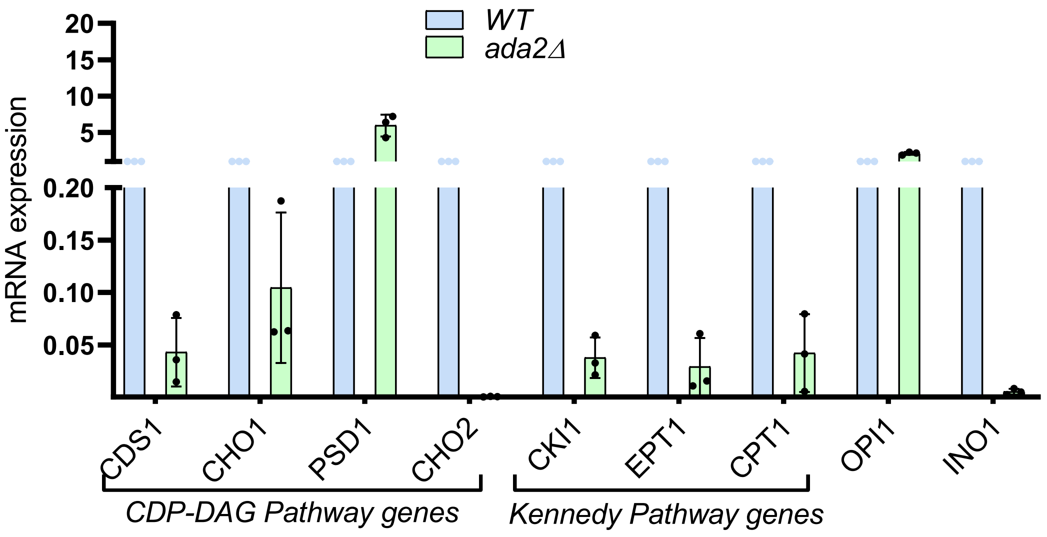


**Figure S4. Transcript and protein expression levels of key genes in the phospholipid biosynthesis pathway in wildtype and *ada2Δ* cells.** RT-qPCR analysis was conducted to evaluate the expression levels of genes involved in phospholipid metabolism, specifically those associated with the Cytidine Diphosphate Diacylglycerol (CDP-DAG) and Kennedy pathways. We also analyzed the gene expression of key inositol regulatory gene, *INO1*. Interestingly, the results indicated that the rate-limiting genes, *CDS1* and *INO1* are significantly downregulated in *ada2Δ* strain compared to wildtype (WT). This suggests that lipid metabolism is potentially altered in *ada2Δ.* All experiments were performed in three biological replicates (n=3; error bar ±SD).

S5

A)

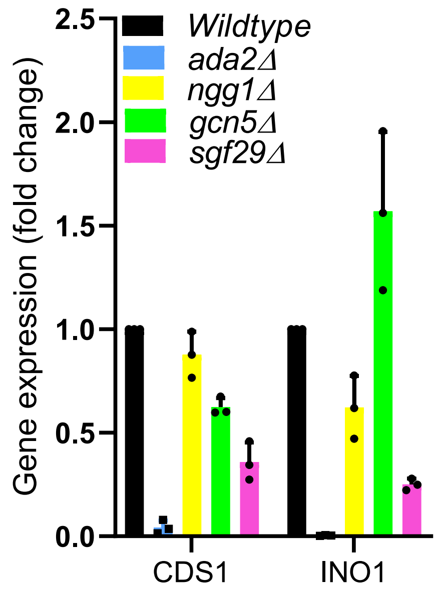


B)

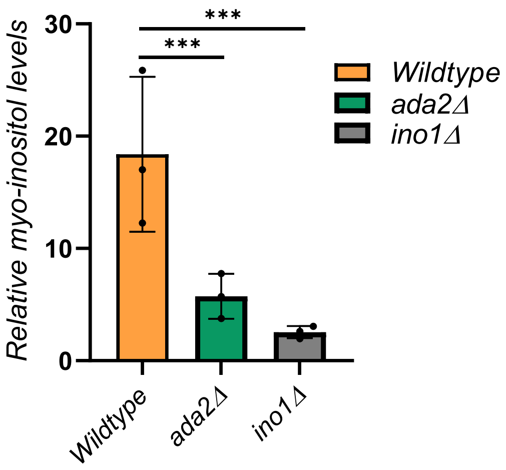


**Figure S5. Analysis of *CDS1* and *INO1* gene expression, and myo-inositol levels in wildtype and *ada2Δ*. A)** RT-qPCR data demonstrates mRNA transcript levels of *CDS1 and INO1* in *WT* and histone acetyltransferase complex mutants (n=3; error bar ±SD). **B)** Reduced endogenous inositol levels in *ada2Δ* and *ino1Δ* mutants measured by GC–MS. The *ada2Δ* and *ino1Δ* mutants show reduced inositol levels compared to wildtype (WT). Data represent three biological replicates, and Student's *t* test was performed to measure the level of significance (n=3; P values: ***=<0.001; error bar ±SD).

**S6**

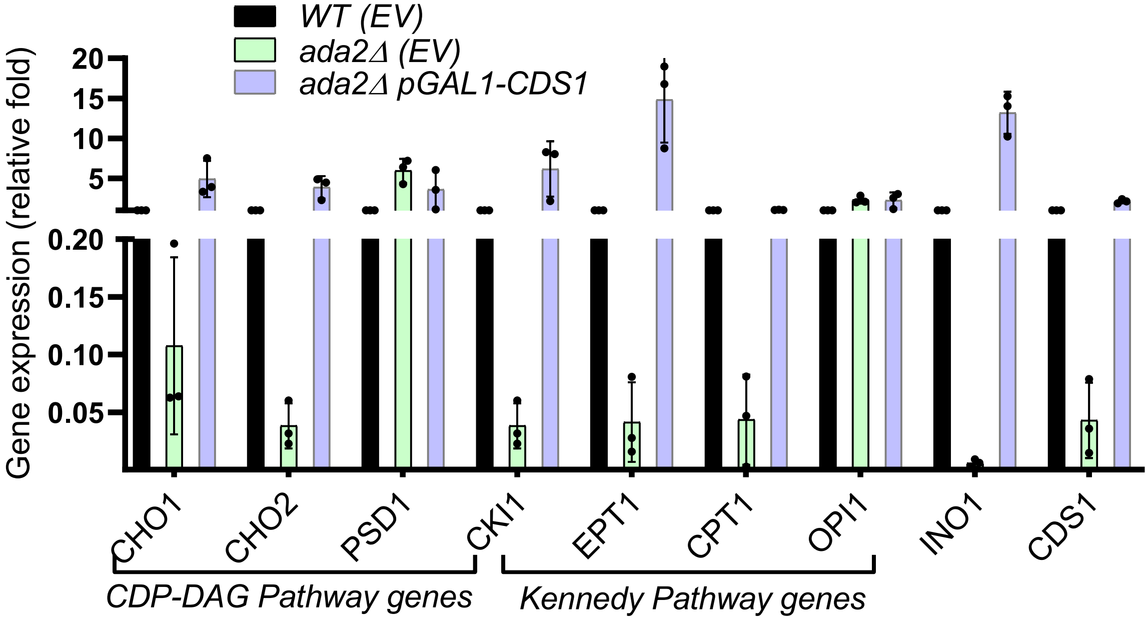


**Figure S6. mRNA transcript levels of phospholipid pathway genes in wildtype, *ada2Δ* and *ada2Δ pGAL1-CDS1* strains.** The RT-qPCR results show that *INO1* expression is significantly elevated in *CDS1* overexpressed *ada2Δ* as compared to *ada2Δ* empty vector (EV) control. Other pathway genes, such as *CHO2, EPT1, CHO1, PSD1, and CKI1*, also showed increased transcript levels, notably, *INO1* is significantly elevated. The experiment was performed in three biological replicates (n=3). WT (EV), Wildtype cells transformed with empty vector (pESC-LEU2d); ada2Δ *(EV), ada2Δ* transformed with empty vector *(*pESC-LEU2d). Error bar indicates ±SD.

**Fig.S7**

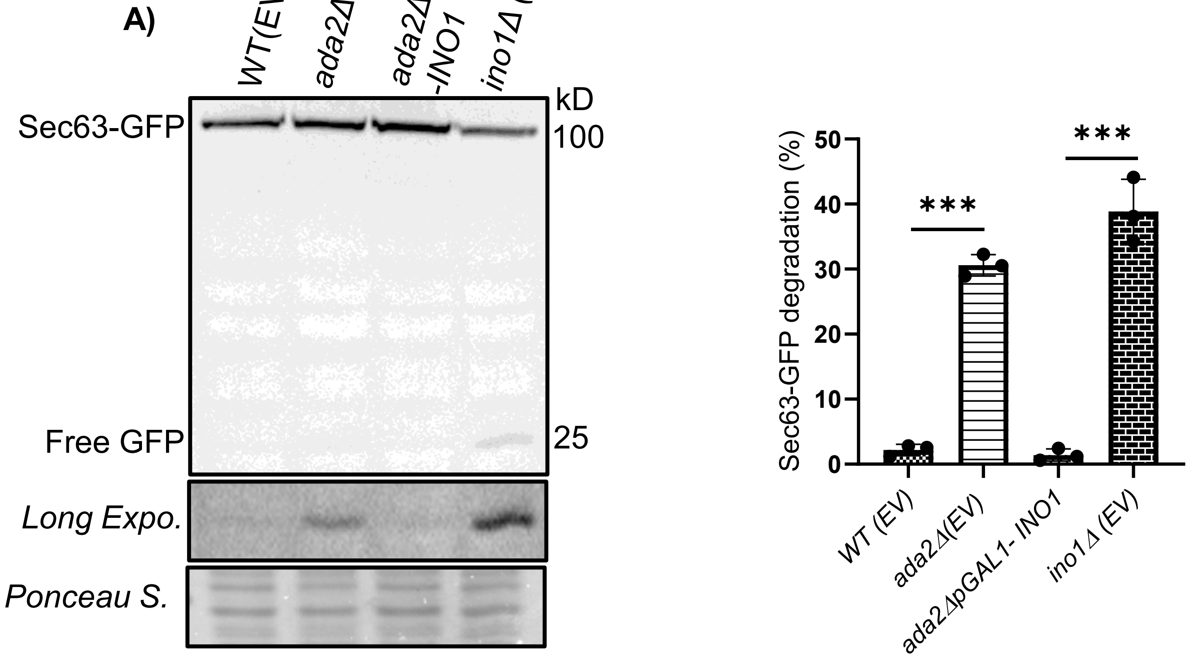

B)

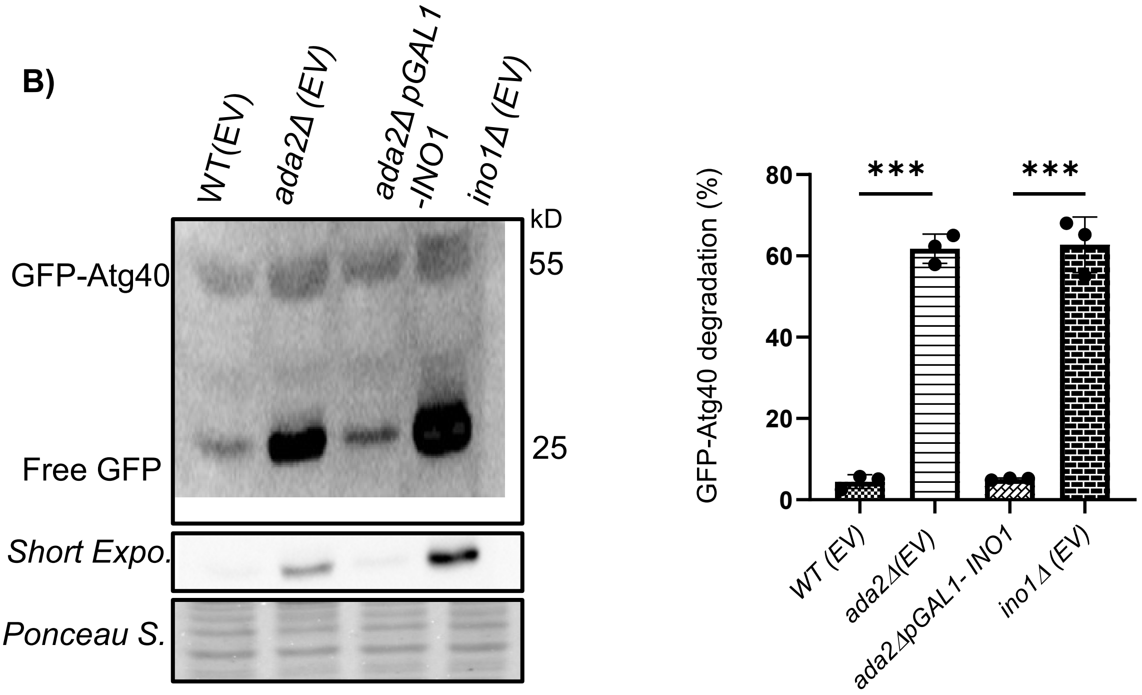

WT(EV)
ada2Δ (EV)
ada2Δ pGAL1 -INO1
ino1Δ (EV)
kD
GFP-Atg40
55
Free GFP
25
Short Expo.
Ponceau S.
GFP-Atg40 degradation (%)
80
60
40
20
0
***
***
WT (EV)
ada2Δ(EV)
ada2ΔpGAL1- INO1
ino1Δ (EV)


C)

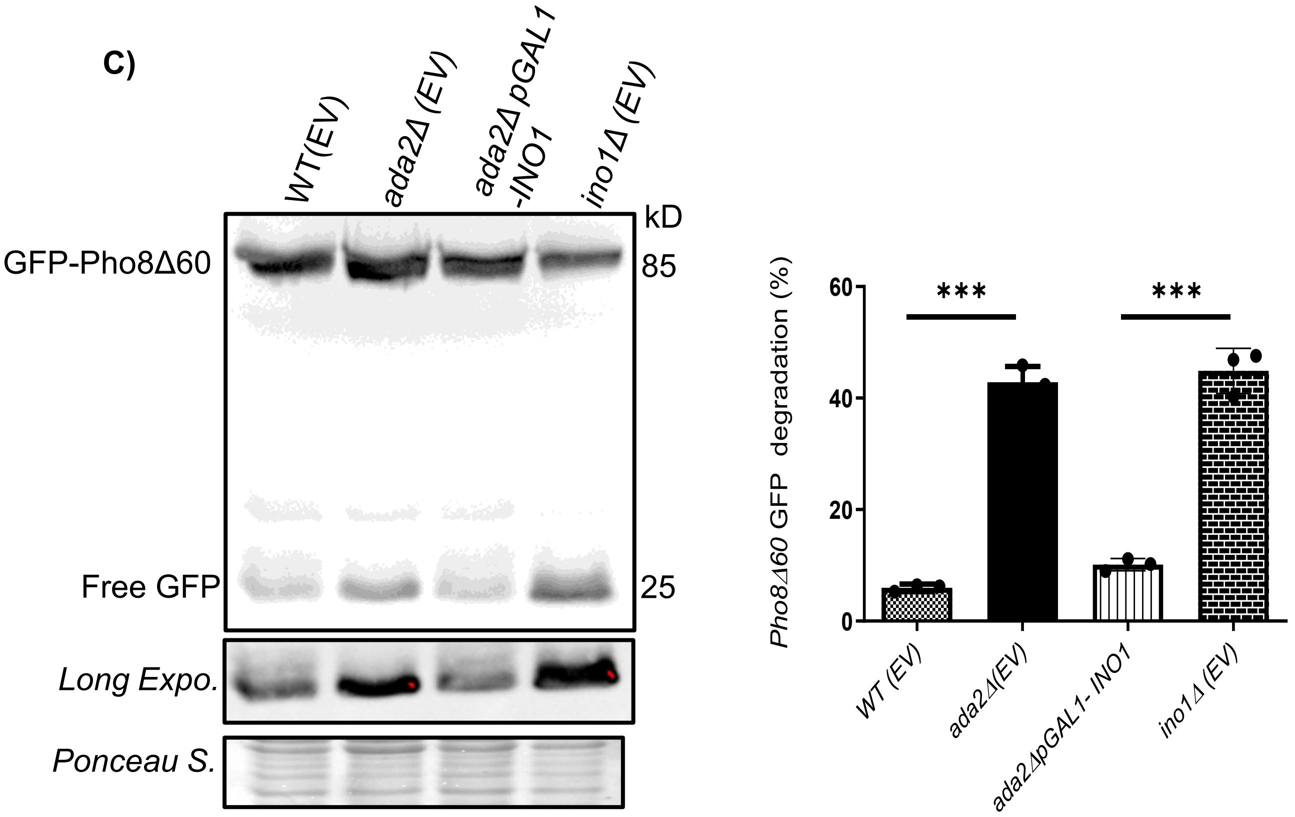

WT(EV)
ada2Δ (EV)
ada2Δ pGAL1 -INO1
ino1Δ (EV)
kD
GFP-Pho8Δ60
85
Free GFP
25
Long Expo.
Ponceau S.
Pho8Δ60 GFP degradation (%)
60
40
20
0
***
***
WT (EV)
ada2Δ(EV)
ada2ΔpGAL1- INO1
ino1Δ (EV)

D)

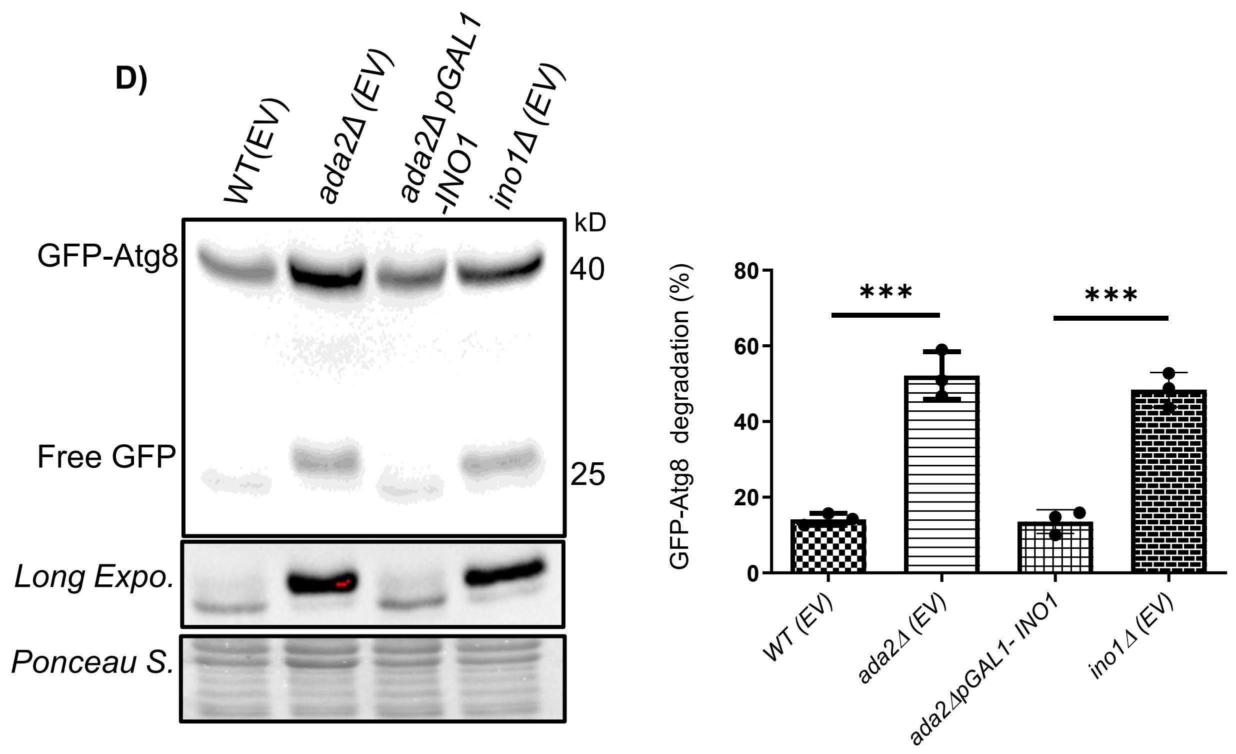


E)

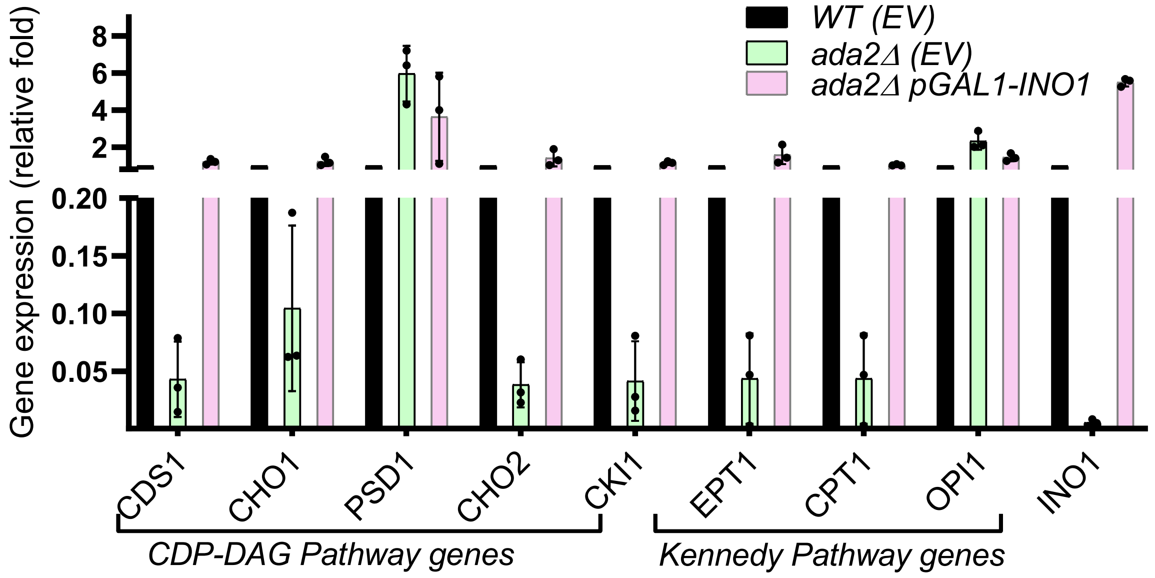


**Figure S7. Overexpression of *INO1* modulates ER-phagy, general autophagy, and phospholipid pathway gene expression in wildtype and *ada2Δ*. (A-D)** Sec63-GFP **(A)**, GFP-Atg40 **(B)**, Pho8Δ60-GFP **(C)** and GFP-Atg8 **(D)** processing assay showing free GFP release, an indicative of ER stress, general and macro autophagy levels (n=3, Student's *t* test, P values: ***=<0.001). **E)** The RT-qPCR data of *ada2Δ* pGAL1-*INO1* strain restored the expression levels of the phospholipid pathway genes, similar to WT empty vector control. The experiment was conducted with three biological replicates (n=3). WT (EV), Wildtype cells transformed with empty vector (Moclo-HIS3); *ada2Δ (EV), ada2Δ* transformed with empty vector. The error bar indicates ±SD.

**Fig.S8**

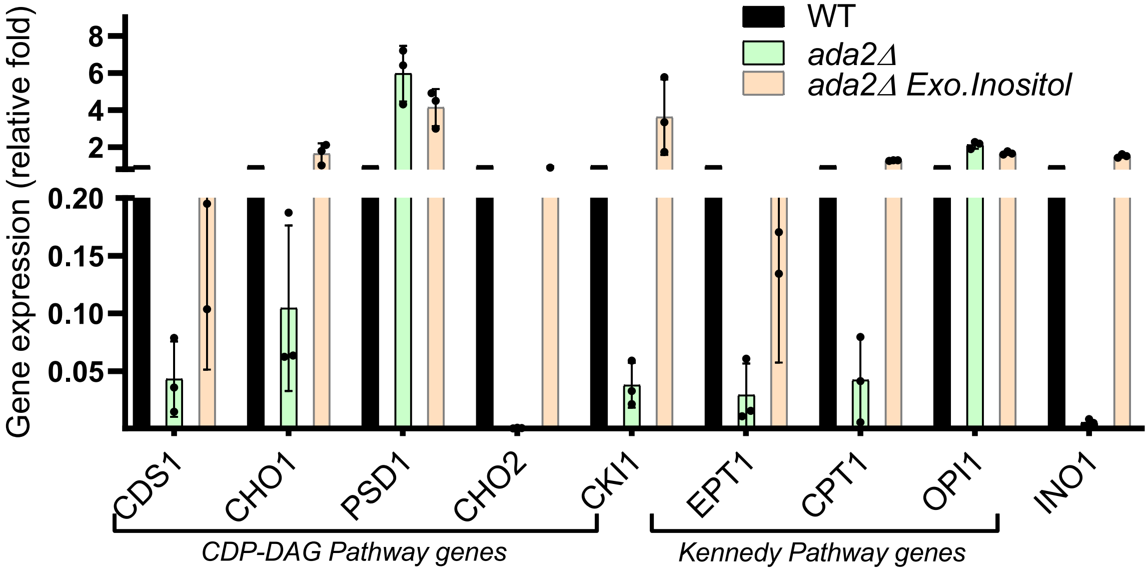


**Figure S8. Exogenous inositol supplementation restores transcript levels of phospholipid pathway genes in *ada2Δ*.** RT-qPCR analysis of *ada2Δ* supplemented with exogenous inositol at a concentration of 200 μM reveals a restoration of transcript levels for phospholipid pathway genes to those comparable with wildtype (WT). This indicates that the addition of exogenous inositol effectively rescues the expression of phospholipid pathway genes in *ada2Δ* strain. The experiments were conducted using Power SYBR Green master mix, and all analyses were performed in three biological replicates (n=3; Error bar, ±SD).

**S9**

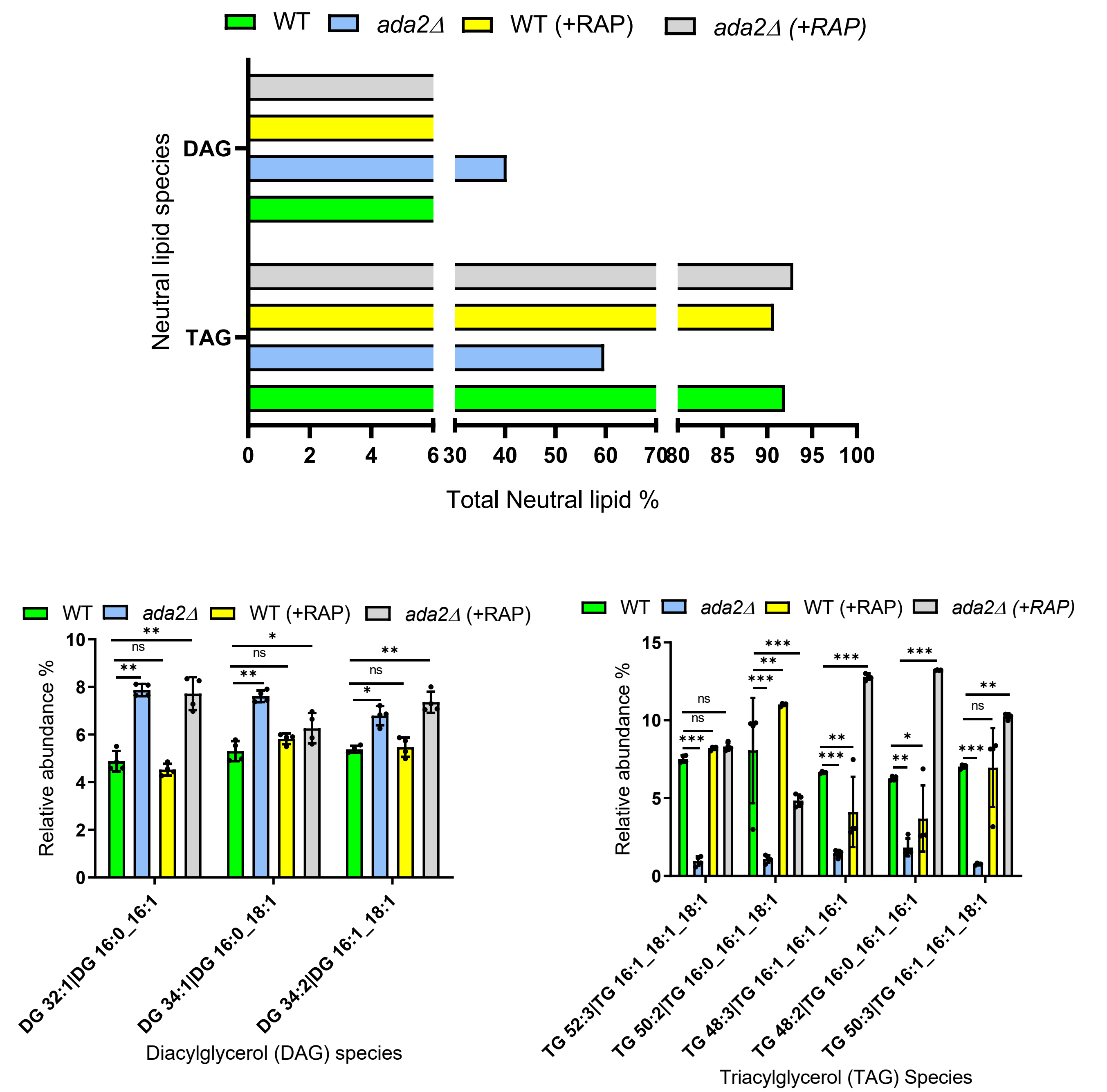


**Figure S9**. **Comparative neutral lipid profiling of wildtype and *ada2Δ* ± rapamycin.** Lipid extraction and analysis were carried out as described in S4 figure legend. TAG and DAG levels were shown as relative abundance in percentage of total neutral lipids. Individual molecular species within these classes were also resolved. Error bar indicate ±SD of four biological replicates (n = 4; Student's *t* test, P value: ns = non significant, *=<0.1, **=<0.01; ***=<0.001). Representation of species DAG(Diacylglycerol), TAG(Triacylglycerol). +RAP indicates rapamycin addition experiment.

**S10**

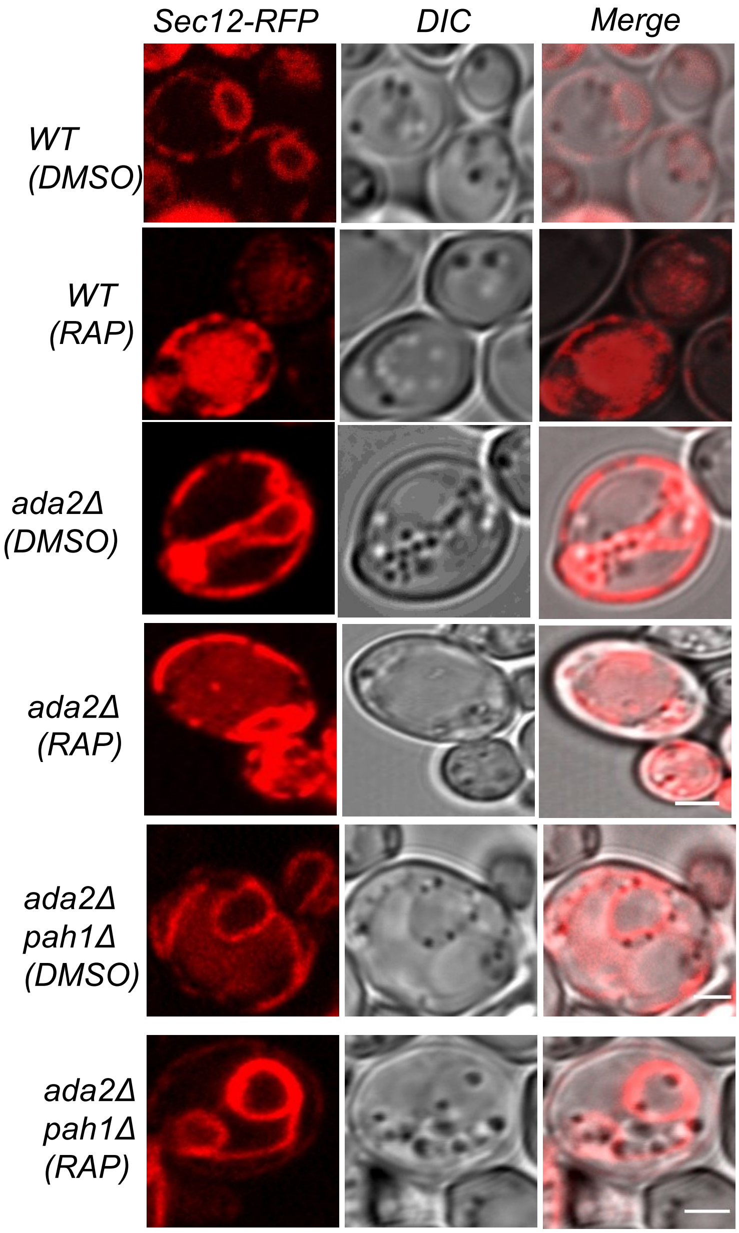


**Figure S10. Confocal microscopy analysis shows ER morphology in wildtype, *ada2Δ* and *ada2Δpah1Δ* cells with or without rapamycin treatment.** The Sec-12-RFP, serves as ER specific marker. The RFP signals were acquired using an excitation wavelength of 488 nm and an emission wavelength of 540 nm to visualize the ER dynamics. A total of seven Z-stacks images were collected for each sample and analyzed using ImageJ software (scale bar, 2 μm). The experiments were conducted in three biological replicates (n=3).

**S11**

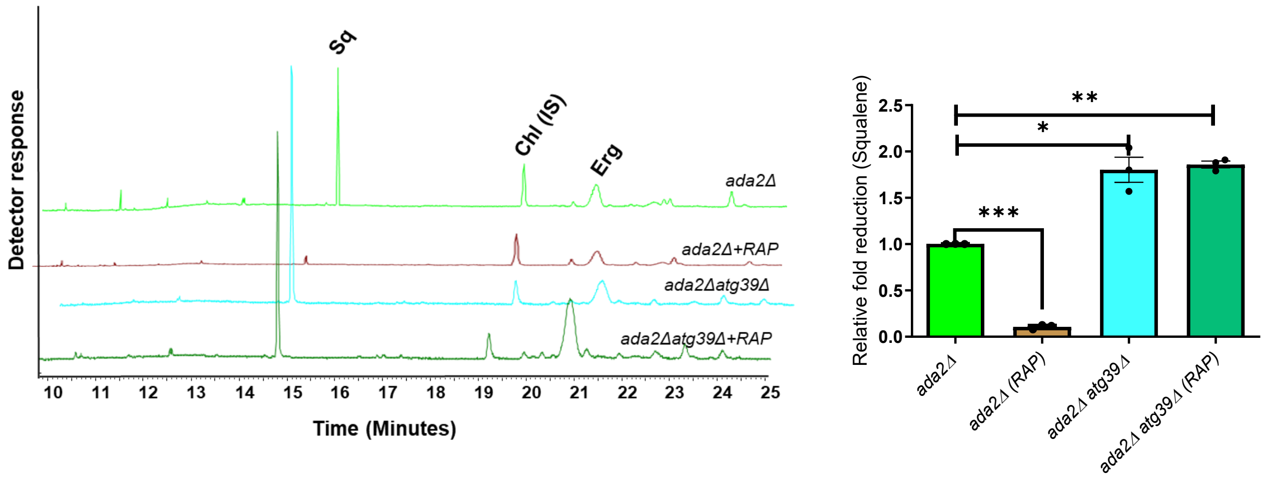


**Figure S11. Gas chromatogram shows no reduction in squalene (SQ) level in *ada2Δatg39Δ* and *ada2Δatg39Δ*-treated rapamycin strains.** The Atg39, nucleophagy receptor, elimination in *ada2Δ* increases the SQ accumulation, indicating lack of nER-phagy. The bar diagram represents the relative fold change in SQ accumulation in indicated strains. (n=3; error bar indicates ±SD; Student's *t* test; P values, ***p<0.0001,**p<0.001, *p<0.01).
