## Supplementary Tables for "Decoding the role of ADAptor2 (ADA2) of HAT complex in autophagy and phospholipid metabolism to maintain ER homeostasis and triterpene regulation"

| ***Supplementary Table 1. List of yeast strains use in this study*** | | |
| --- | --- | --- |
| ***Strain*** | ***Relevant genotype*** | ***Source*** |
| *BY4741* | *MATα his3Δ1 leu2Δ0 met15Δ0 ura3Δ0* | *Euroscarf* |
| *ada2Δ* | *MATα his3Δ1 leu2Δ0 met15Δ0 ura3Δ0, ada2::kanMX* | *Euroscarf* |
| *ada2Δpah1Δ* | *MATα his3Δ1 leu2Δ0 met15Δ0 ura3Δ0, ada2::kanMX,pah1::URA3* | *This study* |
| *ada2Δatg39Δ* | *MATα his3Δ1 leu2Δ0 met15Δ0 ura3Δ0, ada2::kanMX, atg39::HIS3* | *This study* |
| *BY4741 GFP-ATG8* | *BY4741, pRS423-ATG8-EGFP,HIS3* | *This study* |
| *BY4741 GFP -SEC63* | *BY4741, YCpLac33-sec63-GFP,URA3* | *This study* |
| *BY4741 GFP- Pho8Δ60* | *BY4741 pRS416-P_GPD_-GFP-Pho8Δ60-T_CYC1_ ,URA3* | *This study* |
| *BY4741 RFP-SEC12* | *BY4741, pRS316-SEC12-mcherry RFP,URA3* | *This study* |
| *BY4741 ATG40-EGFP* | *BY4741, pRS316-ATG40-EGFP, URA3* | *This study* |
| *BY4741 GFP-OPI1* | *BY4741, pPM494-Opi1 GFP, LEU2* | *This study* |
| *ada2Δ ATG8-GFP* | *BY4741, ada2::kanMX, pRS423-ATG8-EGFP, HIS* | *This study* |
| *ada2Δ GFP-SEC63* | *BY4741, ada2::kanMX, YCpLac33-sec63-GFP,URA3* | *This study* |
| *ada2Δ GFP- Pho8Δ60* | *BY4741, ada2::kanMX, pRS416-P_GPD_-GFP-Pho8Δ60-T_CYC1_ ,URA3* | *This study* |
| *ada2Δ RFP-SEC12* | *BY4741, ada2::kanMX, pRS316-SEC12-mcherry RFP,URA3* | *This study* |
| *ada2Δ GFP-ATG40* | *BY4741, ada2::kanMX, pRS316-ATG40-EGFP, URA3* | *This study* |
| *ada2Δ GFP-OPI1* | *BY4741, ada2::kanMX, pPM494-Opi1 GFP, LEU2* | *This study* |
| *ada2Δ pGAL1-INO1* | *BY4741, ada2::kanMX, MoClo pGAL1-INO1 HIS* | *This study* |
| *ada2Δ (empty vector)* | *BY4741, ada2::kanMX, MoClo HIS* | *This study* |
| *ada2Δ pGAL1-INO1 GFP-SEC63* | *BY4741, ada2::kanMX, MoClo pGAL1-INO1 HIS,YCpLac33-sec63-GFP URA3* | *This study* |
| *ada2Δ pGAL1-INO1 GFP- Pho8Δ60* | *BY4741, ada2::kanMX, MoClo pGAL1-INO1 HIS, pRS416-P_GPD_-GFP-Pho8Δ60-T_CYC1_ URA3* | *This study* |
| *ada2Δ pGAL1-INO1 RFP-SEC12* | *BY4741, ada2::kanMX, MoClo pGAL1-INO1 HIS, pRS316-SEC12-mcherry RFP URA3* | *This study* |
| *ada2Δ pGAL1-INO1 GFP-ATG40* | *BY4741, ada2::kanMX, MoClo pGAL1-INO1 HIS, ATG40-GFP URA3* | *This study* |
| *ada2Δ pGAL1-INO1 GFP-OPI1* | *BY4741, ada2::kanMX, MoClo pGAL1-INO1 HIS, pPM494-Opi1 GFP LEU2* | *This study* |
| *ada2Δ pGAL1-CDS1* | *BY4741, ada2::kanMX, pESCLeu2d- pGAL1-CDS1 LEU* | *This study* |
| *ada2Δ (empty vector)* | *BY4741, ada2::kanMX, pESCLeu2d LEU* | *This study* |
| *ada2Δ pGAL1-CDS1 GFP-SEC63* | *BY4741, ada2::kanMX, pESCLeu2d- pGAL1-CDS1 LEU,YCpLac33-sec63-GFP URA3* | *This study* |
| *ada2Δ pGAL1-CDS1 GFP- Pho8Δ60* | *BY4741, ada2::kanMX, pESCLeu2d- pGAL1-CDS1 LEU, pRS416-P_GPD_-GFP-Pho8Δ60-T_CYC1_ URA3* | *This study* |
| *ada2Δ pGAL1-CDS1 RFP-SEC12* | *BY4741, ada2::kanMX, pESCLeu2d- pGAL1-CDS1 LEU, pRS316-SEC12-mcherry RFP URA3* | *This study* |
| *ada2Δ pGAL1-CDS1 GFP-ATG40* | *BY4741, ada2::kanMX, pESCLeu2d- pGAL1-CDS1,LEU pRS316-ATG40-EGFP,URA3* | *This study* |
| *ada2Δ pGAL1-CDS1 GFP-ATG8* | *BY4741, ada2::kanMX, pESCLeu2d- pGAL1-CDS1 LEU, pRS423-ATG8-EGFP, HIS* | *This study* |
| *ada2Δ pGAL1-ADA2* | *BY4741, ada2::kanMX, pESCLeu2d- pGAL1-ADA2 LEU* | *This study* |
| *ada2Δ (empty vector)* | *BY4741, ada2::kanMX, pESCLeu2d LEU* | *This study* |
| *ada2Δ pGAL1- ADA2 GFP-SEC63* | *BY4741, ada2::kanMX, pESCLeu2d- pGAL1- ADA2 LEU,YCpLac33-sec63-GFP URA3* | *This study* |
| *ada2Δ pGAL1- ADA2 GFP- Pho8Δ60* | *BY4741, ada2Δ::kanMX, pESCLeu2d- pGAL1- ADA2 LEU, pRS416-P_GPD_-GFP-Pho8Δ60-T_CYC1_ URA3* | *This study* |
| *ada2Δ pGAL1- ADA2 RFP-SEC12* | *BY4741, ada2::kanMX, pESCLeu2d- pGAL1- ADA2 LEU, pRS316-SEC12-mcherry RFP URA3* | *This study* |
| *ada2Δ pGAL1- ADA2 GFP-ATG40* | *BY4741, ada2::kanMX, pESCLeu2d- pGAL1- ADA2 LEU, pRS316-ATG40-EGFP,URA3* | *This study* |
| *ada2Δ pGAL1- ADA2 GFP-ATG8* | *BY4741, ada2::kanMX, pESCLeu2d- pGAL1- ADA2 LEU, pRS423-ATG8-EGFP, HIS* | *This study* |
| *ada2Δ pah1Δ GFP ATG8* | *BY4741, ada2::kanMX,pah1::URA3 pRS423-ATG8-EGFP, HIS* | *This study* |
| *ada2Δ pah1Δ GFP OPI1* | *BY4741, ada2::kanMX,pah1::URA3, pPM494-Opi1 GFP LEU2* | *This study* |
| *ada2Δ GFP-OPI1 RFP-PUS1* | *BY4741, ada2::kanMX, pPM494-Opi1 GFP LEU2,pRS426-SPO20-RFP URA* | *This study* |
| *BY4741 GFP-OPI1 RFP-PUS1* | *BY4741, pPM494-Opi1 GFP LEU2,pRS426-SPO20-RFP URA* | *This study* |
| *ada2Δ pah1Δ pUL9* | *BY4741, ada2::kanMX,pah1::URA::LEU* | *This study* |
| *ada2Δ pah1Δ pUL9 Sec12-RFP* | *BY4741, ada2::kanMX,pah1::LEU pRS316-SEC12-mcherry RFP URA3* | *This study* |
| *ada2Δpah1Δ pUL9 Sec63-GFP* | *BY4741, ada2::kanMX,pah1::LEU, YCpLac33-sec63-GFP URA3* | *This study* |
| *ada2Δ pah1Δ DGK1 GFP* | *BY4741, ada2::kanMX,pah1::URA3 YCplac111-DGK1-GFP* | *This study* |
| *spo7Δ* | *MATα his3Δ1 leu2Δ0 met15Δ0 ura3Δ0, ada2::kanMX* | *Euroscarf* |
| *nem1Δ* | *MATα his3Δ1 leu2Δ0 met15Δ0 ura3Δ0, ada2::kanMX* | *Euroscarf* |
| *BY-Sch9 Cterm-6HA* | *BY4741, Sch9::NAT* | *This study* |
| *ada2Δ-Sch9 Cterm.-6HA* | *BY4741, ada2::kanMX, Sch9::NAT* | *This study* |
| *BY-CKI1 Cterm-GFP* | *BY4741, CKI1::HYG* | *This Study* |
| *ada2Δ-CKI1 Cterm-GFP* | *BY4741, ada2::kanMX, CKI1::HYG* | *This Study* |
| *BY-EKI1 Cterm-GFP* | *BY4741, EKI1::HYG* | *This Study* |
| *ada2Δ-EKI1 Cterm-GFP* | *BY4741, ada2::kanMX, EKI1::HYG* | *This Study* |
| *BY-OPI3 Cterm-GFP* | *BY4741, OPI3::HYG* | *This Study* |
| *ada2Δ-OPI3 Cterm.-GFP* | *BY4741, ada2::kanMX, OPI3::HYG* | *This Study* |
| *BY-EPT1 Cterm.-GFP* | *BY4741, EPT1::HYG* | *This Study* |
| *ada2Δ-EPT1 Cterm.-GFP* | *BY4741, ada2::kanMX, EPT1::HYG* | *This Study* |

***Supplementary Table 2. List of plasmids used in study***

| ***Plasmid*** | ***Genotype*** | ***Source*** |
| --- | --- | --- |
| *SEC12-mRFP* | *pRS316 SEC12 mCherry-RFP* | *Autophagy lab, JNCASR* |
| *GFP-Pho8Δ60* | *pRS416-P_GPD_-GFP-Pho8Δ60-T_CYC1,_ URA3* | *CW Wang’s lab, Taipei, Taiwan (67)* |
| *GFP-OPI1* | *pPM494-Opi1 GFP* | *Dr. J. Pedro Fernandez Murray (38)* |
| *GFP-SPO20* | *pRS426 SPO20 GFP/RFP* | *Dr Symeon siniossoglouas* |
| *RFP-PUS1* | *pRS426 PUS1 RFP* | *This study* |
| *GFP-ATG40* | *pRS316-ATG40-EGFP* | *Dr. Hitoshi Nakatogawa (68)* |
| *GFP-ATG8* | *pRS423-ATG8-EGFP* | *Prof. Ravi Manjithaya, JNCASR, Autophagy Lab* |
| *SEC63-GFP* | *YCpLac33-sec63-GFP* | *Dr Symeon siniossoglouas (69)* |
| *pGAL1-INO1* | *MoClo pGAL1-INO1 HIS* | *This Study* |
| *pGAL1-CDS1* | *pESC-pGAL1-CDS1-Leu2d* | *This Study* |
| *pGAL1-ADA2* | *pESC-pGAL1-ADA2-Leu2d* | *This Study* |
| *pUL9* | *LEU2-kanR* | *Addgene (71)* |
| *DGK1-GFP* | *YCplac111-DGK1-GFP* | *Our Laboratory stock* |
| *Sch9-6HA* | *pFA6a-natMX6* | *Dr. Sunil Laxman, In-STEM NCBS* |
| *GFP-TAG* | *pFA6a-hphMX6-GFP* | *Dr. Sunil Laxman, In-STEM NCBS* |

| ***Supplementary Table 3. List of cloning and knockout primers used in this study.*** | | |
| --- | --- | --- |
| ***Primer*** |  | ***Sequence 5’‐>3’*** |
| *ADA2_GAL1* |  | *ATAATGGATCCATGTCAAACAAGTTTCACTGTGACGTT*  *ATAATCTCGAGTTACATCCAATTCTGGCTCTGGAAAAAA* |
| *pYM ATG39 KO* |  | *CAGTGACGATAATAGAGACTAGTAAAACAGTCGAGTTGTCGGACCTAAAATGCGTACGCTGCAGGTCGAC*  *CGTTTTTTTTTTCTTTTGTTAATTTCATTCTTCATGCTGGGTTTTGGATGATCTAATCGATGAATTCGAGCTCG* |
| *Pah1 KO- URA* |  | *TACAGGGAAGAAATTACTGAAGATAGACACATCGGTCGATTATGCCAGATTGTACTGAGAGTGCACCA*  *GTATGGATCGTTATAAATAATATTCGGCTACAAGAATCTTTACCTGATGCGGTATTTTCTCCTTACG* |
| *PUS1_RFP* |  | *ATAATTCTAGAATGTCTGAAGAGAATTTGAGGCCTG*  *ATAATCTCGAGCTAATTAGCTGCCGCTTCCGG* |
| *INO1_GAL1* |  | *ATAGAGGATCCAACACAATGACAGAAGATAATATTGCTCCAA*  *ATAACTCGAGTTACAACAATCTCTCTTCGAATCTT* |
| *CDS1_GAL1* |  | *ATAGAGGATCCAACACAATGTCTGACAACCCTGAGATGAAACC*  *ATAAAAGCTTTCAAGAGTGATTGGTCAATGATTTCTTGG* |
| *Sch9 TagF2*; *Sch9 R1*; *HAC1 SP F* *HAC1 SP R* |  | *ACACATGGATGACGAATTTGTCAGTGGAAGATTCGAAATACGGATCCCCGGGTTAATTAA*; *AAGGAAAAGAAGAGGAAGGGCAAGAGGAGCGATTGAGAAAGAATTCGAGCTCGTTTAAAC* *ACGACGCTTTTGTTGCTTCT* *TCTTCGGTTGAAGTAGCACAC* |
| *OPI3 C' tag pFA6 F*  *OPI3 C' tag pFA6 R* |  | *CATGATCTACGCTAACCGTGATAAGGCCAAAAAGAATATGcggatccccgggttaattaa*  *GGCTTCTAACATTATAGAATATATAGAAATAGAGCACgaattcgagctcgtttaaac* |

| ***Supplementary Table 4. List of RT qPCR primers used in this study.*** | | |
| --- | --- | --- |
| ***Primer*** |  | ***Sequence 5’‐>3’*** |
| *ATG39 RT* | *Fwd*  *Rev* | *CTGGAGTTCCCACTATTTCAGAG*  *GTTTCGACCCCTCCATACTTG* |
| *ATG40 RT* | *Fwd*  *Rev* | *CCTCAGTTGCCATTCCTTTG*  *CAGCACTTGCCTTTGAGAAG* |
| *ATG41 RT* | *Fwd*  *Rev* | *AACTTTTCCGCTATCGTACCC*  *TGCAATCGTCTTCAGTACTCG* |
| *INO1 RT* | *Fwd*  *Rev* | *TCAAGATGAGAGAGCCAATAACT*  *GTCATTAACACCAGGAGATACTTCTA* |
| *OPI1 RT* | *Fwd*  *Rev* | *GCACAGTCAAGAAAGTCTACTC*  *ACAGTTGGCATAATGAAACGA* |
| *OPI3 RT* | *Fwd*  *Rev* | *ATTGCGTGAACAGCCTAC*  *ATCCATCAGGATGCCGA* |
| *ACT1 RT* | *Fwd*  *Rev* | *CTGAAAGAGAAATTGTCCGTG*  *GGGCTCTGAATCTTTCGTT* |
| *CDS1 RT* | *Fwd*  *Rev* | *ATCATAAGTTTATCTGCTATTGTCTCT*  *GATGATCAAATGAGCCTGAAATAC* |
| *EKI1 RT* | *Fwd*  *Rev* | *ATTTAGGACCCAAACTAGAAGG*  *AATGGAACAGTACAATGCAAC* |
| *CHO1 RT* | *Fwd*  *Rev* | *ATATGGCAGATTACATTACTATGCT*  *CGTCAAGGAAATCGAAACAC* |
| *CHO2 RT* | *Fwd*  *Rev* | *ATTCTTGTGGCTTTACAAACAT*  *CTGGGTGATTCAAATATCTGTAG* |
| *EPT1 RT* | *Fwd*  *Rev* | *CCGTATCAATCAATCAGGACC*  *ATTCGTCAGCATCGCAA* |
| *CKI1 RT* | *Fwd*  *Rev* | *TGCCCAATACGGCAATTTAC*  *GGCGGGTTGATTATATCATCT* |
| *CPT1 RT* | *Fwd*  *Rev* | *TTTGCGATATTGTGCAGCTT*  *AGATGGTTTGAGGTCCGTA* |
| *PSD1 RT* | *Fwd*  *Rev* | *AGATGGAAGATCCTGATTTGACA*  *GATTTCGCCAGTTTCAGAGT* |
| *ATG1 RT* | *Fwd*  *Rev* | *ATCTAAGATGGCCGCACATATG*  *AGGGTAGTCACCATAGGCATTC* |
| *ATG8 RT* | *Fwd*  *Rev* | *GAAGGCCATCTTCATTTTTGTC*  *TTCTCCTGAGTAAGTGACATAC* |
| *ATG9 RT* | *Fwd*  *Rev* | *CGTACTAACAGAGTCTTTCCTTG*  *CTAAGACACCACCCTTATTGAG* |
| *ATG29 RT* | *Fwd*  *Rev* | *ATGAGGCGTTACAACATTTGC*  *TCGTCATCTGAACTACCGCAC* |
| *ATG32 RT* | *Fwd*  *Rev* | *GGGCAAAATGAATACTTTTGTCTTGCATGC*  *CCCAGTGCCAAAATCCGATTAGATTCATC* |
| *DPRT21 (KAR2)*  *DPRT22 (KAR2)* | *Fwd*  *Rev* | *TCTGAAGGTGTCTGCCACAG*  *TTAGTGATGGTGATAGATTCGGATT* |
| *DPRT23 (IRE1)*  *DPRT24 (IRE1)* | *Fwd*  *Rev* | *AGTCAGAATTTTCCATCTTTAGTCG*  *CGGTCACTGGAAGCGTATCT* |
| *DPRT25 (HAC1)*  *DPRT26 (HAC1)* | *Fwd*  *Rev* | *GAAACAGTCTACCCTTTGACAAT*  *CTGCGCTTCTGGATTACG* |
| *Rpl30 qRT*  *Rpl30 qRT* | *Fwd*  *Rev* | *ACTACTTCCAAGGTGGTAAC*  *AGCCAAGGTGGTCAAGATATC* |
| *Rpl16A qRT (F)*  *Rpl16A qRT (R)* | *Fwd*  *Rev* | *TTGGTTACTCCTCAAAGATTGC*  *CAGACAATCTCTTAGCCAACAA* |
| *Rpl32 qRT (F)*; *Rpl32 qRT (R)* | *Fwd*  *Rev* | *CACCAAGACTTACGCCGCT*  *CCCTTTGGGTTGGTGACCT* |
